# White matter microstructure and macrostructure brain charts across the human lifespan

**DOI:** 10.1101/2025.05.08.652953

**Authors:** Michael E. Kim, Chenyu Gao, Karthik Ramadass, Nancy R. Newlin, Praitayini Kanakaraj, Sam Bogdanov, Gaurav Rudravaram, Derek Archer, Timothy J. Hohman, Angela L. Jefferson, Victoria L. Morgan, Alexandra Roche, Dario J. Englot, Susan M. Resnick, Lori L. Beason Held, Laurie Cutting, Laura A. Barquero, Micah A. D’archangel, Tin Q. Nguyen, Kathryn L. Humphreys, Yanbin Niu, Sophia Vinci-Booher, Carissa J. Cascio, The HABS-HD Study Team, Alzheimer’s Disease Neuroimaging Initiative, The BIOCARD Study Team, L. Taylor Davis, Zhiyuan Li, Simon N. Vandekar, Panpan Zhang, John C. Gore, Bennett A. Landman, Kurt G. Schilling

## Abstract

Normative reference charts are widely used in healthcare, especially for assessing the development of individuals by benchmarking anatomic and physiological features against population trajectories across the lifespan. Recent work has extended this concept to gray matter morphology in the brain, but no such reference framework currently exists for white matter (WM) even though WM constitutes the essential substrate for neuronal communication and large-scale network integration. Here, we present the first comprehensive WM brain charts, which describe how microstructural and macrostructural features of WM evolve across the lifespan, by leveraging over 35,120 diffusion MRI scans from 50 harmonized studies. Using generalized additive models for location, scale, and shape (GAMLSS), we estimate age- and sex-stratified trajectories for 72 individual white matter pathways, quantifying both tract-specific microstructural and morphometric features. We demonstrate that these WM brain charts enable four important applications: (1) defining normative trajectories of WM maturation and decline across distinct pathways, (2) identifying previously uncharacterized developmental milestones and spatial gradients of tract maturation, (3) detecting individualized deviations from normative patterns with clinical relevance across multiple neurological disorders, and (4) facilitating standardized, cross-study centile scoring of new datasets. By establishing a unified, interpretable reference framework for WM structure, these brain charts provide a foundational metric for research and clinical neuroscience. The accompanying open-access trajectories, centile scoring tools, and harmonization methods facilitate precise mapping of WM development, aging, and pathology across diverse populations. We release the brain charts and provide an out-of-sample alignment process as a Docker image: https://zenodo.org/records/17561821.

## Introduction

We report our development of *White Matter Brain Charts*, the first comprehensive, large-scale normative framework that documents changes in white matter (WM) microstructure and macrostructure across the human lifespan. By integrating and analyzing diffusion MRI (dMRI) data from over 35,120 individual brains (composed of 4,253,251 imaging volumes), across 50 different population studies, we have systematically characterized axonal density and dispersion, as well as tract volume, length, and shape, in functionally relevant WM pathways, from early life to mature adulthood. These *White Matter Brain Charts* may be used in several important applications including: (1) defining normative trajectories of microstructural and macrostructural features of specific white matter pathways, (2) revealing previously uncharacterized developmental milestones, refining our understanding of critical periods of WM growth and decline, (3) enabling sensitive detection of deviations from normative pathways, with implications for identifying early markers of neurodevelopmental and neurodegenerative disorders, and (4) enabling standardized centile scoring of new datasets to quantify WM alterations in clinical populations.

The human brain is an intricately organized network, with white matter (WM) forming the backbone of large-scale neural communication [1,2]. While much focus in neuroscience has been placed on gray matter (GM) morphology and its association with cognition, behavior, and disease, WM comprises nearly half of total brain volume and serves as the fundamental conduit for information transfer between cortical and subcortical regions. Disruptions in WM integrity are implicated across a spectrum of neurodevelopmental, neuropsychiatric, and neurodegenerative disorders [3,4]. However, despite these crucial roles of WM in brain functions, no standardized normative reference data have previously been derived for characterizing individual variability, developmental milestones, or pathological deviations in WM structure. These may now be derived using appropriate image analysis tools by leveraging the large array of imaging data sets that are publicly available.

The concept of normative brain charts - analogous to pediatric growth charts - has recently been introduced [5] to benchmark some neuroanatomical trajectories across the lifespan, including GM features such as cortical thickness and volume [5–8]. These efforts have elucidated fundamental principles of neurodevelopment and aging, enabling robust comparisons across individuals and clinical populations. However, in the context of WM, these previous efforts were predominantly limited to global volumetric analyses and did not provide the tract-specific, pathway-level phenotypes needed to localize deviations and to relate white-matter architecture to circuit function and dysfunction. This constraint is largely due to methodological challenges of dMRI, the only non-invasive imaging modality that allows for studying directionality and organization of WM pathways in the human brain. These challenges include complexities in fiber tractography, inter-site variability, and historically limited large-scale, harmonized datasets [9]. Given the increasing recognition of WM’s role in both normal brain functions and disease, establishing a rigorous, data-driven, normative reference for WM is a pressing scientific need. Just as with pediatric growth charts, defining this standard of healthy maturation is the necessary prerequisite for identifying and quantifying atypical development and decline.[10,11]

Diffusion MRI provides a rich array of information on WM structure including microstructural indices that reflect axonal density and dispersion [12], as well as macrostructural properties such as WM tract volume, length, and shape [13]. By integrating methodological advances in tract segmentation, data harmonization, and statistical modeling that charts the full population distribution (e.g., location, scale, and skew) of inter-individual variability, we have established age- and sex-stratified trajectories of WM organization from early childhood through advanced aging. By establishing standardized benchmarks for WM maturation and degeneration we provide the foundational data to support discovery in both fundamental neuroscience and for clinical translation, facilitating individualized assessments of brain health. These reference charts, along with openly accessible processing tools and harmonization pipelines, create a robust foundation for future research, large-scale collaborations, and personalized approaches to studying WM alterations in health and disease.

## Results

### Mapping Normative White Matter Growth

We included and analyzed dMRI data from a cohort of 35,120 individuals spanning 50 population studies and including 4,253,251 imaging volumes, representing typical development and aging with no known neurological or psychiatric conditions. We modeled both global and tract-specific WM phenotypes - including 72 anatomically defined pathways - using generalized additive models for location, scale, and shape (GAMLSS) [14], a flexible statistical framework endorsed by the World Health Organization for modeling non-linear biological trajectories [15]. GAMLSS enabled simultaneous estimation of age-dependent changes in location, scaling, and skewness, while accounting for study-level batch effects. Microstructural (fractional anisotropy - FA, mean diffusivity - MD, axial diffusivity - AD, radial diffusivity - RD) and macrostructural (tract volume, length, surface area) features were modeled across the full lifespan (**Figure 1; Supplemental Table SB.T1**; demographic information in **Supplemental Table SB.T2**).

**Figure 1.**
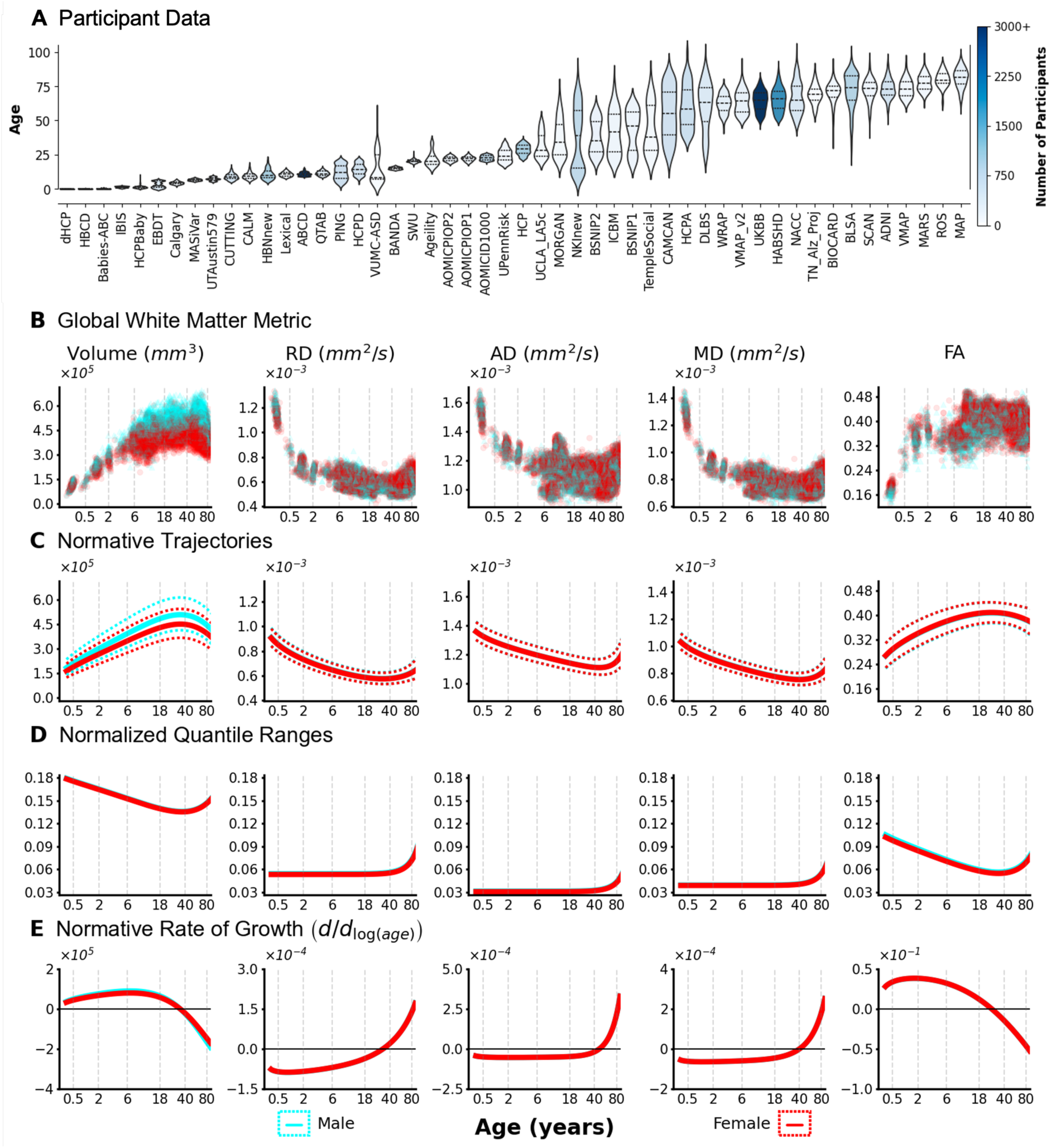
Global White Matter Brain Charts Across the Human Lifespan. A) Data were aggregated from 50 imaging studies, comprising N=35,120 scans from typically developing participants aged 0 to 100 years, forming the basis for the lifespan WM brain charts. Violin plots show the age distribution for each study colored by the number of typically developing/aging participants. B) Raw data points illustrate the distribution of global WM metrics (Volume, FA, MD, AD, RD) across the lifespan for males (cyan) and females (red). C) Normative trajectories for each metric, modeled using GAMLSS, show the median (solid lines) and 2.5th/97.5th percentiles (dotted lines). D) Normalized quantile ranges (difference between 25th and 75th percentiles divided by the median) depict changes in population variability across the lifespan for each global WM feature. E) The normative rate of change (first derivative of the median centile) highlights that growth and decline are most rapid early in life and that different WM features reach their peak or minimum values at distinct ages. The horizontal line (*y* = 0) denotes critical points where the derivative of the trajectories change sign. Note: The x-axes (age in years) are log-scaled to emphasize developmental and aging periods.

Lifespan trajectories of global WM features revealed distinct, feature-specific patterns of growth and decline (**Figure 1b,c**). Cerebral WM volume increased rapidly during early development, peaking at 34 years, followed by gradual atrophy in later life. FA exhibited a similar, though slightly earlier, inflection - rising during childhood and adolescence, peaking around age 24, then steadily declining. In contrast, diffusivity measures (MD, RD, AD) demonstrated inverted trajectories: decreasing steeply during early life, reaching their lowest points in middle adulthood (RD at ∼32 years, MD at ∼40 years, AD at ∼46 years), followed by progressive increases into older age. These divergent trajectories delineate critical inflection points in WM maturation and degeneration.

GAMLSS-estimated population variability revealed distinct age-related trends across WM phenotypes (**Figure 1d**). Variability in metrics of diffusivity remained low or declined during early life, followed by substantial increases during mid- to late adulthood. FA variability peaked during early childhood, declined steadily through midlife, and increased again in older age - suggesting age-dependent fluctuations in structural heterogeneity. Rates of change (**Figure 1e**) were steepest during early childhood, reflecting rapid developmental transitions, and gradually diminished through late childhood and early adolescence (ages 6–12), mirroring the extended maturation period of myelination and axonal remodeling described in histological and neuroimaging studies[16–18].

### Tract-specific White Matter Phenotypes

To move beyond global cerebral measures and characterize regional heterogeneity, we applied the GAMLSS framework to derive tract-specific normative trajectories for white matter microstructural and macrostructural features across the lifespan. We estimated trajectories for 72 anatomically defined WM pathways, capturing the full spectrum of association, commissural, projection, thalamic, and limbic systems. Representative trajectories are shown for five exemplar tracts: the arcuate fasciculus and cingulum bundle (association), corticospinal tract and anterior thalamic radiation (projection), and genu of the corpus callosum (commissural) for microstructural phenotypes (**Figure 2; Supplemental Figures SA.F3-SA.F5, SA.F1** for contralateral) and macrostructural phenotypes (**Figure 3; Supplemental Figures SA.F6,SA.F7, SA.F**2 for contralateral; **Supplemental Figures SA.F8-SA.F16** for trajectories of normalized measures).

**Figure 2.**
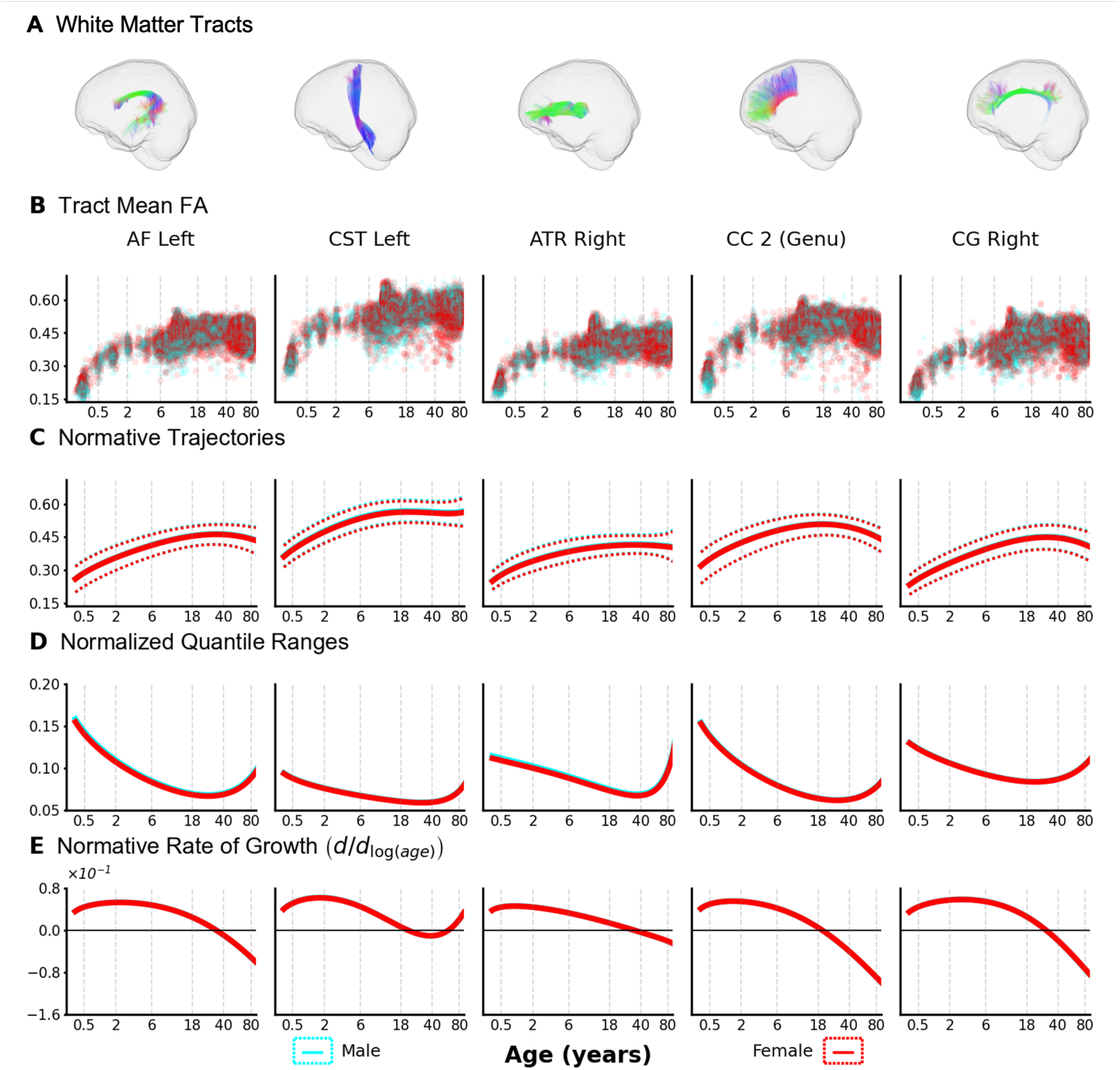
Tract-specific Microstructural Brain Charts. Lifespan brain charts for white matter *microstructure* (FA shown; see **Supplemental Figures SA.F3-SA.F5** for MD, AD, RD; see **Supplemental Figure SA.F1** for contralateral) reveal distinct developmental trajectories and variability across different WM pathways. A) Five exemplar tracts are shown (left to right): left arcuate fasciculus (AF Left); left corticospinal tract (CST Left); right anterior thalamic radiation (ATR Right); genu of the corpus callosum (CC 2 (Genu)); and the right cingulate gyrus (CG Right). B) Raw FA data points for these tracts extend across the lifespan. C) Normative GAMLSS trajectories show FA increasing during development, plateauing in adulthood, and declining in later life, where the timing and magnitude vary by tract. Median (solid lines) and 2.5th/97.5th percentiles (dotted lines) are shown. D) Normalized quantile ranges indicate FA variability is generally lowest in middle age and increases later in life, though developmental variability patterns differ between tracts. E) The normative rate of change (first derivative) suggests that FA peaks at different ages and changes at different rates depending on the specific tract. Note: The x-axes (age in years) are log-scaled to emphasize developmental and aging periods.

**Figure 3.**
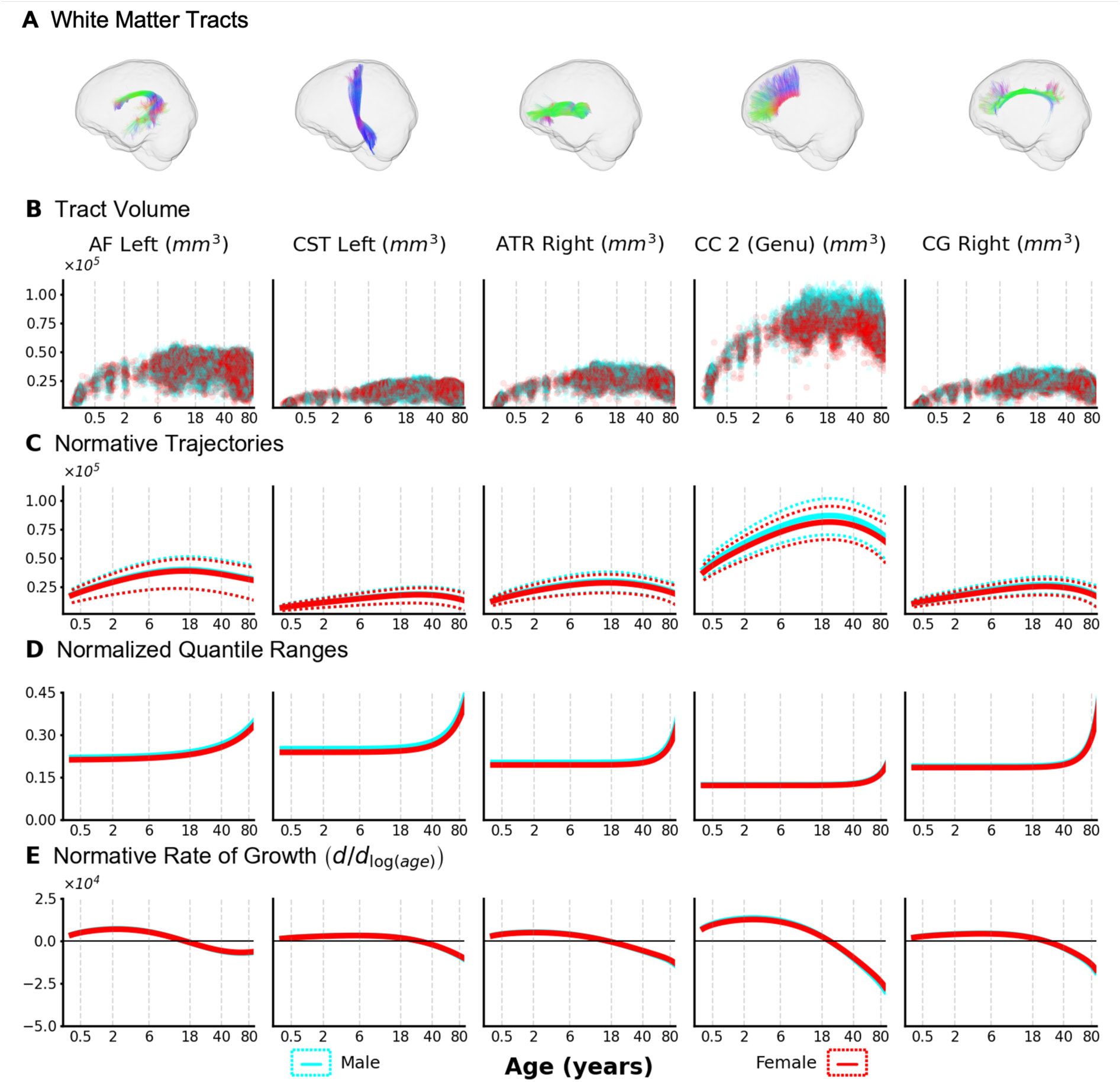
Tract-specific Macrostructural Brain Charts. Lifespan brain charts for white matter *macrostructure* (pathway volume shown; see **Supplemental Figures SA.F4,SA.F5** for length and surface area; see **Supplemental Figure SA.F2** for contralateral) reveal distinct developmental trajectories and variability across different WM pathways. A) Five exemplar tracts are shown (left to right): left arcuate fasciculus (AF Left); left corticospinal tract (CST Left); right anterior thalamic radiation (ATR Right); genu of the corpus callosum (CC 2 (Genu)); and the right cingulate gyrus (CG Right). B) Raw tract volume data points for these tracts span the lifespan, illustrating differences in typical volume ranges between tracts. C) Normative GAMLSS trajectories show tract volume increasing during development, peaking in adolescence or early adulthood, and declining in later life, with tract-specific timing and magnitude. Median (solid lines) and 2.5th/97.5th percentiles (dotted lines) are shown. D) Normalized quantile ranges indicate that volume variability increases later in life and differs between sexes, being generally lowest in middle age. E) The normative rate of change (first derivative) suggests that tract volume peaks at different ages and the rate of subsequent decline varies depending on the specific tract. Note: The x-axes (age in years) are log-scaled to emphasize developmental and aging periods.

Tract-wise trajectories of FA revealed consistent developmental phases but variable temporal profiles across pathways (**Figure 2**). FA increased rapidly during infancy and childhood, reached a plateau in early adulthood, and declined gradually in later life. Yet the magnitude and timing of these inflection points varied substantially by tract. For instance, the corticospinal tract exhibited a prolonged plateau of high anisotropy into midlife, while the genu of the corpus callosum and cingulum bundle reached peak FA earlier, followed by earlier declines. Inter-individual variability in FA followed a similar pattern - remaining low and stable during early development, then increasing with age in a tract-dependent manner. First-derivative estimates highlighted that FA change rates were steepest in early development but differed in timing and slope across pathways.

Macrostructural trajectories of tract volume followed characteristic inverted U-shaped curves (**Figure 3**), with rapid volumetric expansion during early development, peaks occurring in adolescence or early adulthood, and progressive atrophy in older age. As with microstructure, these patterns were highly tract-dependent, with marked differences in peak timing and slope of decline. Inter-individual variability in volume increased with age, and the steepest rates of volumetric change occurred during early childhood. However, the timing and magnitude of these dynamics varied across tracts, reflecting anatomical and developmental diversity in WM maturation and degeneration.

Together, these tract-specific brain charts delineate the heterogeneity of WM development and aging, capturing normative ranges, variability, and rates of change across anatomically distinct pathways. This reference framework enables precise interpretation of age-related trajectories at the pathway level and establishes a foundation for identifying atypical patterns in both developmental and clinical contexts.

### White Matter Developmental Milestones

We extracted tract-resolved milestones from lifespan centile trajectories by locating critical points (local extrema) for microstructural and macrostructural features (Figure 4a). Examples from the left arcuate and right cingulum illustrate two points that generalize: (i) feature-specific timing differs within a tract, and (ii) timing differs across tracts. Across pathways, macrostructural metrics (length, surface area, volume) typically peak earlier than microstructural metrics (**Supplemental Figure SD.F8**). The bottom context panel orients the trajectories within external timelines - marking cohort age coverage, literature-based diagnostic ages for the major disorders represented[19], and developmental transcriptomic windows[20] - highlighting periods of pronounced gene-expression change and clinical vulnerability.

**Figure 4.**
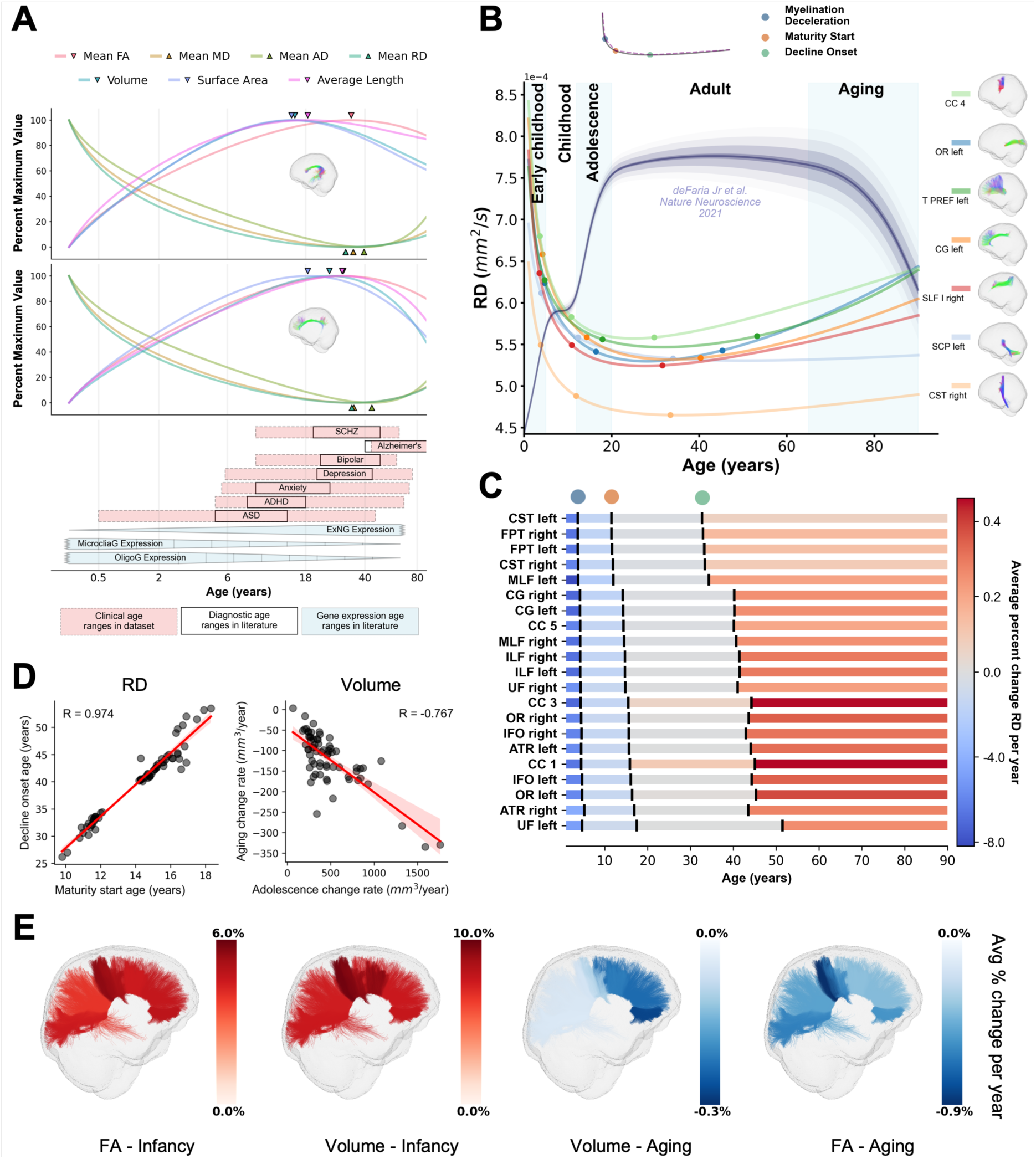
Tract-specific Developmental Milestones, Neurobiological Patterns, and Spatial Gradients. Lifespan trajectories reveal tract-specific developmental milestones and allow for investigation into neurobiological hypotheses and spatio-temporal dynamics of brain microstructure. A.) Median centile trajectories for all features of the left arcuate fasciculus (top) and the right cingulum (middle) show pathway-specific developmental maturation as peaks (downward triangles) or troughs (upward triangles). Below, a graphical summary situates these trajectories alongside non-MRI milestones. Pink shaded boxes denote the age range of clinical disorders represented in the dataset; black boxes denote diagnostic age ranges from literature[19]; blue shaded boxes indicate transcriptomic development windows [20]. B.) Radial diffusivity (RD) median trajectories for exemplar tracts illustrate waves of development and late-life change. These in vivo patterns are consistent with proposed myelin development and aging patterns [22] based on foundational histological work from Yakovlev and Lecours [24] (myelin changes qualitatively visualized as blue curve reproduced from deFaria Jr et al.[22]) C.) Relative timing and magnitude of change for RD trajectories across selected tracts. Vertical black lines represent the identified myelination timepoints (e.g., rapid early myelin increase, maturity start, decline onset). The color of the bars represents the average percent change per year during different life epochs (darker blue indicates a faster decrease; darker red indicates a faster increase), revealing tract-specific variability in rates of change. D.) Neurobiological hypothesis tests derived from tract milestones and rates. Left: last-in-first-out - tracts that mature later tend to show earlier onset of decline (age at maturity vs. age at decline, RD). Right: gain-predicts-loss - greater adolescent growth in volume predicts steeper decline with aging (slope–slope relation). E.) Spatial organization. Anterior-to-posterior gradients emerge in rates of change across tracts (both microstructure and macrostructure), indicating regionally patterned maturation and decline.

We then used the charts to investigate *in vivo* microstructural changes reflecting myelination. To relate milestones to myelination, we used RD [21] as a myelin-sensitive proxy and segmented each RD trajectory piecewise (Methods). This captures the early-life RD decrease (consistent with progressive myelination) and late-life increases, with tract ordering that mirrors classical staging[22–24]: earlier for projection systems (e.g., CST, FPT) and later for association pathways (e.g., UF) (**Figure 4b,c**)[25,26]. These patterns are in line with the established view that human myelination is protracted into mid-adulthood, and proceeds in non-simultaneous waves across systems rather than as a single schedule.[24,27] We summarize the heterogeneity in timing and dynamics across pathways (**Figure 4c**), reporting ages of key RD timepoints intended to reflect distinct waves of myelination[22] (end of early rapid myelination, maturation throughout adolescence, onset of myelin decline) alongside the rates of change between each timepoint. We emphasize RD as a sensitive (not exclusive) *in vivo* proxy for myelination.[28]

Because the charts yield tract-specific milestones, we can quantify links between development and aging (**Figure 4d**). Under a retrogenesis framework[29], we test two hypotheses using tract centile trajectories. Last-In-First-Out posits that later-maturing tracts show earlier onset of decline[30]; operationally, we relate RD-defined age of maturity to RD-defined age at which decline begins. Gain-predicts-loss proposes that the magnitude of developmental growth predicts the magnitude of aging-related decline[31]; operationally, we compared adolescent macrostructural growth rates with later-life atrophy. Both relationships are frequently observed, in both micro- and macro-structure, indicating that the centile trajectories capture principled couplings between maturation and later-life change.

Finally, the charts localize spatial organization of white-matter change (**Figure 4e**).[32] We summarize the mean rate of change for FA and volume in infancy and aging along the sections of the corpus callosum. During infancy, microstructural rates show an anterior–posterior “inside-out” trend, with relatively slower change at the most anterior and posterior pathways and faster change in intermediate systems. During aging, front-to-back volumetric differences emerge with stronger anterior declines. These gradients differ by feature and life stage and are not a mirror image of development, indicating that retrogenesis provides a partial but not complete account of the spatial organization of change.

These results collectively define a fine-grained atlas of white matter developmental milestones. The timing, variability, and rates of change in both microstructural and macrostructural features differ substantially across pathways, offering a comprehensive reference for interpreting normative WM maturation and age-related decline. This framework supports future efforts to investigate how developmental timing relates to functional specialization, vulnerability to disease, and lifespan trajectories of brain structure.

### Individualized Centile Scores of Tract Metrics

We computed individualized centile scores to benchmark each participant’s white matter (WM) measurements against normative age- and sex-stratified trajectories (**Methods**, ‘Centile scores and case–control differences’). These scores quantify the relative typicality or atypicality of microstructural and macrostructural features on a unified scale, with values near the extremes (0th or 100th percentile) reflecting substantial deviation from the normative population. Leveraging a clinically diverse dataset, we applied this framework to systematically examine case–control differences across several diagnostic groups, including neurodevelopmental, neurocognitive, and neurodegenerative disorders (**Figure 5; Supplemental Table SC.T1**).

**Figure 5.**
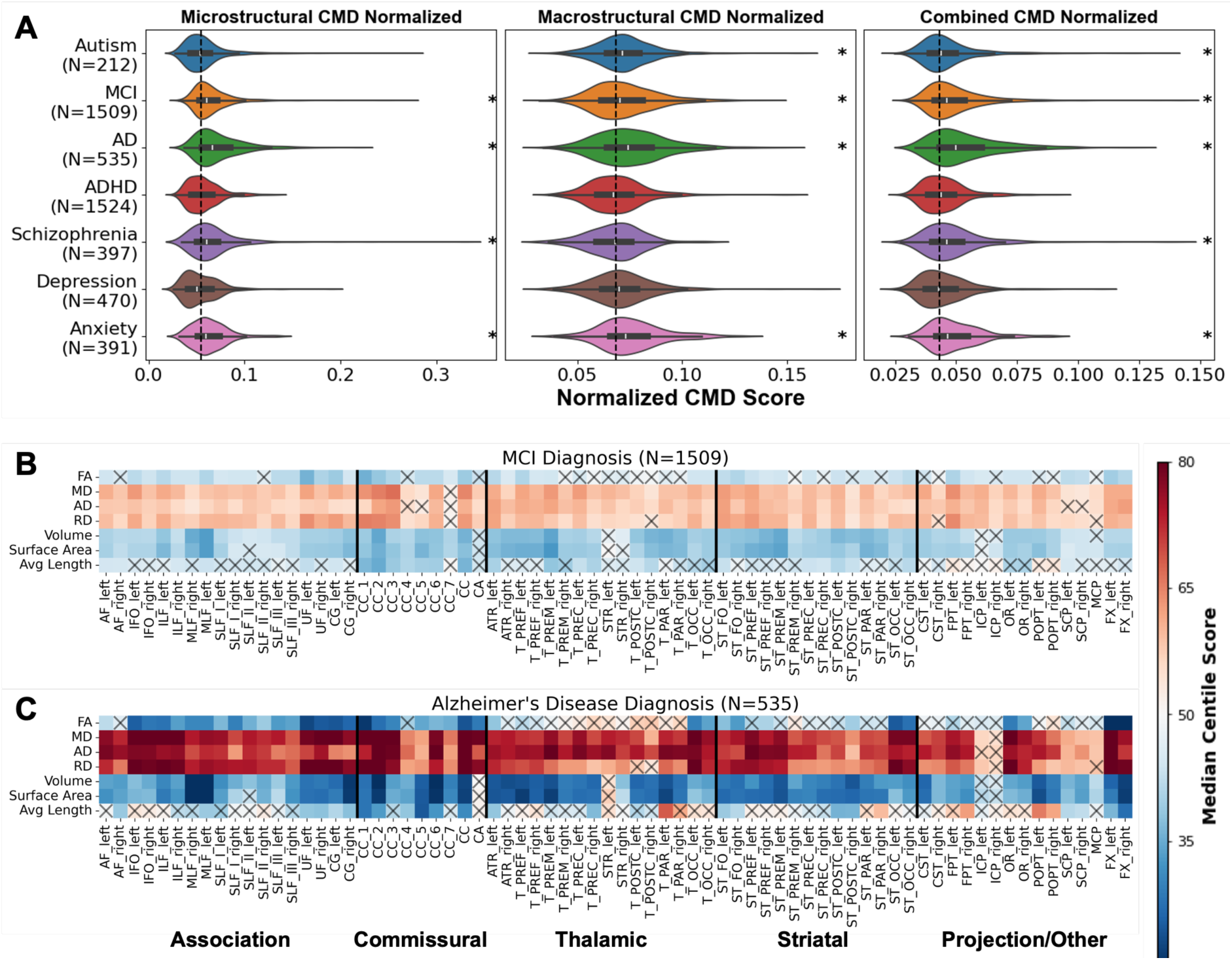
Case-control Differences Quantified by White Matter Centile Scores. Deviations from normative white matter trajectories, measured using individualized centile scores (ranging 0-100, with 50 being the population median), reveal significant differences between typically developing controls and patient populations with various neurological conditions. A.) Violin plots show distributions of the normalized centile Mahalanobis distance (nCMD) scores, an aggregate measure of deviation across all tracts, for microstructural, macrostructural, and combined features in several diagnostic groups. Elevated nCMD scores indicate greater overall deviation from the normative control population (median nCMD for typically developing and aging participants shown as vertical dashed lines). Significant differences from controls (Bonferroni-corrected Wilcoxon test) are marked with an asterisk (*), notably for mild cognitive impairment (MCI), Alzheimer’s disease, and anxiety groups across all aggregated measures (significant at *p* < 0.00714). Heatmaps display the median centile score for each WM feature across all 72 tracts for individuals diagnosed with (B) MCI and (C) Alzheimer’s disease. Colors represent the median centile relative to the control median (blue < 50, red > 50). Widespread deviations are evident, particularly lower FA/volume and higher diffusivities in MCI and AD. Gray “X” indicates features where the median centile was not significantly different from 50 after Bonferroni correction (significant at *p* < 1 ∗ 10^-4^).

To summarize deviations across multiple features and tracts, we computed a normalized centile Mahalanobis distance (nCMD) for each individual, capturing aggregate atypicality within microstructural, macrostructural, and combined feature spaces (Methods, ‘Centile Scores Across Cognitive Groups’). Across diagnostic groups, nCMD values were consistently elevated relative to controls (**Figure 5a**; see **Supplemental Table SC.T2** for effect sizes). This composite score allowed dimensionality reduction while preserving individual variation, revealing disorder-specific patterns - some characterized primarily by microstructural deviation, others by macrostructural or mixed profiles.

Significant deviations from normative centile scores were also observed for specific tract microstructure and macrostructure. Diagnostic groups of mild cognitive impairment (MCI) and Alzheimer’s disease showed significant deviations across most tracts and features compared to neurotypical controls (**Figure 5b,c**; see **Supplemental Figures SC.F1-SC.F3** for other diagnostic groups; see **SC.F4-SC.F6** for effect sizes). MCI was associated with widespread increases in diffusivity, reductions in FA, and lower tract volumes and surface areas—changes that were even more pronounced in Alzheimer’s disease.

These findings demonstrate that individualized centile scores offer a sensitive, interpretable framework for quantifying WM atypicality - capturing both global deviations and pathway-specific abnormalities across diverse neurological diagnostic groups.

### Centile Scoring of New Datasets

A central challenge in applying normative brain charts is estimating individualized centile scores for new, out-of-sample (OOS) datasets not included in the original reference cohort. To address this, we implemented a maximum likelihood estimation (MLE) framework to align external datasets to existing normative trajectories on a feature- and pathway-specific basis (**Figure 6**; **Methods**, ‘Out-of-sample centile estimation’). This alignment enables accurate and interpretable centile scoring, thereby extending the utility of WM brain charts to new datasets for studying both typical and pathological brain structure.

**Figure 6.**
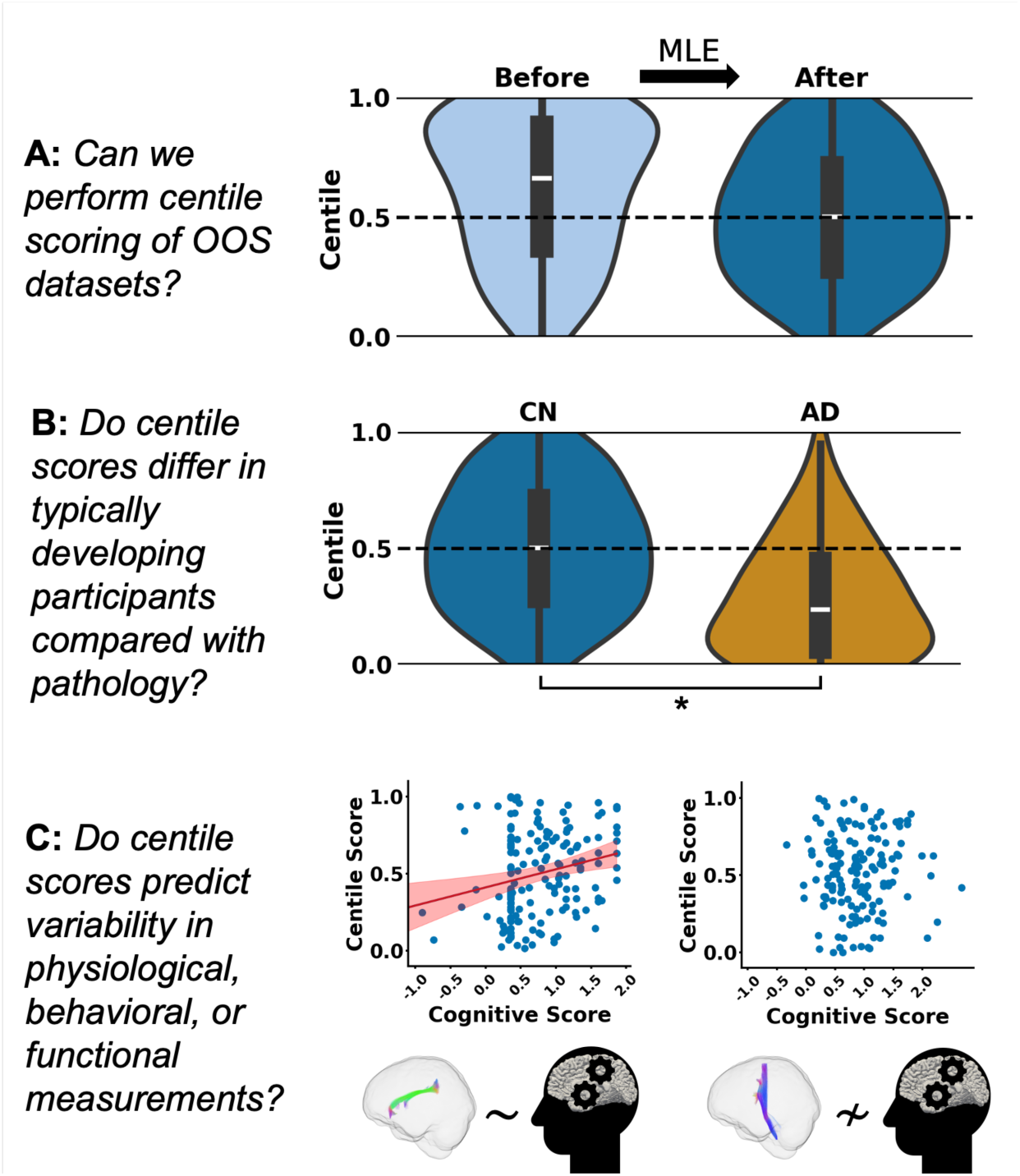
Applications of Out-of-Sample (OOS) Centile Scoring. Normative white matter brain charts enable standardized centile scoring for new OOS datasets, facilitating harmonized analyses. Exemplar applications are demonstrated using the ADNI dataset (withheld from the models used for this figure for demonstration purposes). A) OOS datasets can be aligned to the reference trajectories using maximum likelihood estimation (MLE) on typically developing/aging participants to estimate study-specific offsets. This process yields a correctly centered centile distribution for OOS controls (shown before and after alignment). B) Aligned centile scores allow for standardized comparisons between diagnostic groups within the OOS dataset, illustrated here comparing typically developing/aging, or cognitively normal (CN), individuals and those with Alzheimer’s disease (AD). C) Aligned centile scores serve as standardized metrics to investigate relationships with external variables, such as cognitive performance. The example shows functional scores are significantly correlated with aligned FA centiles in the right SLF III (left plot), but not the right CST (right plot), highlighting pathway-specific associations.

As an initial validation, we estimated study-specific offsets using cognitively healthy controls from a held-out dataset, yielding a near-uniform centile distribution centered around the population median (50th centile; **Figure 6a**). This enabled two key applications in a new Alzheimer’s disease (AD) cohort. First, aligned centile scores provided a consistent framework for evaluating case–control differences, revealing specific white matter features and pathways that significantly diverged in AD relative to controls (**Figure 6b**). Second, the centile framework enabled associations between WM structure and cognitive performance to be examined, identifying pathway-specific features most strongly linked to clinical variability (**Figure 6c**). In this way, the reference charts can be used to relate abnormality indices to a broad range of external clinical data.

Together, these findings demonstrate that centile scoring of out-of-sample datasets provides a rigorous and generalizable framework for evaluating group differences, identifying pathway-specific deviations, and exploring structure-function relationships. By grounding these analyses in normative white matter trajectories, our approach enables biologically interpretable comparisons across diagnostic groups and datasets. (See Supplementary Information for how to perform alignment with an out-of-sample dataset)

## Discussion

We present the first comprehensive normative brain charts of white matter (WM) microstructure and macrostructure spanning the human lifespan, derived from over 35,120 diffusion MRI sessions across 50 harmonized cohorts. These charts establish normative references for both global and tract-specific WM properties, enabling age- and sex-stratified benchmarking of WM development, aging, and pathology. Our analyses delineate normative trajectories, identify tract-specific growth milestones, and demonstrate individualized quantification of WM deviations with applications to diagnostic groups. By addressing prior methodological and dataset limitations, this work marks a critical step toward standardized, lifespan quantification of WM organization.

First, our WM brain charts define robust, sex-stratified trajectories for both microstructural and macrostructural properties across 72 white matter pathways. This includes canonical diffusion metrics (FA, MD, RD, AD) and tract-specific morphometric features such as volume, length, and surface area. Prior lifespan studies have focused largely on gray matter, with established normative models now available for cortical and subcortical volumes (including BrainChart [5], NiChart [8], or CentileBrain [6,7] platforms). While recent efforts have created normative models for WM microstructure based on regional diffusion metrics [33,34], our work extends these frameworks by incorporating both microstructural and macrostructural measures across 72 anatomically defined pathways and providing a more granular and comprehensive view of WM development, aging, and pathology. Our results show that WM pathways can be reliably characterized across development and aging using harmonized tractography pipelines.

While DTI-derived microstructure has been well-studied [26,33,35], tract-level macrostructural features have received limited attention [25]. Here, we demonstrate their value and provide intuitive metrics, such as tract volume and length, as well as derived features describing shape. Together, these benchmarks enable systematic characterization of WM maturation and support broader applications in developmental neuroscience and clinical research.

Second, our normative charts reveal pathway-specific developmental milestones, capturing the diversity in timing and sequence of maturation and decline across WM tracts. These differences reflect both functional specialization and distinct temporal dynamics of WM plasticity, preservation, and degeneration [36]. Furthermore, our normative charts agree with foundational[24] and more recent works of myelin development across the lifespan, not only with the timings of myelination waves, but also the relative myelination timings of certain WM tracts[22]. The open-access trajectories generated here support testing of neurobiological theories [16,30] such as the “last-in, first-out” hypothesis, where later-maturing pathways are more susceptible to early aging, and the “gain-predicts-loss” hypothesis, which posits that regions undergoing rapid developmental expansion are prone to steeper age-related decline. The observed dissociation between microstructural and macrostructural inflection points further underscores the complex, feature-dependent nature of WM development, inviting future work into the temporal interplay of structural metrics across the lifespan.

Third, we demonstrate that normative WM brain charts derived from typically developing and aging individuals with no known neurological or psychiatric conditions provide a sensitive and interpretable framework for detecting individualized anomalies in white matter structure. Microstructural and macrostructural aberrations in cerebral white matter have been implicated in numerous neurological disorders, as referenced in this work. While we have been able to measure many of these parameters for over a decade, the lack of a reliable robust reference atlas has limited our ability to leverage these measurements as a tool for diagnosis and prognosis on an individual patient level. Just as with any blood or body fluid laboratory test, a benchmark of normal values, ideally in a demographically similar population, is critical to interpreting the result in an individual patient. If we hope to utilize these measurements in a clinically productive manner, it is imperative to establish a readily available normative atlas of white matter structural characteristics. Crucially, the GAMLSS framework moves beyond a single median trajectory and explicitly models the full population distribution (including its location, scale, and shape), allowing quantification of inter-individual variability. By quantifying deviations from these distributions using centile scores, we identified distinct patterns of microstructural and macrostructural atypicality in individuals across a variety of diagnostic groups (**Figure 5; Supplemental Figures SC.F1-SC.F3**). These results highlight the utility of centile-based assessments for mapping disease-related alterations at the level of specific tracts and features. While mechanistic inferences are beyond the scope of this study, the sensitivity of this framework to detect subtle and widespread WM abnormalities underscores its potential for clinical applications in both diagnosis and tracking of neurological disorders.

Finally, our framework enables standardized effect size estimation and tract-level quantification of white matter deviation in out-of-sample datasets. By aligning new data to normative trajectories using maximum likelihood estimation, we enable centile scoring even in external cohorts collected across disparate sites and protocols. This facilitates cross-cohort comparisons, phenotype-feature associations, and broader reproducibility across neuroimaging studies. As demonstrated with Alzheimer’s disease cohorts, these aligned centile scores provide a robust foundation for exploring the relationship between WM features and clinical or cognitive outcomes. We note that as the brain charts were constructed with typically aging and developing individuals, aligning out-of-sample datasets requires some typically aging and developing individuals in the out-of-sample dataset as well (See **Supplemental Information**: “Out-of-sample Alignment Analysis”).

Several limitations warrant consideration. Although the aggregated dataset is among the largest to date, age distribution was uneven, with relative under-representation of infancy and mid-adulthood (**Supplemental Information**). To maintain the statistical power of our normative reference model we did not include phenotypes beyond sex (race/ethnicity, genotype status, etc.) due to limited data availability and reduced sample size (see **Supplemental Table SB.T2**).

Further, normative trajectories were derived from cross-sectional data; future longitudinal validation will be important, as individual trajectories can differ from population-level trends.[37] Our cross-sectional approach was chosen to ensure statistical power and model stability for out-of-sample alignment, as detailed in the Supplementary Information (“Considerations for Cross-sectional vs Longitudinal Brain Charts”). Although there was data representation from several continents, the datasets used to create the brain charts are predominantly from European and North American populations (see **Supplemental Table SB.T3**). This bias is unfortunately common in neuroimaging and genetics, echoing limitations in even foundational clinical references like the CDC and WHO growth charts.[11] This highlights a critical need for the global scientific community to increase ethnic, socioeconomic, and demographic diversity in future MRI research. Tractography-based metrics are inherently sensitive to preprocessing parameters and may be less reliable in certain populations or developmental stages [9,38]. Future work may also include alternative bundle segmentation methods [13], with varying levels of specificity and sensitivity [39], alternative definitions and interpretations of pathways, or under-investigated pathways of the brain stem or short association fibers [26]. While we employed conventional DTI for microstructural features, alternative multicompartment diffusion models may offer increased biological specificity [40,41].[7][37] Finally, we caution that these WM brain charts are immediately suitable for the quantitative diagnosis of individual patients in a clinical setting. Significant caveats exist even for the diagnostic interpretation of traditional anthropometric growth charts.[42] Similarly, considerable future research and validation will be necessary to translate these white matter charts from a foundational research framework into a clinically validated diagnostic tool.

In conclusion, we have created comprehensive WM brain charts that define normative microstructural and macrostructural properties across the human lifespan, demonstrated their utility in identifying developmental milestones, detecting abnormalities in clinical populations, and assessing relationships with cognitive and clinical variables. By openly sharing the underlying trajectories, scoring tools, and harmonization pipelines, we aim to accelerate research into the structural basis of cognition, disease, and aging. This work lays the foundation for a new class of precision neuroscience tools grounded in normative white matter architecture. Researchers may find the white matter brain charts and code for aligning out-of-sample datasets in a Docker image at https://zenodo.org/records/17561821.

## Methods

### Data

We aggregated diffusion-weighted imaging (DWI) and T1-weighted (T1w) data from 50 independent studies spanning 0 to 100 years of age (**Supplementary Table ST1**), encompassing 67,723 DWI scans from cognitively normal and clinical participants. All data were converted from DICOM to NIfTI using *dcm2niix* and organized in BIDS format in accordance with previous work [43].

### MRI Processing Pipeline

Diffusion-weighted imaging (DWI) data were preprocessed using the PreQual pipeline[44], which corrects for susceptibility-induced, motion-related, and eddy current distortions, and performs slice-wise signal imputation. Specifically, image denoising was performed using the MRtrix3[45] (version 3.0.4) implementation of Marchenko-Pastur principal component analysis. Following this, TOPUP from the FSL (version 6.0.4) software library[46] is used for susceptibility-distortion correction. For DWI without reverse phase encoding scans acquired, TOPUP is run using a synthetic b0 image created from a T1-weighted scan from the same imaging session via the Synb0-DISCO[47] algorithm (version 3.1). FSL’s EDDY is then used for motion and eddy-current-distortion correction, also performing slice-wise signal imputation with the flag ‘*--repol*’. [44][47]Following preprocessing, diffusion tensor imaging (DTI) models were fit to all volumes with b-values ≤ 1500 s/mm² [41,48] using *dwi2tensor* from MRtrix3. DTI-derived scalar maps - including fractional anisotropy (FA), mean diffusivity (MD), axial diffusivity (AD), and radial diffusivity (RD) - were computed using *tensor2metric* from MRtrix3 [45].

To enable consistent tract segmentation, all diffusion data were resampled to 1 mm isotropic resolution using MRtrix3 [49]. Tractography was performed using TractSeg [50] (version 2.8), which automatically segments 72 anatomically defined cerebral white matter pathways. For each tract, we computed streamline density-weighted averages of DTI metrics (FA, MD, AD, RD) as well as macrostructural features - volume, length, and surface area - using the Scilpy toolkit (version 1.5.0) (https://github.com/scilus/scilpy.git) scripts *scil_compute_bundle_mean_std.py* and *scil_evaluate_bundles_individual_measures.py* respectively. Both raw and ratio-normalized macrostructural features are provided, where volume, surface area, and average length of tracts are normalized to total brain volume (excluding ventricles), estimated total intracranial volume, and cerebral white matter volume.

T1-weighted images were included only when acquired in the same session as DWI data. Brain segmentation was performed using the *recon-all* command from FreeSurfer (v7.2.0) [51], yielding estimates of cerebral white matter volume, brain volume excluding ventricles, and total intracranial volume. For participants aged ≤2 years, we employed *infant_recon_all* from the infant FreeSurfer pipeline [52] (v0.0) to account for age-specific brain morphology. Cerebral WM masks were defined using MRtrix3 [45] 5TT labels, excluding the cerebellum and brainstem. T1w images and corresponding WM segmentations were rigidly registered to DWI space using FSL’s [46] *epi_reg*. Whole-brain WM DTI metrics were then computed by averaging values within the WM mask.

The PreQual preprocessing pipeline can be downloaded at: https://zenodo.org/records/14593034. We also make the entire postprocessing pipeline available at https://zenodo.org/records/17144461.

### Data Selection

Following the approach of Bethlehem et al. [5], we restricted our analysis to cross-sectional data. Participant scans were excluded if key demographic information - age, sex, or cognitive status - was missing or not reported for that participant. For each dataset, the sex demographic was used as reported and encoded as a binary variable, whereas age was a continuous value in years. Regarding infants in the BABIES-ABC, HBCD, and dHCP datasets, we used corrected age based on the due date to better account for prematurity and provide a more accurate developmental context.

Quality control (QC) procedures followed our previously established framework for large-scale diffusion MRI analysis [53]. QC metrics were visually reviewed across *PreQual* preprocessing outputs, FreeSurfer segmentations, and T1w-DWI registration results. Scans were excluded if any preprocessing step failed or was deemed unusable. TractSeg outputs were excluded if more than 12 of the 72 tracts failed to reconstruct. We note it is possible that FreeSurfer failed with successful tractography, and vice-versa, in which cases we chose to retain data from the contrast that passed QA only. The number of sessions and participants retained after QC is detailed in Supplementary Table T1.

Normative trajectories were modeled using only participants labeled as “typically developing” or “control” within their respective studies to reflect patterns of healthy white matter development and aging. Participants who were initially labeled as typically aging but later transitioned to a clinical diagnosis were also excluded when fitting normative trajectories. At the tract level, features were retained only if the tract was reconstructed with the full target of 2,000 streamlines - the default setting in TractSeg.

### Normative Modeling of White Matter Features

We employed generalized additive models for location, scale, and shape (GAMLSS) [14], a flexible regression framework endorsed by the World Health Organization (WHO) for constructing normative growth curves [15,54]. GAMLSS extends generalized linear and additive models by allowing the simultaneous estimation of multiple distribution parameters - not only the mean, but also variance, skewness, and kurtosis - through functions that vary with age and other covariates. This flexibility enables precise modeling of lifespan trajectories for white matter (WM) features.

Formally, GAMLSS defines each parameter of the assumed response distribution through additive predictors:

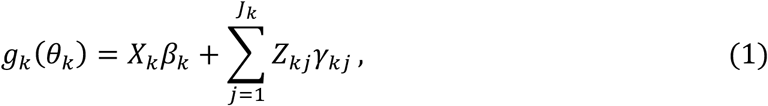

where *g_k_*(⋅) is the link function for the *k*-th parameter, *X_k_* and *β_k_* are the design matrix and fixed effects. The summation incorporates *J_k_* smooth functions where *Z_kj_* are the design matrices for the basis expansions, and *γ_kj_* are the corresponding smoothing coefficients. This formulation allows flexible, non-linear modeling of the entire distribution as a function of covariates such as age, sex, and dataset.

Previous work has shown dMRI-derived features to have skewed distributions. Following Bethlehem et al. [5], we used the generalized gamma (GG) distribution with fractional polynomial (fp) fitting to estimate lifespan trajectories. The GG distribution offers substantial flexibility, accommodating a broad range of distributional shapes, and is therefore suitable for modeling both microstructural and macrostructural imaging features. The model parameters, location (μ), scale (σ), and shape (ν), were defined as:

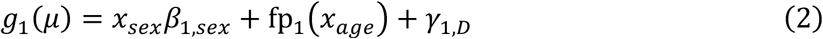

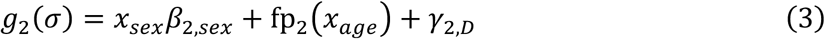

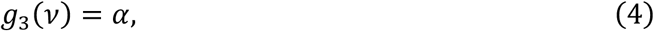

where *g*_1_(⋅)and *g*_2_(⋅) denote log link functions, whereas *g*_3_(⋅) is an identity link. Age was modeled as a continuous predictor using fractional polynomial transformations fp*_k_*(⋅), enabling non-linear representation of age-related effects. Sex was included as a binary fixed effect (*x_sex_*) and dataset-specific variability was modeled via random intercepts (*γ*_1,*D*_ and *γ*_2,*D*_). The GG distribution was parameterized as described in Rigby et al. [55] to enable compatibility with GAMLSS framework. Model selection was performed by the Bayesian Information Criterion (BIC) [56] across all combinations of 1–3 fractional polynomial terms for the *μ* and *σ* parameters. The shape parameter *ν* was treated as a global offset without age- or dataset-specific effects, consistent with Bethlehem et al [5].

We modeled normative trajectories for 509 features, including 288 tract-level (72 tracts × 4 features) microstructural measures (mean FA, MD, AD, RD) and 216 macrostructural (72 tracts × 3 features) features (volume, average length, surface area). An additional five global WM features - mean FA, MD, AD, RD, and WM volume - were also modeled. Macrostructural features were further normalized using total intracranial volume, brain volume excluding ventricles, and global WM volume to account for anatomical scaling.

To derive scaling factors for normalization, we assumed a spherical brain geometry and estimated radius and surface area using:

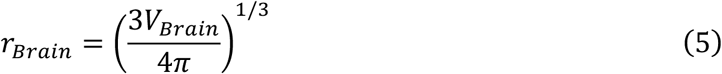

and

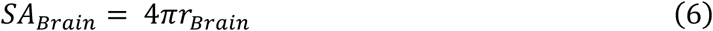

respectively, where *V_Brain_* is the volume of brain tissue excluding ventricles. These derived quantities were used to normalize tract-level volume and surface area metrics accordingly.

When considering both normalized and unnormalized features, we have 1157 features in total that we create lifespan curves for: 509 unnormalized (288 microstructure, 216 macrostructure, and 5 global) and 648 normalized (216 macrostructure normalized in 3 separate ways) features.

### Charting Myelination and Development Across the Lifespan

For assessing myelination, we studied the median RD trajectories of tract-specific brain charts. Following the work from Yeatman et al.[31], we performed piecewise segmentations of the trajectories to assess whether the myelination timepoints align with proposed periods of myelin development and maturation derived from Yakolev and Lecours, Flechsig, and others[22–24]. Specifically, we performed a piecewise segmentation of the median trajectories to identify three break points that minimize the sum of squared errors (SSE) across the resulting linear segments. The segments correspond to myelination periods of rapid development in infancy, slower myelination in adolescence, the stable plateau of young adulthood, and the decline of aging. For each segment, we calculated both the average slope and the percent change per year.

### Centile Scores Across Cognitive Groups

We used the fitted normative trajectories and dataset-specific random effects to compute individualized centile scores for non-control participants across all tract-level features. In total, **2619** participants from **7** clinical groups passed quality control (QC) and were included in the analysis (Supplemental Table S2).

As the analysis was restricted to cross-sectional data, we selected a single scan per participant. For individuals with longitudinal data, we retained the most recent scan corresponding to their most severe clinical diagnosis. For example, participants progressing from cognitively unimpaired to mild cognitive impairment (MCI) and then to Alzheimer’s disease (AD) were classified based on their most recent scan with a clinical AD or dementia diagnosis.

For each clinical group, we compared the distribution of centile scores across tract metrics to the 50th percentile expected in the normative population. In addition, we quantified overall deviation from the normative distribution using the normalized centile Mahalanobis distance (nCMD) for each participant *j* (nCMD*_j_*):

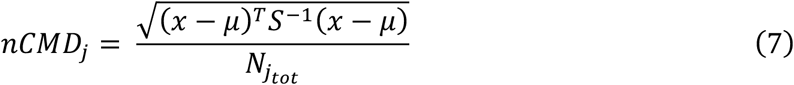

where *N_jtot_* is the number of reconstructed tract features for participant *j*, *x* is a vector of centile scores (ranging from 0 to 1), *μ* = 0.5 denotes the median centile, and *S*^-1^ is the inverse of the covariance matrix for the features.

We computed nCMD separately for tract-level microstructural features, macrostructural features, and their combination. Group-level comparisons were then made by evaluating the median nCMD in each non-control group relative to the distribution observed in the cognitively normal population, where statistical significance was tested using a one-sample Wilcoxon test (two-sided) after Bonferroni correction for multiple comparisons.

### Maximum Likelihood Estimation for Out-of-Sample Datasets

A central utility of normative brain charts is their use as reference models for external, out-of-sample datasets. To enable this, new datasets must be aligned to the fitted trajectories by estimating study-specific offsets for the distributional parameters. Within the GAMLSS framework, each dataset *D* is modeled with random effect terms *γ*_1,*D*_ and *γ*_2,*D*_, which account for dataset-specific variabilities in the location (*μ*) and scale (*σ*) parameters, respectively. For an unseen dataset, *S*, alignment involves estimating these two unknown offsets. We demonstrate this process using the ADNI dataset, where we fit additional models without ADNI for the purpose of demonstrating out-of-sample alignment.

For a given brain chart modeling a tract-level metric, the GAMLSS-based lifespan distribution is defined as:

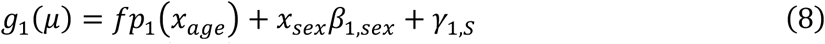

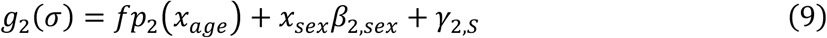

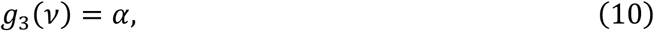

where *γ*_1,*S*_ and *γ*_2.*S*_ are dataset-specific random effects for the ADNI dataset, and all other model parameters are fixed from the trained GAMLSS model fitted without ADNI data.

As all other effects are known from the reference model, alignment requires estimating only the unknown random effects *γ*_1,*W*_ and *γ*_2,*W*_. We initialized both to zero and restricted the estimation procedure to typically aging individuals in the ADNI cohort. To minimize diagnostic confounding, we selected only participants with no history or future diagnosis of MCI or Alzheimer’s disease and used their earliest available scan. This cognitively normal subset is denoted *S_CN_*.

For each cognitively normal participant *j* in *S_CN_*, we computed the likelihood of their observed metric *M_j_* under the GG distribution defined by Equations (8) – (10). Each of the resulting probability densities *P_j_* = *GG*(*M_j_* | *μ*, *σ*, *υ*) were used to calculate the overall negative log-likelihood across all participants in *S_CN_*:

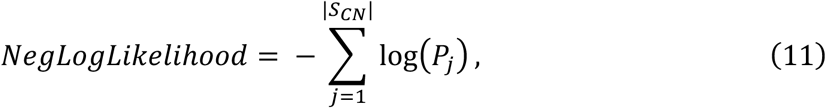

We iteratively re-estimated *γ_1,W_* and *γ*_2,*W*_ by minimizing the negative log-likelihood until convergence. The final estimates 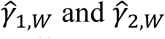, were then applied to all participants in the WASHU dataset to compute adjusted centile scores aligned to the normative reference space.

## Acknowledgements

This work was supported in part by the National Institute of Health through NIH awards K01-EB032898 (Schilling) and K01-AG073584 (Archer), grant number 1R01EB017230-01A1 (Landman), K24-AG046373 (Jefferson), and ViSE/VICTR VR3029, UL1-TR000445, and UL1-TR002243. This work was supported by NIA grants R01-AG034962 (Jefferson), R01-AG056534 (Jefferson), R01-AG062826, and Alzheimer’s Association IIRG-08-88733 (Jefferson) and the NICHD, R01-HD114489 (Vinci-Booher). This work was supported by the Alzheimer’s Disease Sequencing Project Phenotype Harmonization Consortium (ADSP-PHC) that is funded by NIA (U24 AG074855, U01 AG068057 and R01 AG059716). This work was conducted in part using the resources of the Advanced Computing Center for Research and Education (ACCRE) at Vanderbilt University, Nashville, TN. We appreciate the National Institute of HealthS10 Shared Instrumentation grant 1S10OD020154-01, and grant 1S10OD023680-01 (Vanderbilt’s High-Performance Computer Cluster for Biomedical Research). This work was also supported in part by Intramural Research Program of the National Institute on Aging, NIH.

BABIES and ABC datasets were supported by the Jacobs Foundation Early Career Research Fellowship (2017-1261-05) (Humphreys); National Science Foundation CAREER Award (2042285) (Humphreys); Brain and Behavior Research Foundation John and Polly Sparks Foundation Investigator Award (29593) (Humphreys); Vanderbilt Institute for Clinical and Translational Research Grant (VR53419) (Humphreys); Vanderbilt Strong Grant; Vanderbilt Kennedy Center Grant (Humphreys); National Institute of Mental Health (R01MH129634) (Humphreys).

This work was supported by the NIH NINDS through award numbers R01 NS108445 and R01 NS110130 (Morgan), and R01 NS134625 and R01 NS112252 (Englot). Data collected from the MORGAN dataset were acquired at Vanderbilt University, with the aim of studying cognitive patterns in patients with epilepsy both before and after clinical treatment.

This work was supported by the NIH Eunice Kennedy Shriver National Institute Of Child Health & Human Development through grant number F31 HD104385 (Nguyen), P50HD103537 (PI: Neul), R37 HD095519 MERIT Award, R01 HD089474, R01 HD109151, R01 HD067254, and R01 HD044073 (Cutting). This work was also supported by the NIH NINDS through grant number R01 NS049096 (Cutting).

Research reported in this publication was supported by NIGMS of the National Institutes of Health under award number T32GM007347 and T32GM152284.

Data and/or research tools used in the preparation of this manuscript were obtained from the National Institute of Mental Health (NIMH) Data Archive (NDA). NDA is a collaborative informatics system created by the National Institutes of Health to provide a national resource to support and accelerate research in mental health. This manuscript reflects the views of the authors and may not reflect the opinions or views of the NIH or of the Submitters submitting original data to NDA.

The Pediatric Imaging, Neurocognition, and Genetics (PING) dataset was collected and released openly to contribute to the assessment of typical brain development in a pediatric sample (RC2DA029475-01), (https://www.sciencedirect.com/science/article/pii/S1053811915003572).

The data used in this study come from the Human Connectome Project, which aims to map the structural connections and circuits of the brain and their relationships to behavior by acquiring high-quality magnetic resonance images. We used diffusion MRI data from the Baby Connectome Project (HCPBaby), the Human Connectome Project Development (HCPD) study, the Human Connectome Project Young Adult (HCP) study, and the Human Connectome Project Aging (HCPA) study.

Research data from the Infant Brain Imaging Study (IBIS) dataset come from the IBIS Autism project, a collaborative effort by investigators to conduct a longitudinal MRI/DTI and behavioral study of infants at high risk for autism (R01HD055741-01).

The Enhanced NKI-RS is a large cross-sectional sample of brain development, maturation and aging, that is currently funded by the NIMH (BRAINS R01MH094639-01; PI Milham) and Child Mind Institute (PI Milham) to characterize 1000 community-ascertained participants using state-of-the-art multiplex imaging-based resting state fMRI (R-fMRI) and diffusion tensor imaging (DTI), genetics, and a broad neurobehavioral phenotypic characterization protocol. Data were acquired from the DSI studio website: (https://brain.labsolver.org/nki_rockland.html).

The Healthy Brain Network (HBN) is an ongoing initiative focused on building a biobank of data from 10,000 children and adolescents (ages 5-21) in the New York City area (https://www.nature.com/articles/sdata2017181). Data were acquired from the DSI studio website: (https://brain.labsolver.org/hbn.html).

Data collection and sharing for this project was provided by the Cambridge Centre for Ageing and Neuroscience (CamCAN). CamCAN funding was provided by the UK Biotechnology and Biological Sciences Research Council (grant number BB/H008217/1), together with support from the UK Medical Research Council and University of Cambridge, UK. Data used in the preparation of this work were obtained from the CamCAN repository (available at http://www.mrc-cbu.cam.ac.uk/datasets/camcan/). The Cambridge Centre for Ageing and Neuroscience (Cam-CAN) is a large-scale collaborative research project at the University of Cambridge.

VMAP data were collected by Vanderbilt Memory and Alzheimer’s Center Investigators at Vanderbilt University Medical Center. VMAP began in 2012 with the goal of investigating vascular health and brain aging.

BLSA is a prospective cohort study with continuous enrollment that began in 1958. Comprehensive data from BLSA are available upon request by a proposal submission through the cohort website (www.blsa.nih.gov). The BLSA is supported by the Intramural Research Program of the National Institute on Aging, NIH.

The BIOCARD study is designed to identify biomarkers associated with progression from normal cognitive status to cognitive impairment or dementia, with a particular focus on Alzheimer’s Disease.

Data collection and sharing for ADNI were supported by National Institutes of Health Grant U01-AG024904 and Department of Defense (award number W81XWH-12-2-0012). ADNI is also funded by the National Institute on Aging, the National Institute of Biomedical Imaging and Bioengineering, and through generous contributions from the following: AbbVie, Alzheimer’s Association; Alzheimer’s Drug Discovery Foundation; Araclon Biotech; BioClinica, Inc.; Biogen; Bristol-Myers Squibb Company; CereSpir, Inc.; Cogstate; Eisai Inc.; Elan Pharmaceuticals, Inc.; Eli Lilly and Company; EuroImmun; F. Hoffmann-La Roche Ltd and its affiliated company Genentech, Inc.; Fujirebio; GE Healthcare; IXICO Ltd.; Janssen Alzheimer Immunotherapy Research & Development, LLC.; Johnson & Johnson Pharmaceutical Research & Development LLC.; Lumosity; Lundbeck; Merck & Co., Inc.; Meso Scale Diagnostics, LLC.; NeuroRx Research; Neurotrack Technologies; Novartis Pharmaceuticals Corporation; Pfizer Inc.; Piramal Imaging; Servier; Takeda Pharmaceutical Company; and Transition Therapeutics. The Canadian Institutes of Health Research is providing funds to support ADNI clinical sites in Canada. Private sector contributions are facilitated by the Foundation for the National Institutes of Health (www.fnih.org). The grantee organization is the Northern California Institute for Research and Education, and the study is coordinated by the Alzheimer’s Therapeutic Research Institute at the University of Southern California. ADNI data are disseminated by the Laboratory for Neuro Imaging at the University of Southern California. Data used in the preparation of this article were obtained from the Alzheimer’s Disease Neuroimaging Initiative (ADNI) database (adni.loni.usc.edu). The ADNI was launched in 2003 as a public private partnership, led by Principal Investigator Michael W. Weiner, MD. The original goal of ADNI was to test whether serial magnetic resonance imaging (MRI), positron emission tomography (PET), other biological markers, and clinical and neuropsychological assessment can be combined to measure the progression of mild cognitive impairment (MCI) and early Alzheimer’s disease (AD). The current goals include validating biomarkers for clinical trials, improving the generalizability of ADNI data by increasing diversity in the participant cohort, and to provide data concerning the diagnosis and progression of Alzheimer’s disease to the scientific community. For up-to-date information, see adni.loni.usc.edu.

Research reported in this publication was supported by the National Institute on Aging of the National Institutes of Health under Award Numbers R01AG054073 and R01AG058533, R01AG070862, P41EB015922 and U19AG078109. The content is solely the responsibility of the authors and does not necessarily represent the official views of the National Institutes of Health.

The NACC database is funded by NIA/NIH Grant U24 AG072122. NACC data are contributed by the NIA-funded ADRCs: P30 AG062429 (PI James Brewer, MD, PhD), P30 AG066468 (PI Oscar Lopez, MD), P30 AG062421 (PI Bradley Hyman, MD, PhD), P30 AG066509 (PI Thomas Grabowski, MD), P30 AG066514 (PI Mary Sano, PhD), P30 AG066530 (PI Helena Chui, MD), P30 AG066507 (PI Marilyn Albert, PhD), P30 AG066444 (PI John Morris, MD), P30 AG066518 (PI Jeffrey Kaye, MD), P30 AG066512 (PI Thomas Wisniewski, MD), P30 AG066462 (PI Scott Small, MD), P30 AG072979 (PI David Wolk, MD), P30 AG072972 (PI Charles DeCarli, MD), P30 AG072976 (PI Andrew Saykin, PsyD), P30 AG072975 (PI David Bennett, MD), P30 AG072978 (PI Ann McKee, MD), P30 AG072977 (PI Robert Vassar, PhD), P30 AG066519 (PI Frank LaFerla, PhD), P30 AG062677 (PI Ronald Petersen, MD, PhD), P30 AG079280 (PI Eric Reiman, MD), P30 AG062422 (PI Gil Rabinovici, MD), P30 AG066511 (PI Allan Levey, MD, PhD), P30 AG072946 (PI Linda Van Eldik, PhD), P30 AG062715 (PI Sanjay Asthana, MD, FRCP), P30 AG072973 (PI Russell Swerdlow, MD), P30 AG066506 (PI Todd Golde, MD, PhD), P30 AG066508 (PI Stephen Strittmatter, MD, PhD), P30 AG066515 (PI Victor Henderson, MD, MS), P30 AG072947 (PI Suzanne Craft, PhD), P30 AG072931 (PI Henry Paulson, MD, PhD), P30 AG066546 (PI Sudha Seshadri, MD), P20 AG068024 (PI Erik Roberson, MD, PhD), P20 AG068053 (PI Justin Miller, PhD), P20 AG068077 (PI Gary Rosenberg, MD), P20 AG068082 (PI Angela Jefferson, PhD), P30 AG072958 (PI Heather Whitson, MD), P30 AG072959 (PI James Leverenz, MD).

This research has been conducted using the UK Biobank resource, application 16315.

Data contributed from MAP/ROS/MARS was supported by NIA R01AG017917, P30AG10161, P30AG072975, R01AG022018, R01AG056405, UH2NS100599, UH3NS100599,

R01AG064233, R01AG15819 and R01AG067482, and the Illinois Department of Public Health (Alzheimer’s Disease Research Fund). Data can be accessed at www.radc.rush.edu. More information about participant demographics and study information can be found here: https://www.rushu.rush.edu/research-rush-university/departmental-research/rush-alzheimers-disease-center/rush-alzheimers-disease-center-research/epidemiologic-research.

The data contributed from the Wisconsin Registry for Alzheimer’s Prevention was supportedby NIA AG021155, AG0271761, AG037639, and AG054047.

We use generative AI to create code segments based on task descriptions, as well as debug, edit, and autocomplete code. Additionally, generative AI technologies have been employed to assist in structuring sentences and performing grammatical checks. It is imperative to highlight that the conceptualization, ideation, and all prompts provided to the AI originate entirely from the authors’ creative and intellectual efforts. We take accountability for the review of all content generated by AI in this work.

Data collection and sharing for this project was provided by the Centre for Attention, Learning and Memory (CALM). CALM funding was provided by the UK Medical Research Council and University of Cambridge, UK. Data used in the preparation of this work were obtained from CALM resource – https://calm.mrc-cbu.cam.ac.uk/. The study protocol is reported in Holmes et al. (2019).

Data used in the preparation of this work were obtained from the International Consortium for Brain Mapping (ICBM) database (www.loni.usc.edu/ICBM). The ICBM project (Principal Investigator John Mazziotta, M.D., University of California, Los Angeles) is supported by the National Institute of Biomedical Imaging and BioEngineering. ICBM is the result of efforts of co-investigators from UCLA, Montreal Neurologic Institute, University of Texas at San Antonio, and the Institute of Medicine, Juelich/Heinrich Heine University - Germany. Data collection and sharing for this project was provided by the International Consortium for Brain Mapping (ICBM; Principal Investigator: John Mazziotta, MD, PhD). ICBM funding was provided by the National Institute of Biomedical Imaging and BioEngineering. ICBM data are disseminated by the Laboratory of Neuro Imaging at the University of Southern California.

The dataset that we refer to as UTAustin in this manuscript is a longitudinal neuroimaging dataset on language processing in children ages 5, 7, and 9 years old collected at The University of Texas at Austin. The dataset is openly available from Openneuro here: https://openneuro.org/datasets/ds003604/versions/1.0.7. We use version 1.0.7.

The Queensland Twin Adolescent Brain (QTAB) Project was established with the purpose of promoting the conduct of health-related research in adolescence. The QTAB dataset comprises multimodal neuroimaging, as well as cognitive and mental health data collected in adolescent twins over two sessions. The MRI protocol consisted of T1-weighted (MP2RAGE), T2-weighted, FLAIR, high-resolution TSE, SWI, resting-state fMRI, DWI, and ASL scans. The QTAB project resource was produced as a result of i) the goodwill and contribution of 422 twin/triplet participants and their parents, ii) funding from the National Health and Medical Research Council, Australia (APP1078756) and the Queensland Brain Institute, University of Queensland, iii) access to several key resources, including the Centre for Advanced Imaging, the Human Studies Unit, Institute of Molecular Bioscience, and the Queensland Cyber Infrastructure Foundation, at the University of Queensland, local and national twin registries at the QIMR Berghofer Medical Research Institute and Twin Research Australia, as well as the many assessments made available by researchers worldwide, and iv) was established with the purpose of promoting the conduct of health-related research in adolescence. The imaging data and basic demographics are openly accessibly on Openneuro here: https://openneuro.org/datasets/ds004146/versions/1.0.4. We use version 1.0.4.

The Social Reward and Nonsocial Reward Processing Across the Adult Lifespan: An Interim Multi-echo fMRI and Diffusion Dataset (referred to as TempleSocial in this manuscript) comes from a study that aims to investigate whether older adults have a blunted response to some features of social reward. The dataset is openly accessibly on Openneuro here: https://openneuro.org/datasets/ds005123/versions/1.1.3. In particular, we use version 1.1.3.

The Longitudinal Brain Correlates of Multisensory Lexical Processing in Children study (shortened to Lexical in this manuscript) aims to explore developmentally dependent changes in lexical processing for adolescents. The dataset is openly accessibly on Openneuro here: https://openneuro.org/datasets/ds001894/versions/1.4.2. We use version 1.4.2.

The UCLA Consortium for Neuropsychiatric Phenomics LA5c Study (UCLA) is focused on understanding the dimensional structure of memory and cognitive control (response inhibition) functions in both healthy individuals and individuals with neuropsychiatric disorders including schizophrenia, bipolar disorder, and attention deficit/hyperactivity disorder. Neuroimaging data were downloaded from Openneuro here: https://openneuro.org/datasets/ds000030/versions/1.0.0. We use version 1.0.0.

The dataset we refer to as UPennRisk comes from a study at University of Pennsylvania that investigated whether training executive cognitive function could influence choice behavior and brain responses. Neuroimaging data were downloaded from Openneuro here: https://openneuro.org/datasets/ds002843/versions/1.0.1. We use version 1.0.1.

The Dallas Lifespan Brain Study (DLBS) is a longitudinal multi-modal neuroimaging study of the aging mind, which was initiated in 2008 (referred to as Wave 1). Participants returned for two additional waves of data collection with an approximate interval of 4-5 years between waves. The DLBS protocol encompasses various imaging modalities, including structural MRI, diffusion MRI, and functional MRI, as well as comprehensive cognitive and psychosocial assessments. DLBS data can be downloaded from Openneuro here: https://openneuro.org/datasets/ds004856. Specifically, we use version 1.2.0.

The Multisite, Multiscanner, and Multisubject Acquisitions for Studying Variability in Diffusion Weighted Magnetic Resonance Imaging (MASiVar) dataset consists of 319 diffusion scans acquired at 3T from b = 1000 to 3000 s/mm2 across 14 healthy adults, 83 healthy children (5 to 8 years), three sites, and four scanners curated to promote investigation of diffusion MRI variability. In particular, we used only the data coming from healthy children (Cohort IV) for version 2.0.2 of the dataset. Data are available to download from Openneuro here: https://openneuro.org/datasets/ds003416/versions/2.0.2.

Data coming from the Southwestern University (SWU) dataset, part of the Consortium for Reliability and Reproducibility (CoRR), were downloaded via NITRC-IR from the 1000 Functional Connectomes Project. Specifically, the data come from the Emotion and Creativity One Year Retest Dataset subset, comprised of 235 subjects, all of whom were college students. Each subject underwent two sessions of anatomical, resting state fMRI, and DTI scans, spaced one year apart. In order to access the CoRR datasets through NITRC, users must be logged into NITRC at the time of download and registered with the 1000 Functional Connectomes Project / INDI website. More information about this subset can be found here: https://fcon_1000.projects.nitrc.org/indi/CoRR/html/swu_4.html.

The Preschool MRI study in The Developmental Neuroimaging Lab at the University of Calgary (https://www.developmentalneuroimaginglab.ca) uses different magnetic resonance imaging (MRI) techniques to study brain structure and function in early childhood. The study aims to characterize brain development in early childhood, and to offer baseline data that can be used to understand cognitive and behavioural development, as well as to identify deviations from normal development in children with various diseases, disorders, or brain injuries. The MRI techniques used include diffusion tensor imaging (DTI), anatomical imaging, arterial spin labeling (ASL), and resting state functional MRI (rsfMRI). Data can be downloaded from here: https://osf.io/axz5r/.

The Amsterdam Open MRI Collection (AOMIC) is a collection of three datasets with multimodal (3T) MRI data including structural (T1-weighted), diffusion-weighted, and (resting-state and task-based) functional BOLD MRI data, as well as detailed demographics and psychometric variables from a large set of healthy participants. All raw data is publicly available from the Openneuro data sharing platform: ID1000: https://openneuro.org/datasets/ds003097, PIOP1: https://openneuro.org/datasets/ds002785, PIOP2: https://openneuro.org/datasets/ds002790. We use version 1.2.1 for ID1000 and 2.0.0 for PIOP1 and PIOP2.

The Boston Adolescent Neuroimaging of Depression and Anxiety (BANDA) is a study of 215 adolescents ages 14-17, 152 of whom had a current diagnosis of a DSM-5 (APA, 2013) anxious and/or depressive disorder. The BANDA study collected a rich dataset of brain, clinical, and cognitive/neuropsychological measures from these adolescent subjects. The dataset is available to download upon request on the NDA.

We thank Knight ADRC for providing neuroimaging data to us (WASHU dataset). The data contributed through WASHU (Knight ADRC) was supported by grant numbers P30 AG066444, P01 AG03991, and P01 AG026276. For WASHU, Clinical Dementia Ratings (CDRs) are obtained from assessments by experienced clinicians trained in the use of the CDR.

The NACC database is funded by NIA/NIH Grant U24 AG072122. SCAN is a multi-institutional project that was funded as a U24 grant (AG067418) by the National Institute on Aging in May 2020. Data collected by SCAN and shared by NACC are contributed by the NIA-funded ADRCs as follows:

Arizona Alzheimer’s Center - P30 AG072980 (PI: Eric Reiman, MD); R01 AG069453 (PI: Eric Reiman (contact), MD); P30 AG019610 (PI: Eric Reiman, MD); and the State of Arizona which provided additional funding supporting our center; Boston University - P30 AG013846 (PI Neil Kowall MD); Cleveland ADRC - P30 AG062428 (James Leverenz, MD); Cleveland Clinic, Las Vegas – P20AG068053; Columbia - P50 AG008702 (PI Scott Small MD); Duke/UNC ADRC – P30 AG072958; Emory University - P30AG066511 (PI Levey Allan, MD, PhD); Indiana University - R01 AG19771 (PI Andrew Saykin, PsyD); P30 AG10133 (PI Andrew Saykin, PsyD); P30 AG072976 (PI Andrew Saykin, PsyD); R01 AG061788 (PI Shannon Risacher, PhD); R01 AG053993 (PI Yu-Chien Wu, MD, PhD); U01 AG057195 (PI Liana Apostolova, MD); U19 AG063911 (PI Bradley Boeve, MD); and the Indiana University Department of Radiology and Imaging Sciences; Johns Hopkins - P30 AG066507 (PI Marilyn Albert, Phd.); Mayo Clinic - P50 AG016574 (PI Ronald Petersen MD PhD); Mount Sinai - P30 AG066514 (PI Mary Sano, PhD); R01 AG054110 (PI Trey Hedden, PhD); R01 AG053509 (PI Trey Hedden, PhD); New York University - P30AG066512-01S2 (PI Thomas Wisniewski, MD); R01AG056031 (PI Ricardo Osorio, MD); R01AG056531 (PIs Ricardo Osorio, MD; Girardin Jean-Louis, PhD); Northwestern University - P30 AG013854 (PI Robert Vassar PhD); R01 AG045571 (PI Emily Rogalski, PhD); R56 AG045571, (PI Emily Rogalski, PhD); R01 AG067781, (PI Emily Rogalski, PhD); U19 AG073153, (PI Emily Rogalski, PhD); R01 DC008552, (M.-Marsel Mesulam, MD); R01 AG077444, (PIs M.-Marsel Mesulam, MD, Emily Rogalski, PhD); R01 NS075075 (PI Emily Rogalski, PhD); R01 AG056258 (PI Emily Rogalski, PhD); Oregon Health and Science University - P30 AG008017 (PI Jeffrey Kaye MD); R56 AG074321 (PI Jeffrey Kaye, MD); Rush University - P30 AG010161 (PI David Bennett MD); Stanford – P30AG066515; P50 AG047366 (PI Victor Henderson MD MS); University of Alabama, Birmingham – P20; University of California, Davis - P30 AG10129 (PI Charles DeCarli, MD); P30 AG072972 (PI Charles DeCarli, MD); University of California, Irvine - P50 AG016573 (PI Frank LaFerla PhD); University of California, San Diego - P30AG062429 (PI James Brewer, MD, PhD); University of California, San Francisco - P30 AG062422 (Rabinovici, Gil D., MD); University of Kansas - P30 AG035982 (Russell Swerdlow, MD); University of Kentucky - P30 AG028283-15S1 (PIs Linda Van Eldik, PhD and Brian Gold, PhD); University of Michigan ADRC - P30AG053760 (PI Henry Paulson, MD, PhD) P30AG072931 (PI Henry Paulson, MD, PhD) Cure Alzheimer’s Fund 200775 - (PI Henry Paulson, MD, PhD) U19 NS120384 (PI Charles DeCarli, MD, University of Michigan Site PI Henry Paulson, MD, PhD) R01 AG068338 (MPI Bruno Giordani, PhD, Carol Persad, PhD, Yi Murphey, PhD) S10OD026738-01 (PI Douglas Noll, PhD) R01 AG058724 (PI Benjamin Hampstead, PhD) R35 AG072262 (PI Benjamin Hampstead, PhD) W81XWH2110743 (PI Benjamin Hampstead, PhD) R01 AG073235 (PI Nancy Chiaravalloti, University of Michigan Site PI Benjamin Hampstead, PhD) 1I01RX001534 (PI Benjamin Hampstead, PhD) IRX001381 (PI Benjamin Hampstead, PhD); University of New Mexico - P20 AG068077 (Gary Rosenberg, MD); University of Pennsylvania - State of PA project 2019NF4100087335 (PI David Wolk, MD); Rooney Family Research Fund (PI David Wolk, MD); R01 AG055005 (PI David Wolk, MD); University of Pittsburgh - P50 AG005133 (PI Oscar Lopez MD); University of Southern California - P50 AG005142 (PI Helena Chui MD); University of Washington - P50 AG005136 (PI Thomas Grabowski MD); University of Wisconsin - P50 AG033514 (PI Sanjay Asthana MD FRCP); Vanderbilt University – P20 AG068082; Wake Forest - P30AG072947 (PI Suzanne Craft, PhD); Washington University, St. Louis - P01 AG03991 (PI John Morris MD); P01 AG026276 (PI John Morris MD); P20 MH071616 (PI Dan Marcus); P30 AG066444 (PI John Morris MD); P30 NS098577 (PI Dan Marcus); R01 AG021910 (PI Randy Buckner); R01 AG043434 (PI Catherine Roe); R01 EB009352 (PI Dan Marcus); UL1 TR000448 (PI Brad Evanoff); U24 RR021382 (PI Bruce Rosen); Avid Radiopharmaceuticals / Eli Lilly; Yale - P50 AG047270 (PI Stephen Strittmatter MD PhD); R01AG052560 (MPI: Christopher van Dyck, MD; Richard Carson, PhD); R01AG062276 (PI: Christopher van Dyck, MD); 1Florida - P30AG066506-03 (PI Glenn Smith, PhD); P50 AG047266 (PI Todd Golde MD PhD)

Data used in the preparation of this article were obtained from the HEALthy Brain and Child Development (HBCD) Study (https://hbcdstudy.org/), held in the NIH Brain Development Cohorts Data Sharing Platform. This is a multisite, longitudinal study designed to recruit approximately 7,000 families and follow them from pregnancy to early childhood.

The HBCD Study is supported by the NIH and additional federal partners under award numbers U01DA055352, U01DA055353, U01DA055366, U01DA055365, U01DA055362, U01DA055342, U01DA055360, U01DA055350, U01DA055338, U01DA055355, U01DA055363, U01DA055349, U01DA055361, U01DA055316, U01DA055344, U01DA055322, U01DA055369, U01DA055358, U01DA055371, U01DA055359, U01DA055354, U01DA055370, U01DA055347, U01DA055357, U01DA055367, U24DA055325, and U24DA055330. A full list of supporters is available at https://hbcdstudy.org/federal-partners/.

A full list of participating sites is available at: https://hbcdstudy.org/recruitment-sites/. HBCD Study Consortium investigators designed and implemented the study and/or provided data but did not necessarily participate in the analysis or writing of this report. This manuscript reflects the views of the authors and may not reflect the opinions or views of the NIH or the HBCD Study Consortium investigators.

Data used in the preparation of this article were obtained from the Adolescent Brain Cognitive DevelopmentSM (ABCD) Study (https://abcdstudy.org), held in the NIMH Data Archive (NDA). This is a multisite, longitudinal study designed to recruit more than 10,000 children age 9-10 and follow them over 10 years into early adulthood. The ABCD Study® is supported by the National Institutes of Health and additional federal partners under award numbers U01DA041048, U01DA050989, U01DA051016, U01DA041022, U01DA051018, U01DA051037, U01DA050987, U01DA041174, U01DA041106, U01DA041117, U01DA041028, U01DA041134, U01DA050988, U01DA051039, U01DA041156, U01DA041025, U01DA041120, U01DA051038, U01DA041148, U01DA041093, U01DA041089, U24DA041123, U24DA041147. A full list of supporters is available at https://abcdstudy.org/federal-partners.html. A listing of participating sites and a complete listing of the study investigators can be found at https://abcdstudy.org/consortium_members/. ABCD consortium investigators designed and implemented the study and/or provided data but did not necessarily participate in the analysis or writing of this report. This manuscript reflects the views of the authors and may not reflect the opinions or views of the NIH or ABCD consortium investigators.

The Bipolar & Schizophrenia Consortium for Parsing Intermediate Phenotypes (BSNIP1; 10.15154/tnzs-a323) dataset (R01MH078113-01, R01MH077852-01, R01MH077851-01, R01MH077945-01, R01MH077862-01) and its renewal (BSNIP2; 10.15154/8v3w-et72) (R01MH103368-01, R01MH103366-01) were collected with the aim to improve diagnosis and clinical management of psychosis by defining biologically based biotypes that better predict symptoms, course, and treatment response across major psychoses.

The Early Brain Development in Twins (EBDT) dataset was collected to understand the role of genetic and environmental contributions to brain structure and function in the crucial period of development that is early childhood (U01MH070890-11). (10.15154/1822-fs72)

The Age-ility Project (Phase 1) dataset was collected with the aim of developing novel analysis approaches that integrate information across multiple brain imaging modalities (e.g., dMRI, fMRI, electrophysiology) to build a more complete picture of how brain networks are structurally and functionally organized to support cognition.

The Developing Human Connectome Project (dHCP) is an open science study, funded by the European Research Council to obtain and disseminate Magnetic Resonance Imaging (MRI) data which map the brain’s structural and functional development across the period from 20 weeks gestational age to full term. By coupling advances in imaging with bespoke solutions developed for the fetal and neonatal population, principally but not exclusively solving the problems of subject motion, the dHCP captures the development of brain anatomy and connectivity at a systems level (10.15154/92vw-g837).

Data from the VUMC-ASD dataset are collected seeking to elucidate the neural mechanisms underlying atypical sensory processing in autism spectrum disorders (ASD), focusing on tactile and interoceptive sensitivity and their relationship to social and behavioral symptoms. By integrating behavioral, neurophysiological, and neuroimaging approaches, the studies aim to identify how alterations in thalamocortical and salience networks contribute to sensory hypo- and hyper-responsiveness in ASD, ultimately informing early identification and novel intervention strategies. Collection of these data were supported by K01MH090232, R21MH101321, and R01MH102272.

## Code Availability Statement

Code for out-of-sample data alignment, obtaining centile curves, and fitting the GAMLSS models is available in a containerized Docker image that is downloadable at https://zenodo.org/records/17561821. Instructions for running the Docker image can also be found at this link and in the supplementary information. The postprocessing pipeline to obtain the microstructural and macrostructural measurements from dMRI data is also available as a Docker image downloadable at: https://hub.docker.com/r/kimm58/wm_lifespan_processing. Instructions for running the Docker image are available at https://zenodo.org/records/17144461.

## Data Availability Statement

Data from the Alzheimer’s Disease Neuroimaging Initiative (ADNI) are available upon request from https://adni.loni.usc.edu/. Data from the AOMIC-PIOP1 dataset are freely available for download on OpenNeuro: https://openneuro.org/datasets/ds002785/versions/2.0.0. Data from the AOMIC-PIOP2 dataset are freely available for download on OpenNeuro: https://openneuro.org/datasets/ds002790/versions/2.0.0. Data from the AOMIC-ID1000 dataset are freely available for download on OpenNeuro: https://openneuro.org/datasets/ds003097/versions/1.2.1. Data from the Boston Adolescent Neuroimaging of Depression and Anxiety (BANDA) dataset are available upon request from https://www.humanconnectome.org/study/connectomes-related-anxiety-depression. Data from BIOCARD are available upon request after filling out a data use application: https://www.gaaindata.org/partner/BIOCARD. Data from the Baltimore Longitudinal Study of Aging (BLSA) are available upon request from https://www.blsa.nih.gov/. Data from the Calgary Preschool MRI Dataset are freely available for download at: https://doi.org/10.17605/OSF.IO/AXZ5R. Data from the Centre for Attention Learning and Memory (CALM) dataset are available upon request from https://calm.mrc-cbu.cam.ac.uk/researchers/. Data from the Cambridge Center for Ageing Neuroscience (CAMCAN) dataset are available upon request from https://camcan-archive.mrc-cbu.cam.ac.uk/dataaccess/. Data from the Dallas Lifespan Brain Study (DLBS) dataset are freely available for download on OpenNeuro: https://openneuro.org/datasets/ds004856/versions/1.2.0. Data from the Health & Aging Brain Study - Health Disparities (HABS-HD) dataset are available upon request from https://apps.unthsc.edu/itr/reports. Imaging data and basic demographic information for the Healthy Brain Network (HBN) dataset are freely available to download from https://fcon_1000.projects.nitrc.org/indi/cmi_healthy_brain_network/.

Phenotypic data are available upon request by filling out a data use agreement: https://fcon_1000.projects.nitrc.org/indi/cmi_healthy_brain_network/Phenotypic.html. Data from the Human Connectome Project – Aging (HCPA) dataset are available upon request from https://www.humanconnectome.org/study/hcp-lifespan-aging. Data from the Lifespan Baby Connectome Project (HCPBaby) dataset are available upon request https://www.humanconnectome.org/study/lifespan-baby-connectome-project. Data from the Human Connectome Project – Development (HCPD) dataset are available upon request from https://www.humanconnectome.org/study/hcp-lifespan-development. Data from the Human Connectome Project – Young Adult (HCP) dataset are freely available for download from https://www.humanconnectome.org/study/hcp-young-adult. Data from the Infant Brain Imaging Study (IBIS) are available for download from the National Institutes of Mental Health data archive upon request from https://nda.nih.gov/. Data from the International Consortium for Brain Mapping (ICBM) dataset are available upon request from www.loni.usc.edu/ICBM. Data from the Longitudinal Brain Correlates of Multisensory Lexical Processing in Children study (shortened to Lexical in this manuscript) are freely available for download on OpenNeuro: https://openneuro.org/datasets/ds001894/versions/1.4.2. Data from the Memory and Aging Project (MAP), Religious Orders Study (ROS), and the Minority Aging Research Study (MARS) datasets are available upon request from https://www.radc.rush.edu/. Data from the Multisite, Multiscanner, and Multisubject Acquisitions for Studying Variability in Diffusion Weighted Magnetic Resonance Imaging (MASiVar) dataset are freely available for download on OpenNeuro: https://openneuro.org/datasets/ds003416/versions/2.0.2. Data from the National Alzheimer’s Coordinating Center (NACC) and the Standardized Centralized Alzheimer’s & Related Dementias Neuroimaging (SCAN) are available upon request from https://naccdata.org/requesting-data/data-request-process. Imaging data and basic demographic information for the Nathan Kline Institute – Rockland Sample (NKI) dataset are freely available to download from https://fcon_1000.projects.nitrc.org/indi/enhanced/sharing_neuro.html.

Phenotypic data are available upon request by filling out a data use agreement: https://fcon_1000.projects.nitrc.org/indi/enhanced/sharing_phenotypic.html. Data from the Pediatric Imaging, Neurocognition, and Genetics dataset (PING) are available for download from the National Institutes of Mental Health data archive upon request from https://nda.nih.gov/. Data from the Queensland Twin Adolescent Brain (QTAB) dataset are freely available for download on OpenNeuro: https://openneuro.org/datasets/ds004146/versions/1.0.4. Data from the UCLA Consortium for Neuropsychiatric Phenomics LA5c Study (UCLA) dataset are freely available for download on OpenNeuro: https://openneuro.org/datasets/ds000030/versions/1.0.0. Data from the Southwest University (SWU) Longitudinal Imaging Multimodal dataset are freely available for download here: https://fcon_1000.projects.nitrc.org/indi/retro/southwestuni_qiu_index.html. Data from the Social Reward and Nonsocial Reward Processing Across the Adult Lifespan: An Interim Multi-echo fMRI and Diffusion Dataset (referred to as TempleSocial in this manuscript) are freely available for download on OpenNeuro: https://openneuro.org/datasets/ds005123/versions/1.1.3. Data from UK Biobank (UKBB) are available upon request from https://www.ukbiobank.ac.uk/.

Data from the UPennRisk dataset are freely available for download on OpenNeuro: https://openneuro.org/datasets/ds002843/versions/1.0.1. Data from the dataset titled “A longitudinal neuroimaging dataset on language processing in children ages 5, 7, and 9 years old” (referred to as UTAustin579 in this manuscript) are freely available for download on OpenNeuro: https://openneuro.org/datasets/ds003604/versions/1.0.7. Data from the Vanderbilt Memory and Aging Project (VMAP_JEFFERSON, VMAP_2.0, TN Aging Project) are available for download upon request from https://vmacdata.org/vmap/data-requests. Data from the Wisconsin Registry for Alzheimer’s Prevention (WRAP) are available upon request from https://wrap.wisc.edu/data-requests-2/. Data from the HEALthy Brain and Child Development (HBCD) Study are available for download upon request from (https://hbcdstudy.org/data-sharing/). Data from the Adolescent Brain Cognitive Development (ABCD) Study are available for download upon request from (https://abcdstudy.org/scientists/data-sharing/). Data used in this manuscript from the Ageility Project (Phase 1 only) are available for download from (https://www.nitrc.org/projects/age-ility). Data from the Early Brain Development in Twins (EBDT) dataset are available for download from the National Institutes of Mental Health data archive upon request from https://nda.nih.gov/<u>. Data from the Bipolar & Schizophrenia</u> <u>Consortium for Parsing Intermediate Phenotypes dataset (BSNIP1) and its renewal (BSNIP2)</u> are available for download from the National Institutes of Mental Health data archive upon request from https://nda.nih.gov/. Data from the Developing Human Connectome Project (dHCP) are available for download from the National Institutes of Mental Health data archive upon request from https://nda.nih.gov/. Vanderbilt University data (MORGAN, BABIES-ABC, CUTTING, VUMC-ASD) subject to third party restrictions. Please contact corresponding author for data requests.

## Ethics Declaration

**<u>T.J.H</u>**: In addition to having grants from the NIH, serves on the scientific advisory board for Vivid Genomics. Serves as a consultant for Circular Genomics. Serves as Deputy Editor of Alzheimer’s & Dementia: Translational Research and Clinical Intervention. Serves as Senior Associate Editor for Alzheimer’s and Dementia. **<u>S.M.R</u>**: Is an employee of the Intramural Research Program of the NIA/NIH. As such, the BLSA studies and investigators are funded by the NIA. **<u>K.L.H</u>**: Participated in the Federation of Associations in Behavioral & Brain Sciences Hill Day in May 2025, meeting with congressional representatives to advocate for continued National Science Foundation (NSF) funding as part of the Coalition for National Science Funding efforts celebrating 75 years of NSF support. **<u>A.L.J</u>**: In addition to having grants from the NIH, serves as a member of the advisory board for Lantheus - Diagnostic and Therapeutic Innovations, Billerica, MA. Serves as Chair of the Observational Study Monitoring Board, Diverse-VCID: White Matter Lesion Etiology of Dementia in Diverse Populations (Diverse VCID) Study, Bethesda, MD. Member of the Observational Study Monitoring Board, Determinants of Incident Stroke Cognitive Outcomes and Vascular Effects on Recovery (DISCOVERY) Study, Bethesda, MD. Member of the External Advisory Board, Kansas Alzheimer’s Disease Core Center (P30), Kansas City, KS. Member of the Scientific Advisory Committee for Paul B. Beeson Emerging Leaders Career Development Program, John A. Hartford Foundation and American Federations for Aging Research, New York, NY. Member of the External Advisory Committee for the Clin-STAR Coordinating Center, American Federation for Aging Research, New York, NY. Member of the Alzheimer’s Disease and Related Dementia Advisory Council, State of Tennessee. All other authors declare no competing interests.

## Contributions

M.E.K. performed the bulk of the data processing and organization, performed the quality control, conducted analyses, created figures, created the software for out-of-sample alignment and the associated Docker image, and wrote and edited the manuscript and supplementary information. K.G.S. provided a supervisory role in the direction of the manuscript, helped design the study, and provided substantial feedback on the manuscript. B.A.L. provided a supervisory role in the direction of the manuscript. C.G., K.R., P.K., N.R.N., G.R., and Z.L. helped with the processing and preprocessing of the data and figure design. S.N.V. and P.Z. provided direction on the design of the statistical analyses. All other authors, including those mentioned above, made substantial contributions to the conception or design of the work or data acquisition. All authors provided substantial revisions to the manuscript.

