## Supplemental Info for "White matter microstructure and macrostructure brain charts across the human lifespan"

[A] HABS-HD MPIs: Sid E O'Bryant, Kristine Yaffe, Arthur Toga, Robert Rissman, & Leigh Johnson; and the HABS-HD Investigators: Meredith Braskie, Kevin King, James R Hall, Melissa Petersen, Raymond Palmer, Robert Barber, Yonggang Shi, Fan Zhang, Rajesh Nandy, Roderick McColl, David Mason, Bradley Christian, Nicole Phillips, Stephanie Large, Joe Lee, Badri Vardarajan, Monica Rivera Mindt, Amrita Cheema, Lisa Barnes, Mark Mapstone, Annie Cohen, Amy Kind, Ozioma Okonkwo, Raul Vintimilla, Zhengyang Zhou, Michael Donohue, Rema Raman, Matthew Borzage, Michelle Mielke, Beau Ances, Ganesh Babulal, Jorge Llibre-Guerra, Carl Hill and Rocky Vig.

[B] Data used in preparation of this article were obtained from the Alzheimer's Disease Neuroimaging Initiative (ADNI) database (adni.loni.usc.edu). As such, the investigators within the ADNI contributed to the design and implementation of ADNI and/or provided data but did not participate in analysis or writing of this report. A complete listing of ADNI investigators can be found at: http://adni.loni.usc.edu/wp-content/uploads/how_to_apply/ADNI_Acknowledgement_List.pdf

[C] Data used in preparation of this article were derived from BIOCARD study, supported by grant U19 – AG033655 from the National Institute on Aging. The BIOCARD study team did not participate in the analysis or writing of this report, however, they contributed to the design and implementation of the study. A listing of BIOCARD investigators can be found on the BIOCARD website (on the 'BIOCARD Data Access Procedures' page, 'Acknowledgement Agreement' document).

* *Corresponding Author*

### Table of Contents

*A. Additional Brain Charts* ____ *3*

*B. Data Information* ____ *19*

*C. Additional Diagnostic Group Centile Deviations* ____ *24*

- i. Sample Size of Diagnostic Groups ____ 24
- ii. Centile Score Deviations of Other Diagnostic Groups ____ 26
- iii. Effect Sizes of Centile Score Deviations ____ 29

*D. Model Fitting* ____ *33*

- i. Number of fractional polynomial terms for $\mu$ and $\sigma$ ____ 33
- ii. Empirical Model Stability Analysis ____ 34
- iii. Voxel Size Effect on FA ____ 35
- iv. Batch Correction and Site Harmonization ____ 35
- v. Sex-related Effects on White Matter Brain Charts ____ 37
- vi. Maxima and Minima of White Matter Tract Trajectories ____ 39

*E. Comparison to Existing Reference Charts* ____ *39*

- i. White Matter Lifespan Modeling in the Literature ____ 39
- ii. Comparison to Bethlehem et al. ____ 41
- iii. Comparison to Zhu et al. ____ 42

*F. Data Quality* ____ *45*

*G. Anomaly Detection* ____ *48*

*H. Out-of-Sample Alignment* ____ *49*

- i. Stability of Out-of-Sample Alignment ____ 49
- ii. Alignment with Other Preprocessing Pipelines ____ 50
- iii. How to Perform Out-of-Sample Data Alignment ____ 51
- iv. How to obtain centile trajectories of features ____ 53

*I. Considerations for Cross-sectional vs. Longitudinal Brain Charts* ____ *54*

*J. References* ____ *54*

### A. **Additional Brain Charts**

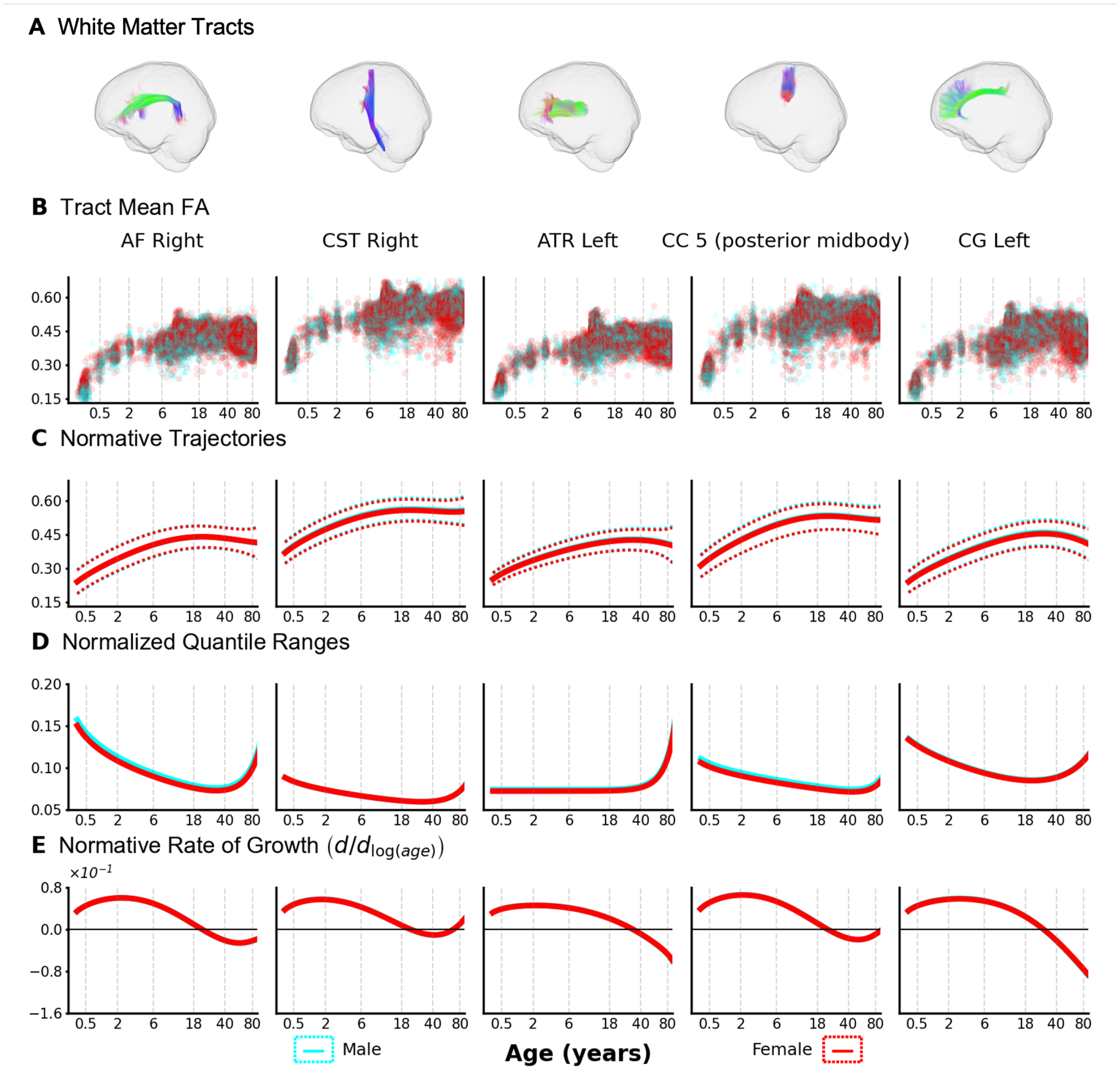


**Supplemental Figure SA.F1. Tract-specific Microstructural Brain Charts - Contralateral.** Lifespan brain charts for white matter microstructure (FA shown for contralateral tracts of Figure 2) reveal distinct developmental trajectories and variability across different WM pathways. A) Five exemplar tracts are shown (left to right): right arcuate fasciculus (AF Right); right corticospinal tract (CST Right); left anterior thalamic radiation (ATR Left); posterior midbody of the corpus callosum (CC 5 (posterior midbody)); and the left cingulate gyrus (CG Left). B) Raw FA data points for these tracts extend across the lifespan. C) Normative GAMLSS trajectories show FA increasing during development, plateauing in adulthood, and declining in later life, where the timing and magnitude vary by tract. Median (solid lines) and 2.5th/97.5th percentiles (dotted lines) are shown. D) Normalized quantile ranges indicate FA variability is generally lowest in middle age and increases later in life, though developmental variability patterns differ between tracts. E) The normative rate of change (first derivative) suggests that FA peaks at different ages and changes at different rates depending on the specific tract. Note: The x-axes (age in years) are log-scaled to emphasize developmental and aging periods.

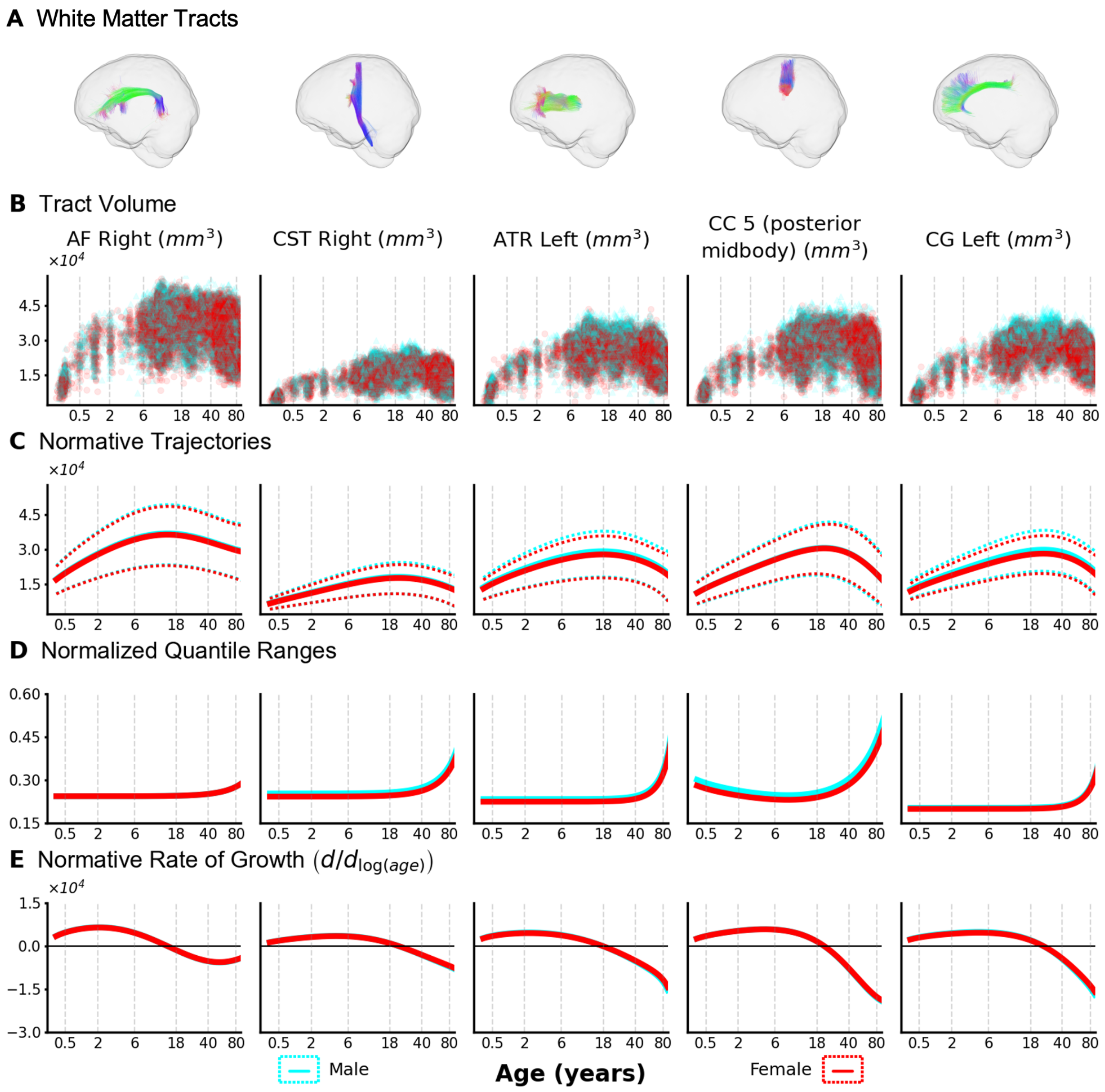


**Supplemental Figure SA.F2. Tract-specific Macrostructural Brain Charts - Contralateral.** Lifespan brain charts for white matter *macrostructure* (tract volume shown for contralateral tracts of Figure 3) reveal distinct developmental trajectories and variability across different WM pathways. A) Five exemplar tracts are shown (left to right): right arcuate fasciculus (AF Right); right corticospinal tract (CST Right); left anterior thalamic radiation (ATR Left); posterior midbody of the corpus callosum (CC 5 (posterior midbody)); and the left cingulate gyrus (CG Left). B) Raw tract volume data points for these tracts span the lifespan, illustrating differences in typical volume ranges between tracts. C) Normative GAMLSS trajectories show tract volume increasing during development, peaking in adolescence or early adulthood, and declining in later life, with tract-specific timing and magnitude. Median (solid lines) and 2.5th/97.5th percentiles (dotted lines) are shown. D) Normalized quantile ranges indicate that volume variability increases later in life and differs between sexes, being generally lowest in younger ages. E) The normative rate of change (first derivative) suggests that tract volume peaks at different ages and the rate of subsequent decline varies depending on the specific tract. Note: The x-axes (age in years) are log-scaled to emphasize developmental and aging periods.

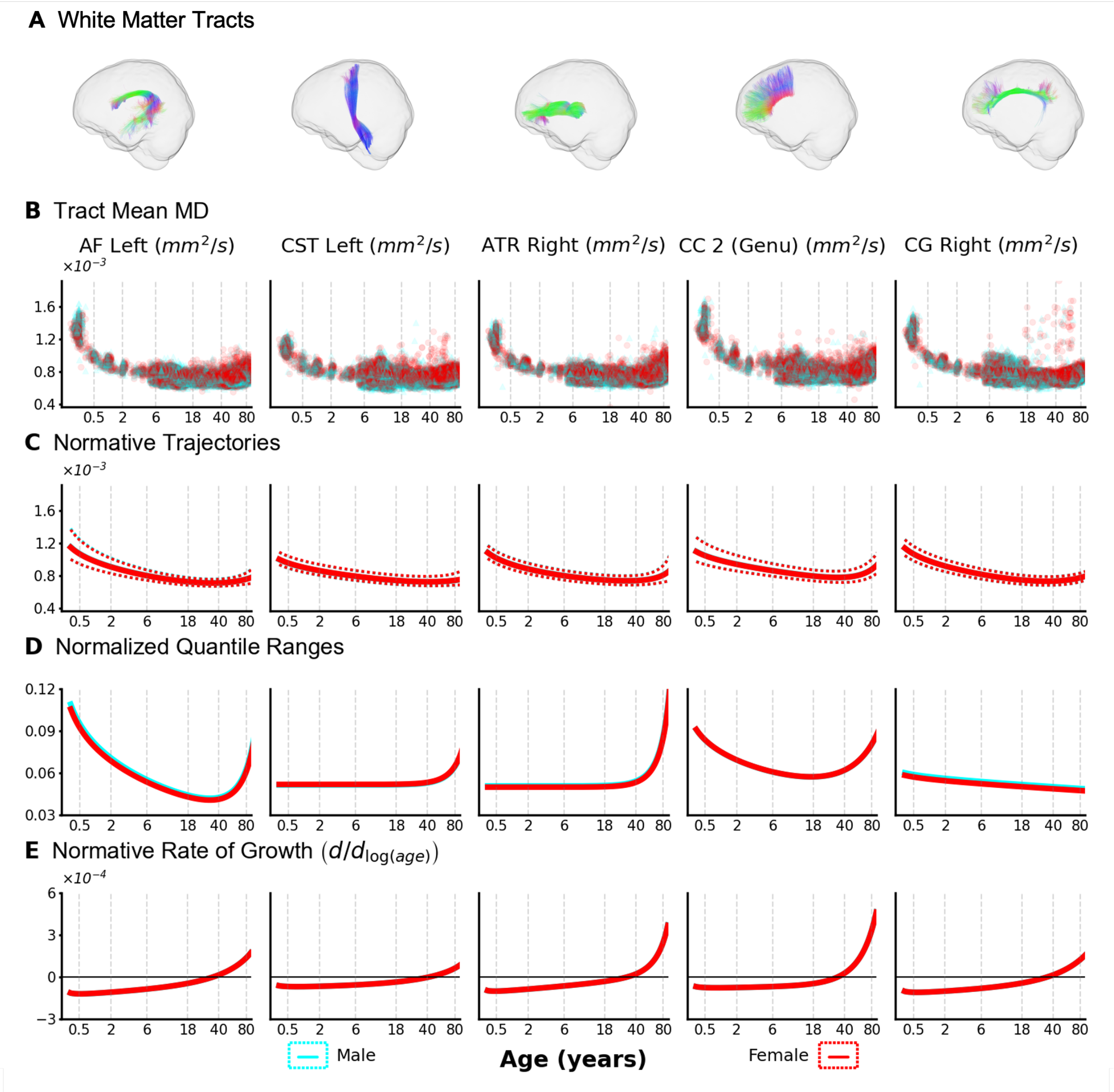


**Supplemental Figure SA.F3. MD microstructural brain charts.** Microstructural brain charts (MD) for five different WM tracts show tract-wise variability and varying developmental milestones across the human lifespan. A.) From left to right, left arcuate fasciculus; left corticospinal tract, right anterior thalamic radiation; genu of the corpus callosum; and the right cingulate gyrus. B.) Raw datapoints span the entirety of the lifespan at high density. C.) Most normative trajectories of MD for different pathways decrease in value during development, reach a minimum in adulthood, and then increase at different rates in aging. For all pathways represented, the male trajectories (blue) sit at nearly equal values to the female trajectories (red). D.) Normalized quantile differences indicate that microstructural variability in MD increases at the end of the human lifespan and is lowest in middle age. Variability in MD during development appears to be tract dependent, with some tracts showing lowest variability in infancy and others decreasing to a minimum in adulthood. E.) Rate of change of MD with respect to age for the represented tracts. MD of tracts reach minima at different ages in the lifespan, indicating tract-specific developmental patterns.

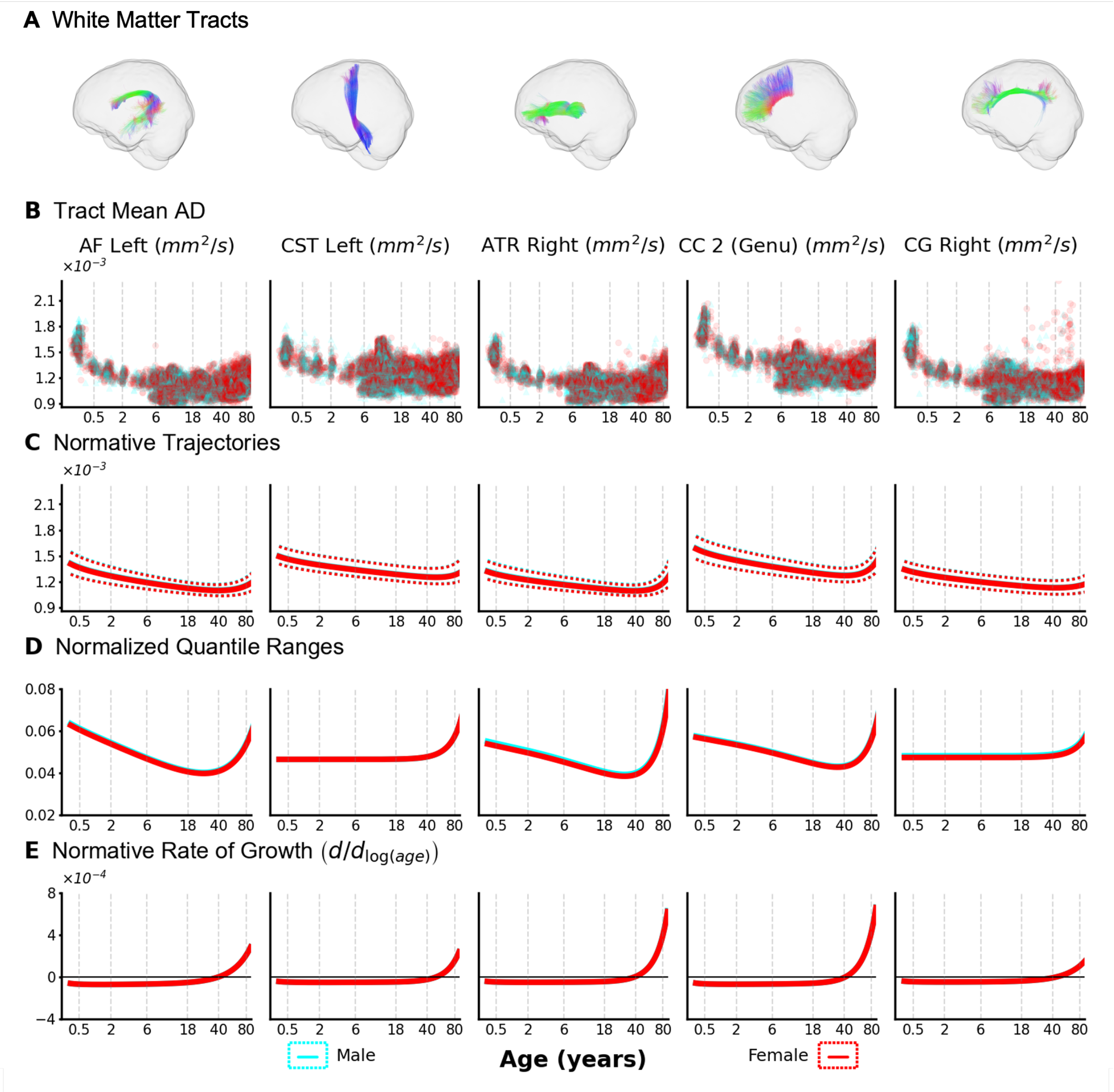


**Supplemental Figure SA.F4. AD microstructural brain charts.** Microstructural brain charts (AD) for five different WM tracts show tract-wise variability and varying developmental milestones across the human lifespan. A.) From left to right, left arcuate fasciculus; left corticospinal tract, right anterior thalamic radiation; genu of the corpus callosum; and the right cingulate gyrus. B.) Raw datapoints span the entirety of the lifespan at high density. C.) Normative trajectories of AD for different pathways decrease in value during development, reach a minimum in adulthood, and then increase at different rates in aging. For all pathways represented, the male trajectories (blue) sit slightly above the female trajectories (red). D.) Normalized quantile differences indicate that microstructural variability in AD increases at the end of the human lifespan and is lowest in middle age. Variability in AD during development appears to be tract dependent, with some tracts showing lowest variability in infancy and others decreasing to a minimum in adulthood. E.) Rate of change of AD with respect to age for the represented tracts. AD of tracts reach minima at different ages in the lifespan, indicating tract-specific developmental patterns.

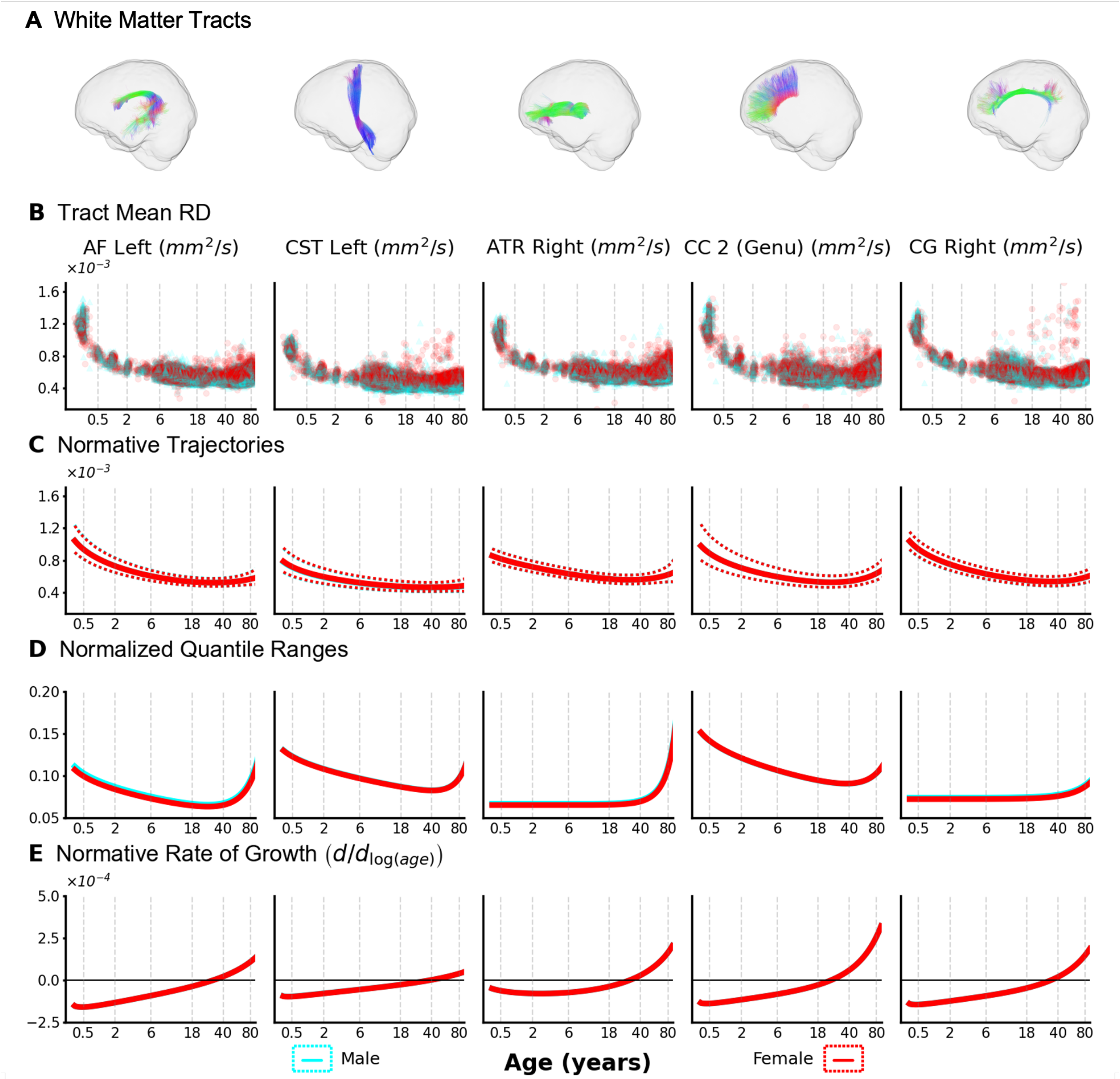


**Supplemental Figure SA.F5. RD microstructural brain charts.** Microstructural brain charts (RD) for five different WM tracts show tract-wise variability and varying developmental milestones across the human lifespan. A.) From left to right, left arcuate fasciculus; left corticospinal tract, right anterior thalamic radiation; genu of the corpus callosum; and the right cingulate gyrus. B.) Raw datapoints span the entirety of the lifespan at high density. C.) Normative trajectories of RD for different pathways typically decrease in value during development, reach a minimum in adulthood, and then increase at different rates in aging. For all pathways represented, the female trajectories (red) sit slightly above the male trajectories (blue). D.) Normalized quantile differences indicate that microstructural variability in RD increases at the end of the human lifespan and is lowest in middle age. RD variability appears to be tract dependent, with some tracts showing lowest variability in infancy and others increasing steadily across the lifespan. E.) Rate of change of RD with respect to age for the represented tracts. RD of tracts reach minima at different ages in the lifespan, indicating tract-specific developmental patterns.

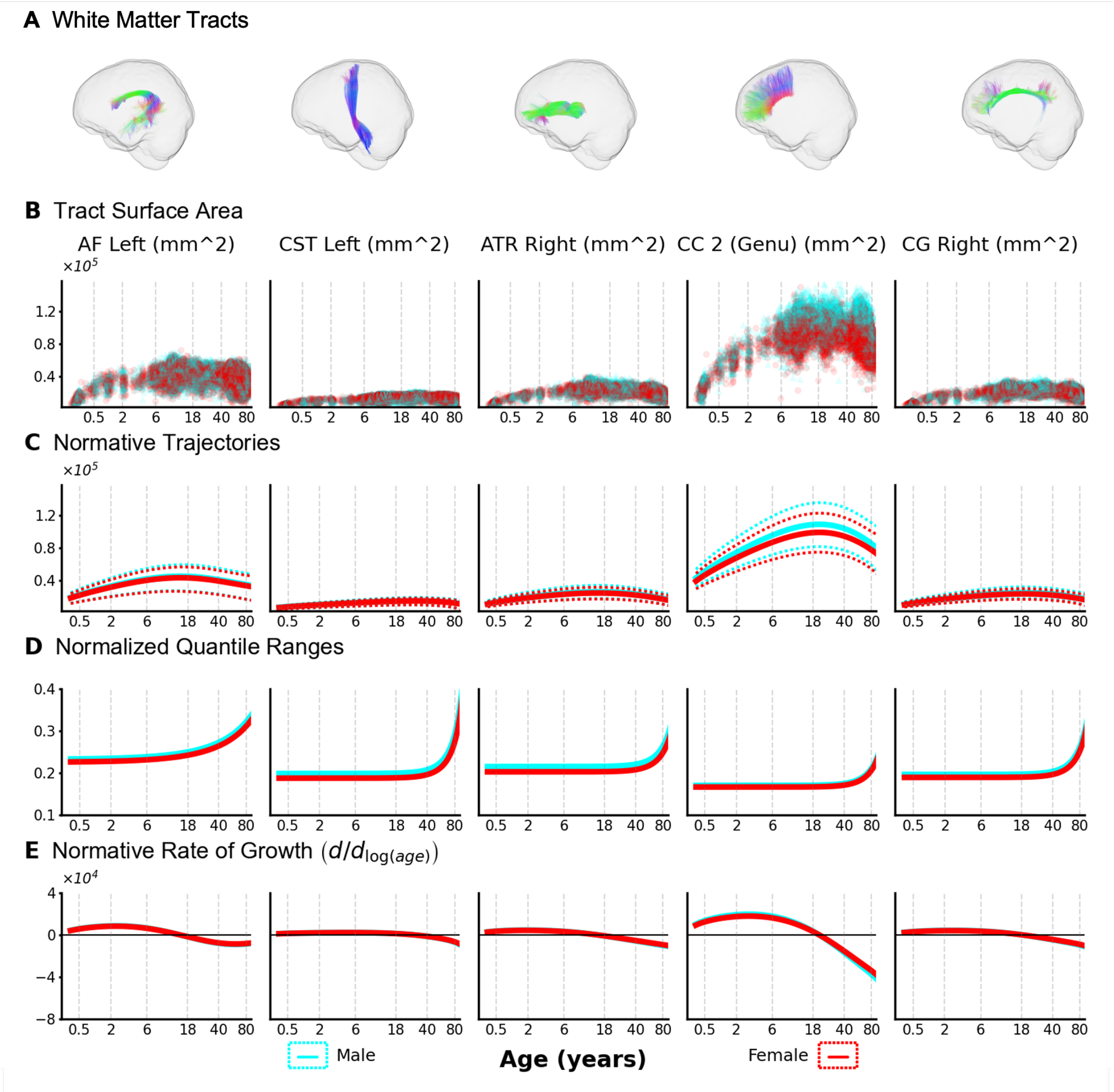


**Supplemental Figure SA.F6. Surface area macrostructural brain charts.** Macrostructural brain charts (surface area) for five different WM tracts show tract-wise variability and varying developmental milestones across the human lifespan. A.) From left to right, left arcuate fasciculus; left corticospinal tract, right anterior thalamic radiation; genu of the corpus callosum; and the right cingulate gyrus. B.) Raw datapoints span the entirety of the lifespan at high density. C.) Normative trajectories of surface area for different pathways increase in value during development, reach a maximum in adulthood, and then decrease at different rates in aging. For all pathways represented, the male trajectories (blue) sit slightly above the female trajectories (red). D.) Normalized quantile differences indicate that microstructural variability in tract surface area increases at the end of the human lifespan and is lowest in middle age. Variability in the male trajectories is consistently greater than that of the female trajectories. E.) Rate of change of surface area with respect to age for the represented tracts. Surface area of tracts reach maxima at different ages in the lifespan, indicating tract-specific developmental patterns.

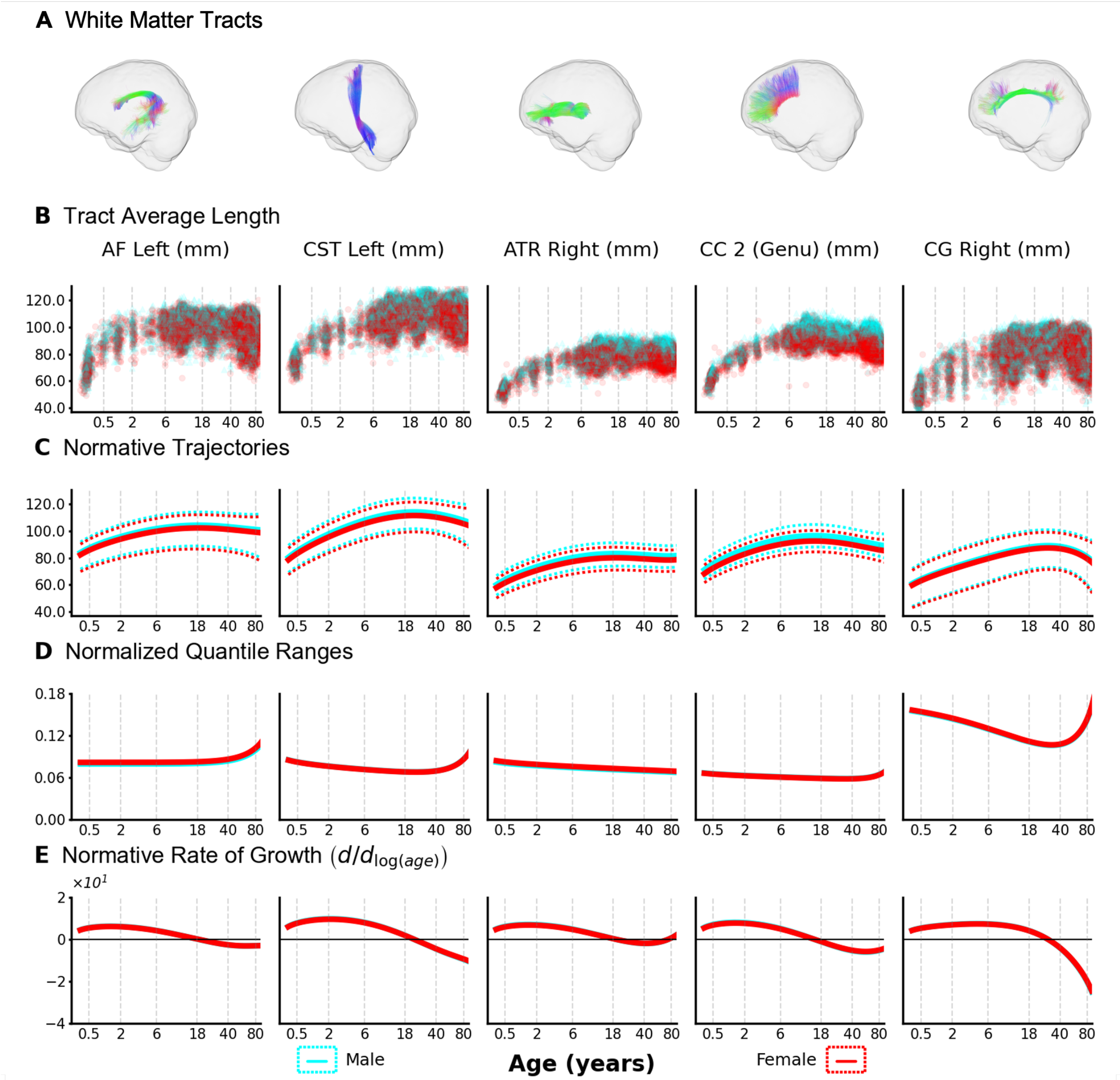


**Supplemental Figure SA.F7. Average length macrostructural brain charts.** Macrostructural brain charts (average length) for five different WM tracts show tract-wise variability and varying developmental milestones across the human lifespan. A.) From left to right, left arcuate fasciculus; left corticospinal tract, right anterior thalamic radiation; genu of the corpus callosum; and the right cingulate gyrus. B.) Raw datapoints span the entirety of the lifespan at high density. C.) Normative trajectories of average length for different represented pathways increase in value during development, reach a maximum in adulthood, and then decrease at different rates in aging. For all pathways represented, the male trajectories (blue) sit slightly above the female trajectories (red). D.) Normalized quantile differences indicate that macrostructural variability is very tract-dependent. Some tracts have the highest variability at the beginning of the lifespan and lowest at the end, whereas others have the opposite trend. The change in variability of average length across the lifespan also appears to be tract dependent, with some tracts decreasing somewhat consistently and others showing sharp increases. E.) Rate of change of average length with respect to age for the represented tracts. Average length of tracts reach maxima at different ages in the lifespan, indicating tract-specific developmental patterns.

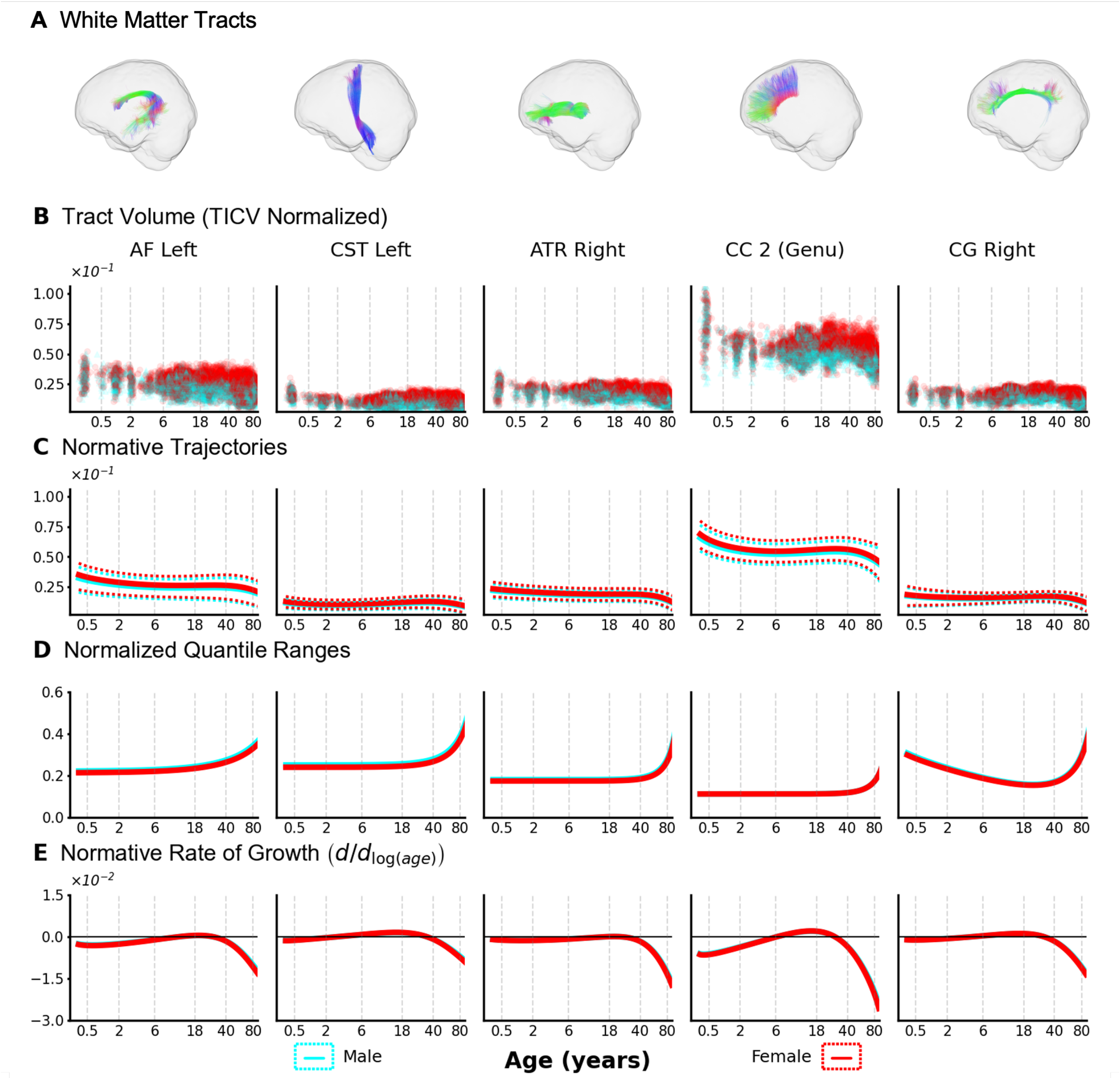


**Supplemental Figure SA.F8. Charts for tract volume normalized by TICV.** Lifespan trajectories of WM tract volume normalized by estimated total intracranial vault volume show most tracts decrease in relative volume in infancy, stabilize in middle age, and then decrease again in late life. A.) From left to right, left arcuate fasciculus; left corticospinal tract, right anterior thalamic radiation; genu of the corpus callosum; and the right cingulate gyrus. B.) Raw data points indicate that TICV-normalized volumes are higher in females than males, with this trend also appearing in C.) the lifespan trajectories for these measures. D.) Additionally, the normalized quantile ranges indicate that variability for TICV-normalized tract volume is lowest at the beginning of the lifespan and monotonically increases throughout life, with a very sharp increase in older age. E.) Like most other features, change is most rapid in infancy and decreases in magnitude in later age ranges.

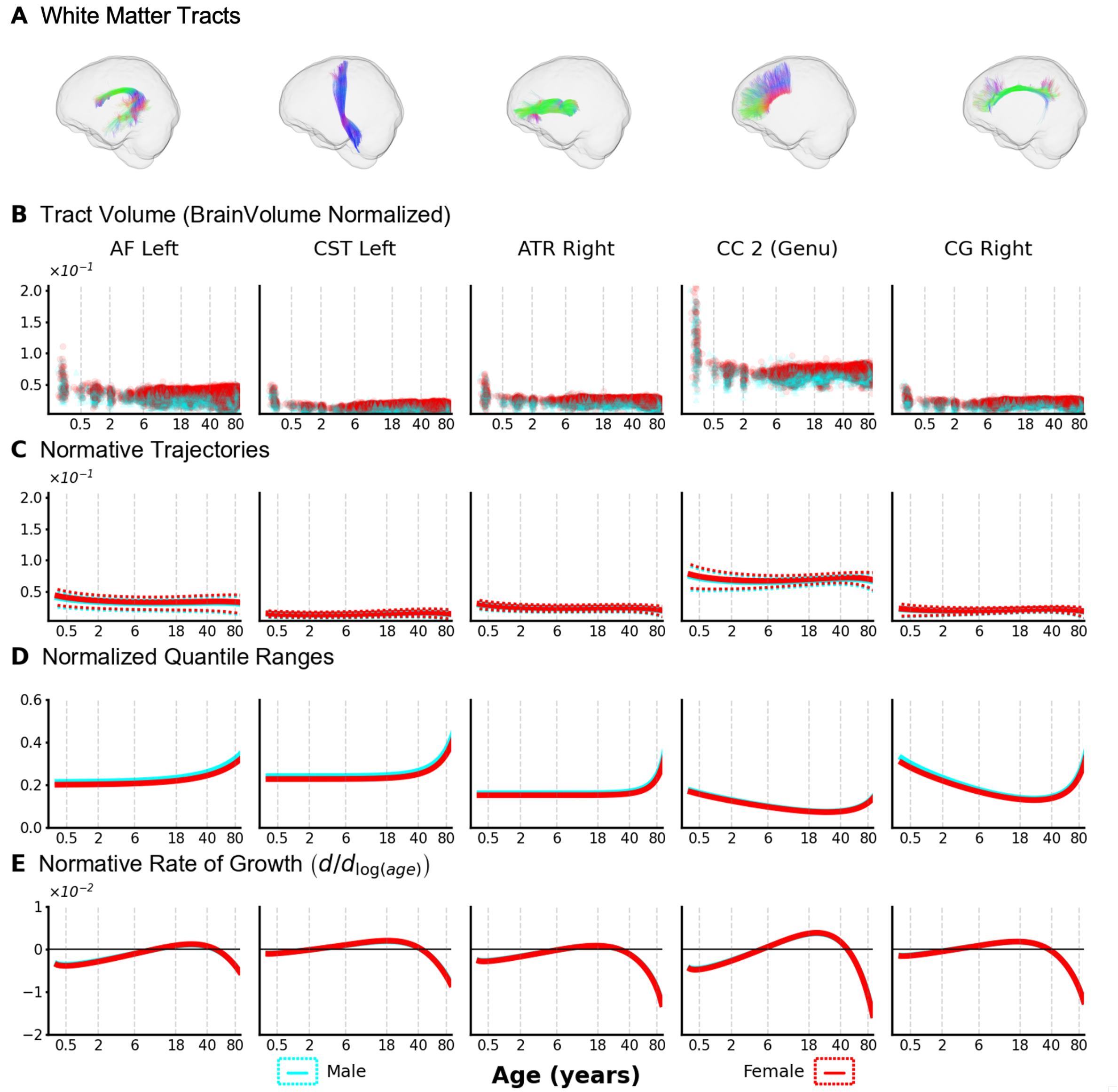


**Supplemental Figure SA.F9. Charts for tract volume normalized by brain volume without ventricles.** Lifespan trajectories of WM tract volume normalized by brain volume without ventricles show most tracts decrease in relative volume in infancy, stabilize or increase in middle age, and then decrease again in late life. A.) From left to right, left arcuate fasciculus; left corticospinal tract, right anterior thalamic radiation; genu of the corpus callosum; and the right cingulate gyrus. B.) Raw data points indicate that tract volumes normalized by brain volume without ventricles are higher in females than males, with this trend also appearing in C.) the lifespan trajectories for these measures. D.) Additionally, the normalized quantile ranges indicate that variability for WM tract volume normalized by brain volume without ventricles is similar compared to the non-normalized counterparts. E.) Like most other features, change is most rapid in infancy and decreases in magnitude in later age ranges.

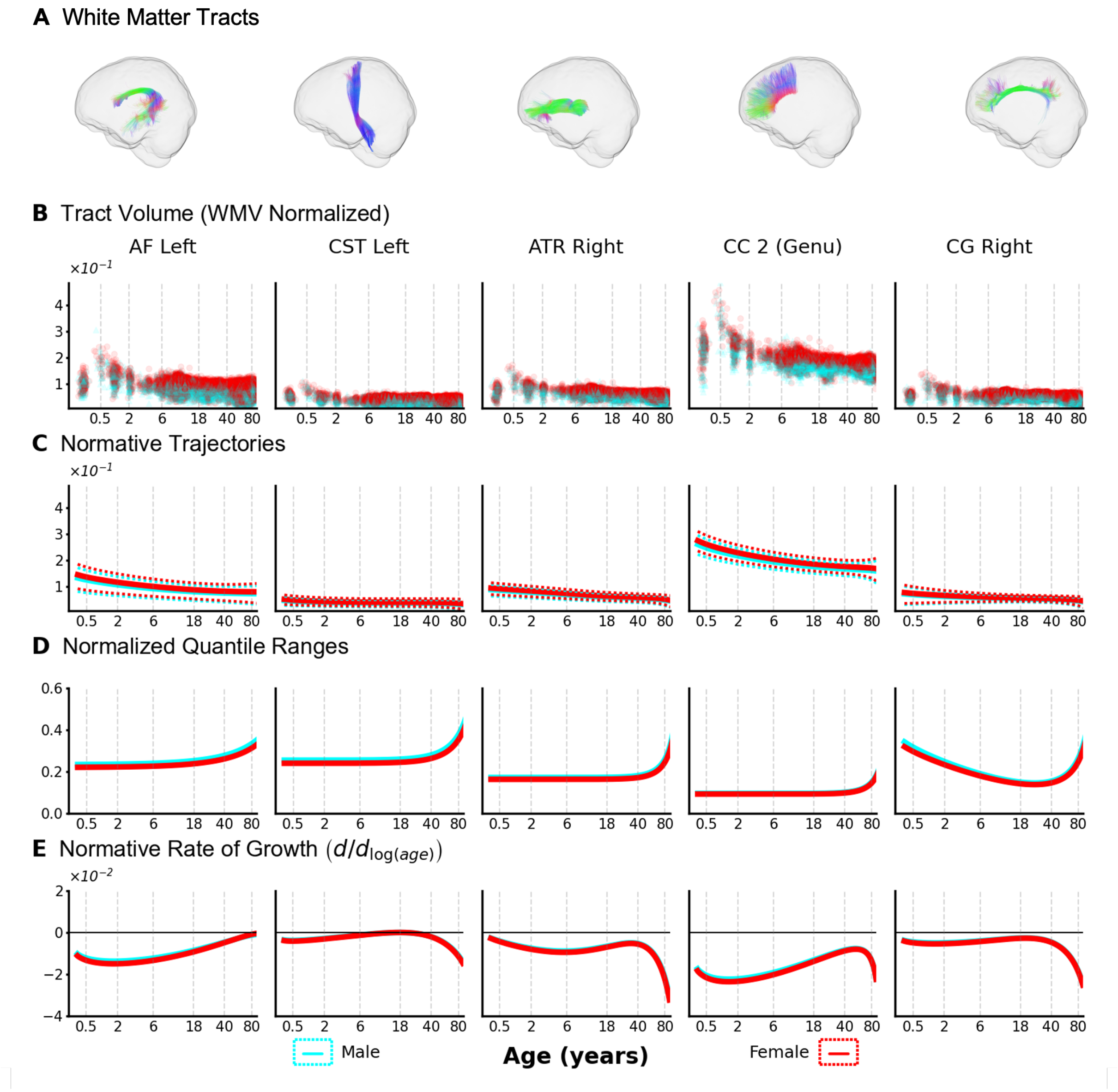


**Supplemental Figure SA.F10. Charts for tract volume normalized by cerebral WM volume.** Lifespan trajectories of WM tract volume normalized by cerebral WM volume show most tracts decrease in relative volume in infancy and then continue to decrease at lower rates throughout the rest of the lifespan. A.) From left to right, left arcuate fasciculus; left corticospinal tract, right anterior thalamic radiation; genu of the corpus callosum; and the right cingulate gyrus. B.) Raw data points indicate that tract volumes normalized by cerebral WM volume are higher in females than males, with this trend also appearing in C.) the lifespan trajectories for these measures. D.) The normalized quantile ranges indicate that variability is higher in males than females throughout the lifespan. Further, variability at the beginning of the lifespan is higher for some tracts when compared to the non-normalized counterparts. E.) Like most other features, change is most rapid in infancy and decreases in magnitude in later age ranges.

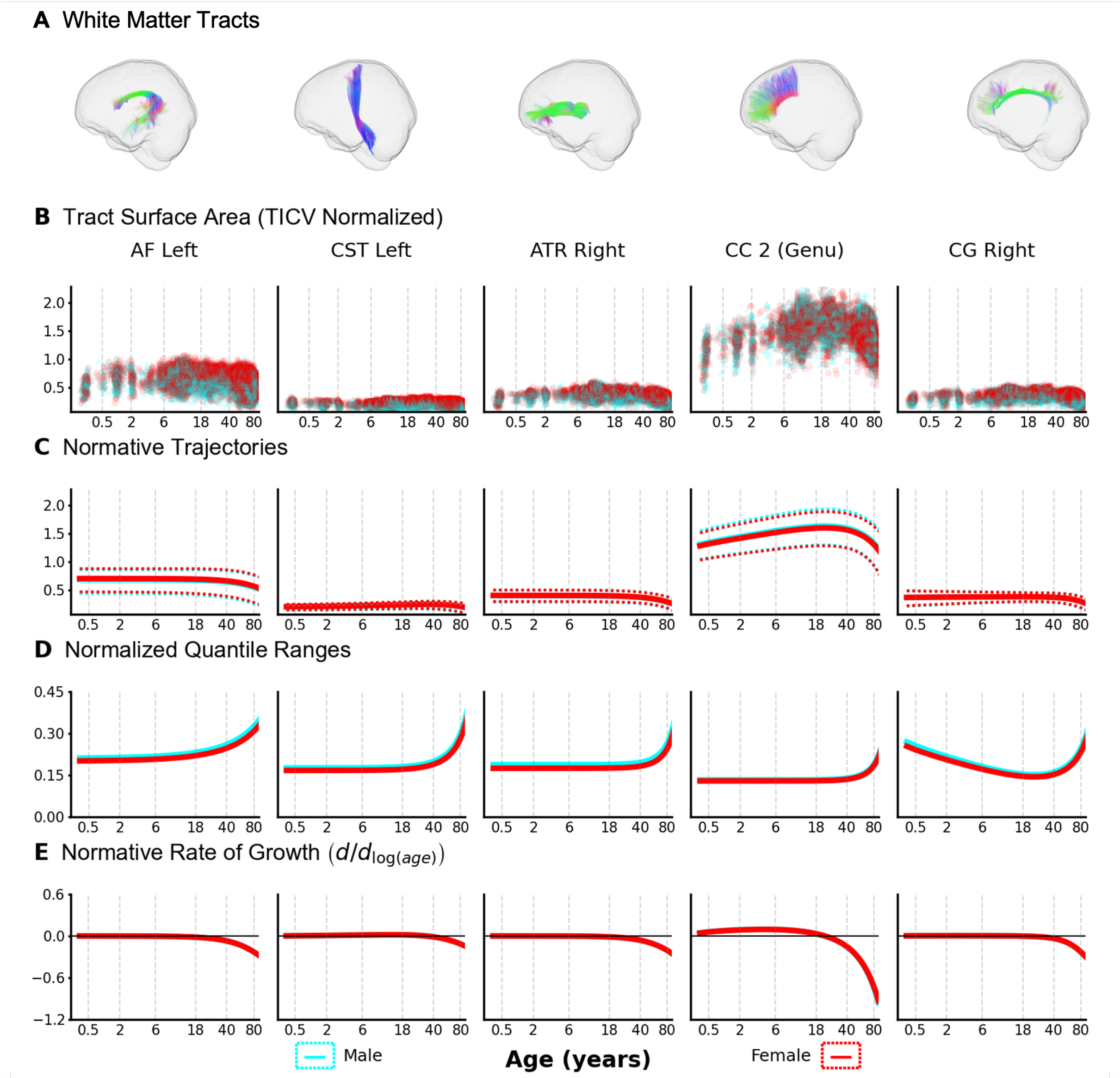


**Supplemental Figure SA.F11. Charts for surface area normalized by TICV.** Lifespan trajectories of WM surface area normalized by surface area derived from estimated total intracranial vault volume show tracts to have a small to zero increase in the ratio across most of the lifespan, with decreases in older age. A.) From left to right, left arcuate fasciculus; left corticospinal tract, right anterior thalamic radiation; genu of the corpus callosum; and the right cingulate gyrus. B.) Raw data points indicate that sex-specific trends in tract surface areas normalized by surface area derived from estimated total intracranial vault volume are tract specific, with some tracts having larger ratios in males and vice versa. This trend also is apparent in C.) the lifespan trajectories for these measures. D.) The normalized quantile ranges indicate that variability is higher in males than females throughout the lifespan. Further, variability is similar for tracts when compared to the non-normalized counterparts, with slight decreases for the normalized trajectories. E.) Unlike most other features, the values do not change across most of the lifespan, however, they do decrease towards the end of the lifespan like most other macrostructural features.

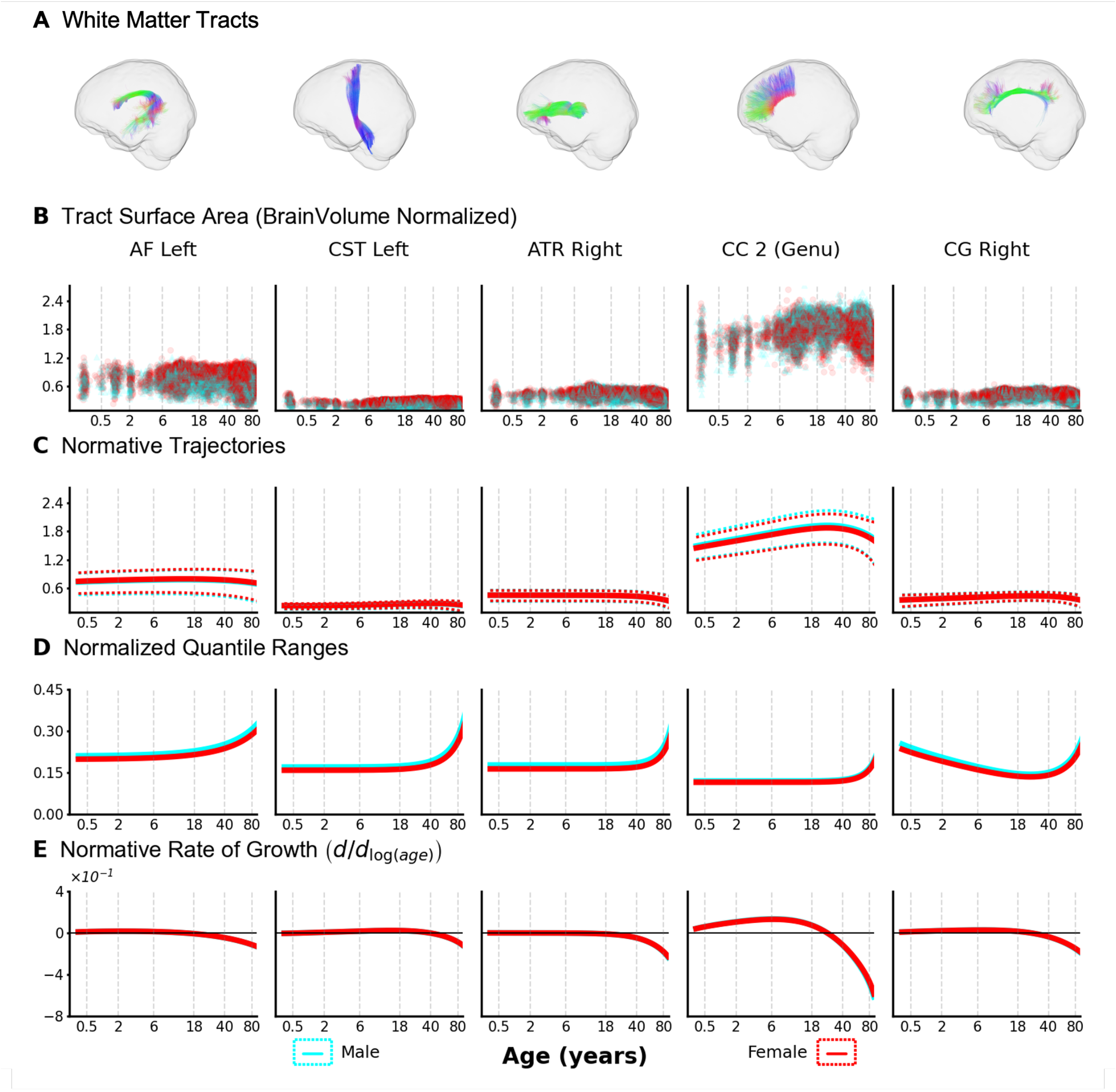


**Supplemental Figure SA.F12. Charts for surface area normalized by brain volume without ventricles.** Lifespan trajectories of WM surface area normalized by surface area derived from brain volume without ventricles show tracts to have a small to zero increase in the ratio across most of the lifespan, with decreases in older age. A.) From left to right, left arcuate fasciculus; left corticospinal tract, right anterior thalamic radiation; genu of the corpus callosum; and the right cingulate gyrus. B.) Raw data points indicate that sex-specific trends in tract surface areas normalized by surface area derived from brain volume without ventricles are tract specific, with some tracts having larger ratios in males and vice versa. This trend also is apparent in C.) the lifespan trajectories for these measures. D.) The normalized quantile ranges indicate that variability is higher in males than females throughout the lifespan. Further, variability is similar for tracts when compared to the non-normalized counterparts, with slight decreases for the normalized trajectories. E.) Unlike most other features, the values do not change across most of the lifespan, however, they do decrease towards the end of the lifespan like most other macrostructural features.

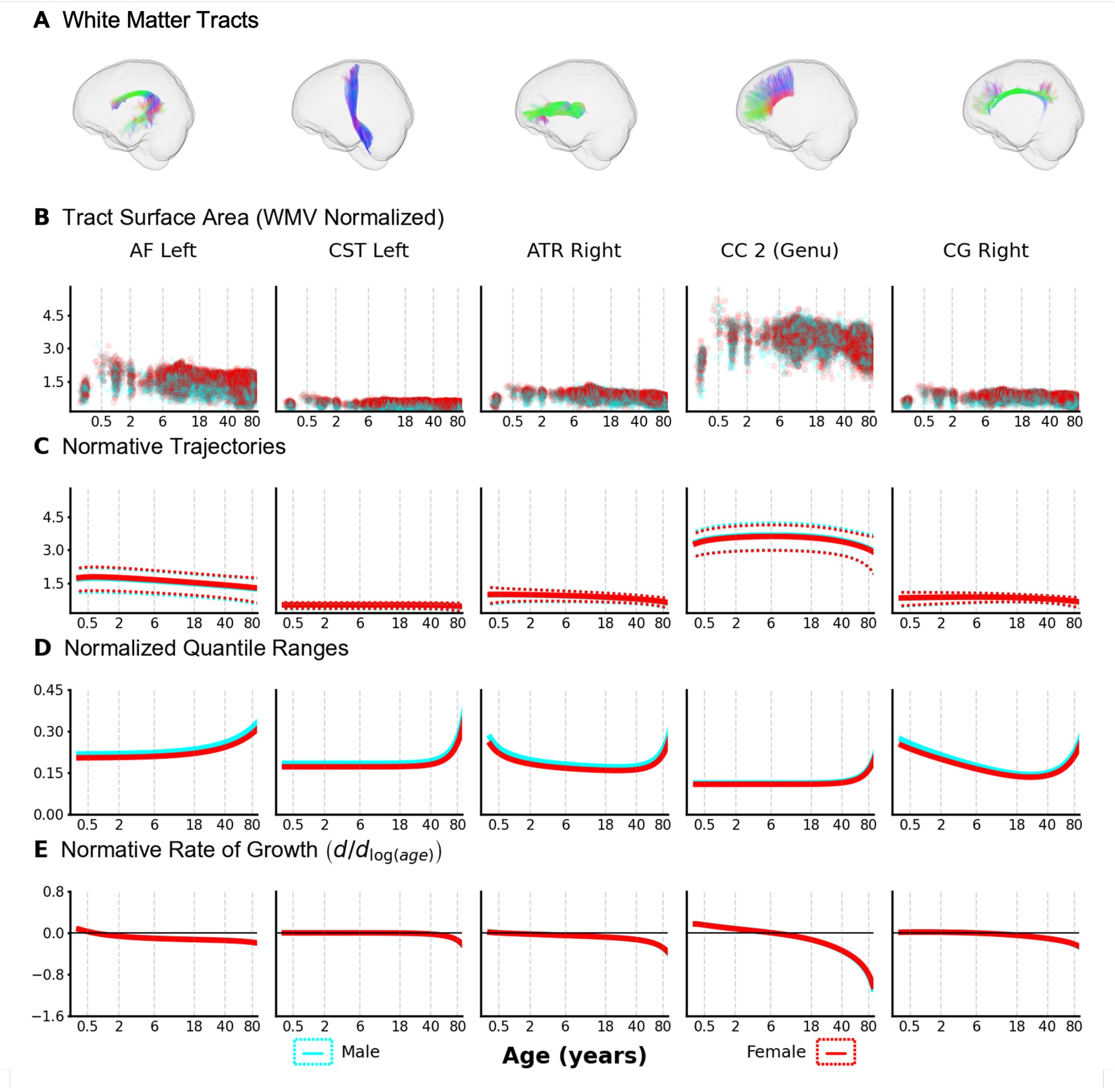


**Supplemental Figure SA.F13. Charts for surface area normalized by cerebral WM volume.** Lifespan trajectories of WM surface area normalized by surface area derived from cerebral WM volume show tracts to have a small decrease in the ratio across most of the lifespan, with larger decreases in older age. A.) From left to right, left arcuate fasciculus; left corticospinal tract, right anterior thalamic radiation; genu of the corpus callosum; and the right cingulate gyrus. B.) Raw data points indicate that sex-specific trends in tract surface areas normalized by surface area derived from cerebral WM volume are tract specific, with some tracts having larger ratios in males and vice versa. This trend also is apparent in C.) the lifespan trajectories for these measures. D.) The normalized quantile ranges indicate that variability is slightly higher in males than females throughout the lifespan. Further, variability is similar for tracts when compared to the non-normalized counterparts. E.) Unlike most other features, the values decrease across the entire lifespan for most tracts.

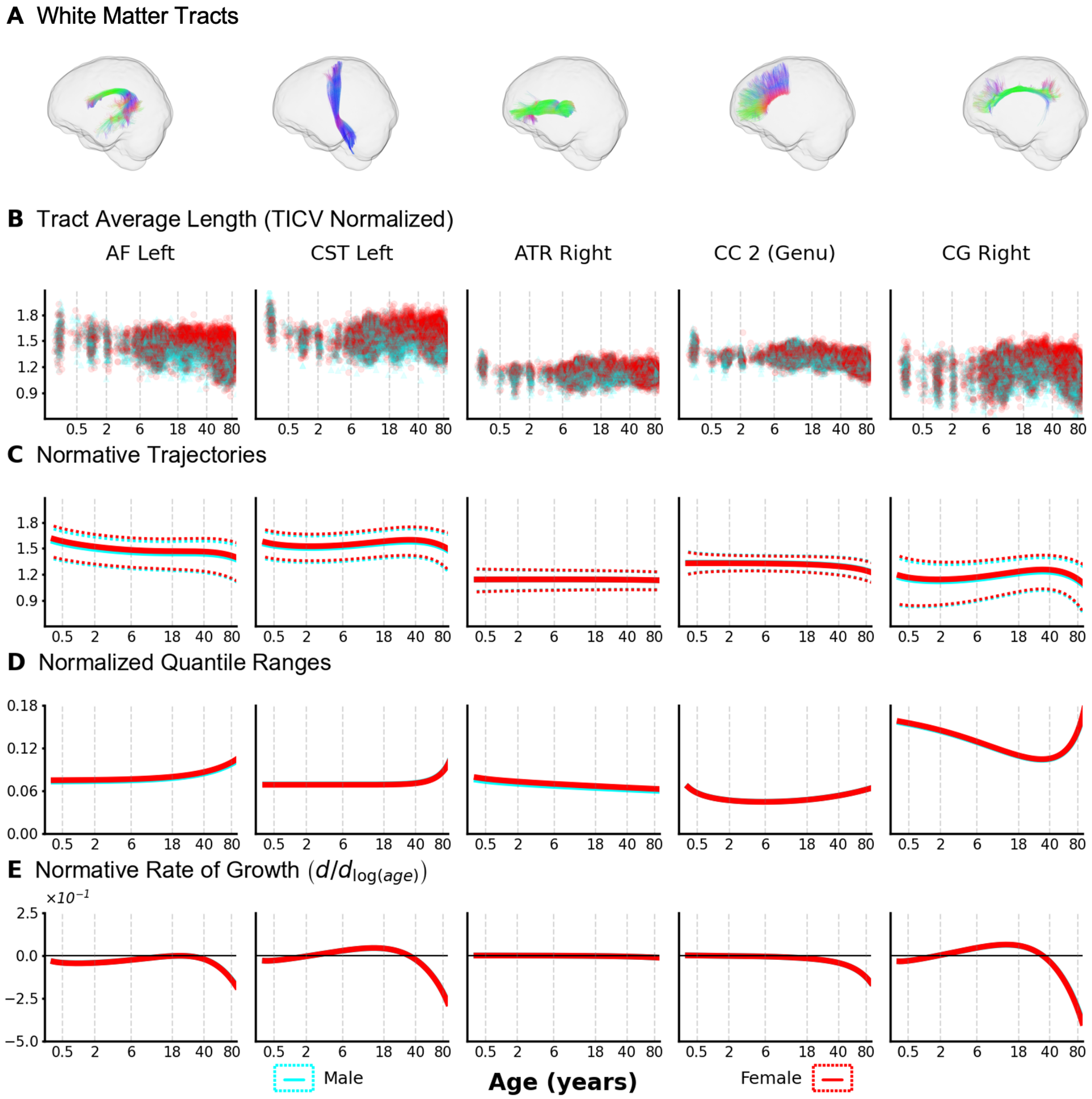


**Supplemental Figure SA.F14. Charts for average tract length normalized by TICV.** Lifespan trajectories of average tract length normalized by radius derived from estimated total intracranial volume (TICV) show tracts to be highly variable with increasing age. A.) From left to right, left arcuate fasciculus; left corticospinal tract, right anterior thalamic radiation; genu of the corpus callosum; and the right cingulate gyrus. B.) Raw data points indicate that tract average lengths normalized by radius derived from estimated total intracranial volume are slightly larger in females than males. This trend also is apparent in C.) the lifespan trajectories for these measures. D.) The normalized quantile ranges indicate that variability is similar between males and females throughout the lifespan. Further, variability is similar for tracts when compared to the non-normalized counterparts. E.) Unlike non-normalized average tract length, the values for TICV radius-normalized average tract length do not have sharp increases at the beginning of the lifespan.

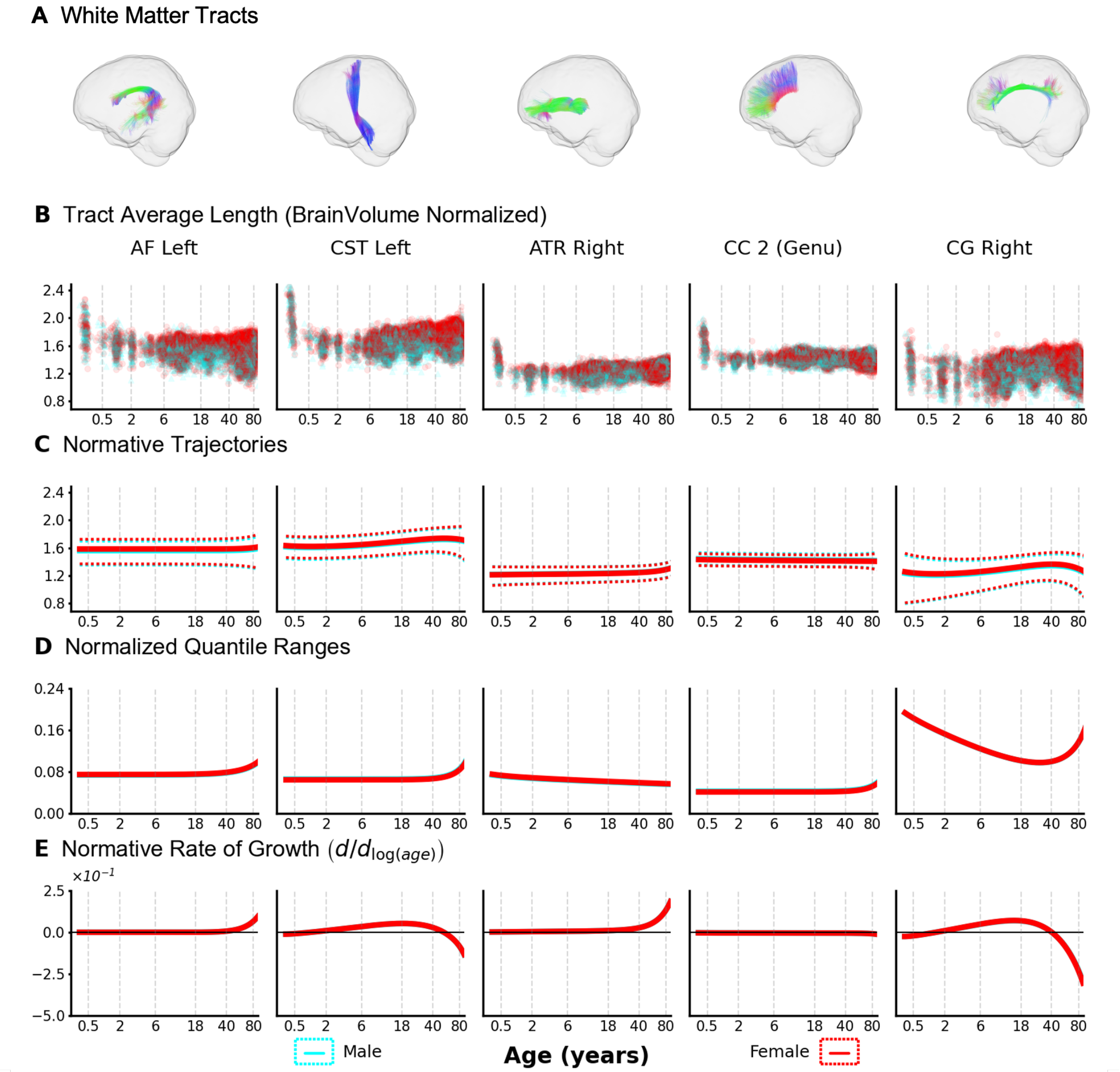


**Supplemental Figure SA.F15. Charts for average tract length normalized by brain volume without ventricles.** Lifespan trajectories of average tract length normalized by radius derived from brain volume without ventricles show tracts to be highly variable with increasing age. A.) From left to right, left arcuate fasciculus; left corticospinal tract, right anterior thalamic radiation; genu of the corpus callosum; and the right cingulate gyrus. B.) Raw data points indicate that tract average lengths normalized by radius derived from brain volume without ventricles are similar for males and females. This trend also is apparent in C.) the lifespan trajectories for these measures. D.) The normalized quantile ranges indicate that variability is similar between males and females throughout the lifespan. Further, variability is similar for tracts when compared to the non-normalized counterparts. E.) Unlike non-normalized average tract length, the values for average tract length normalized by radius from brain volume without ventricles increase at the end of the lifespan for some tracts.

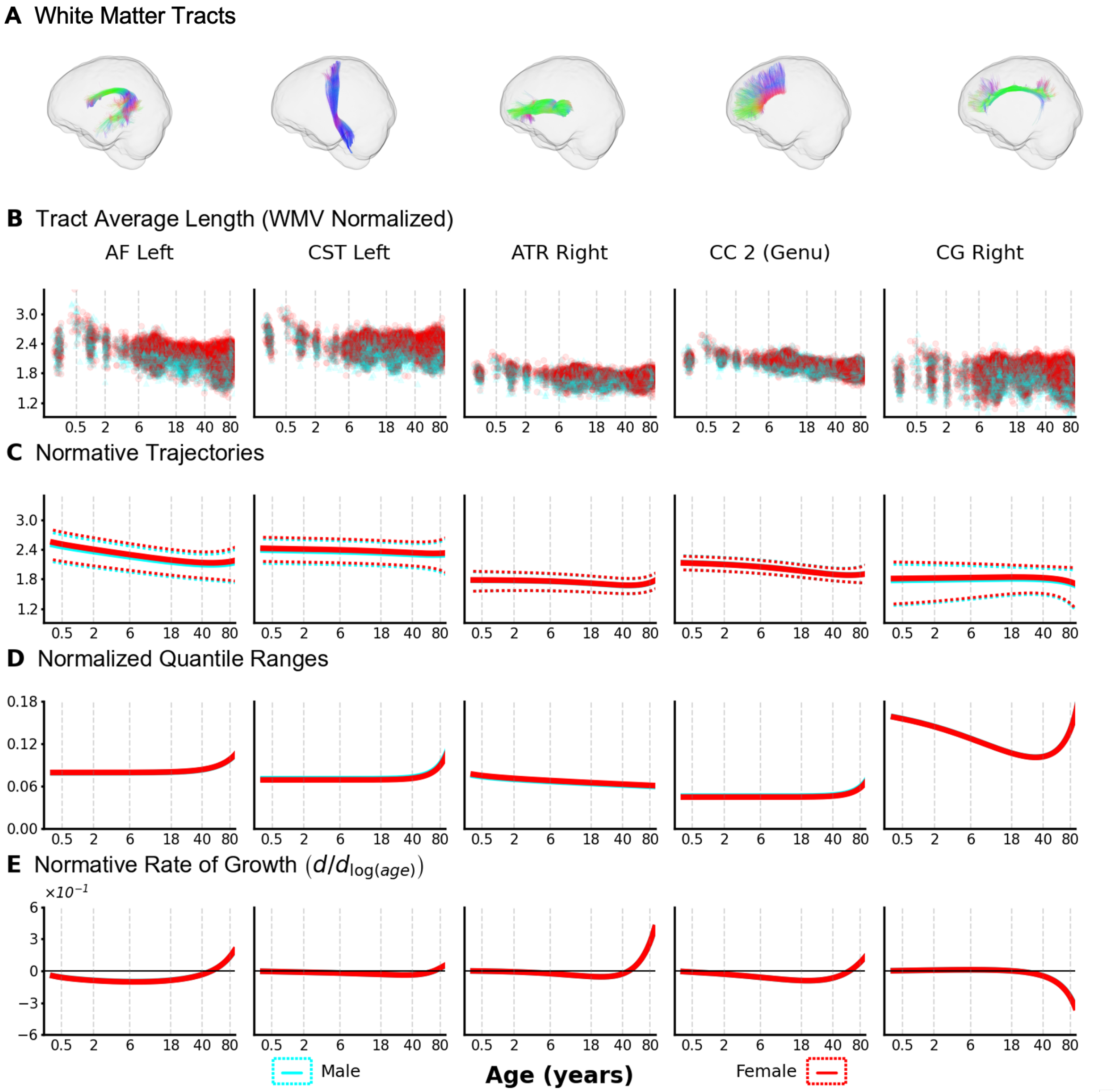


**Supplemental Figure SA.F16. Charts for average tract length normalized by cerebral WM volume.** Lifespan trajectories of average tract length normalized by radius derived from cerebral WM volume show tracts to mostly decrease with increasing age. A.) From left to right, left arcuate fasciculus; left corticospinal tract, right anterior thalamic radiation; genu of the corpus callosum; and the right cingulate gyrus. B.) Raw data points indicate that tract average lengths normalized by radius derived from cerebral WM volume are similar for males and females, with female trajectories sitting slightly higher for some tracts. This trend also is apparent in C.) the lifespan trajectories for these measures. D.) The normalized quantile ranges indicate that variability is similar between males and females throughout the lifespan. Further, variability is similar for tracts when compared to the non-normalized counterparts. E.) Unlike non-normalized average tract length, the values for average tract length normalized by radius from brain volume without ventricles increase at the end of the lifespan for some tracts.

### B. **Data Information**

**Supplemental Table SB.T1.** Participants from each dataset used to create the WM brain charts.

| Dataset | Scans | Participants | Participants After Quality Control | Neurotypical |
|---|---|---|---|---|
| ABCD | 20951 | 10607 | 10311 | 9000 |
| ADNI | 3451 | 1292 | 1229 | 584 |
| AOMIC PIOP1 | 211 | 211 | 204 | 204 |
| AOMIC PIOP2 | 226 | 226 | 224 | 224 |
| AOMIC ID1000 | 925 | 925 | 858 | 858 |
| Ageility | 131 | 131 | 131 | 131 |
| BANDA | 207 | 207 | 207 | 63 |
| BIOCARD | 974 | 210 | 186 | 153 |
| BLSA | 5423 | 1035 | 995 | 942 |
| BSNIP1 | 1400 | 594 | 554 | 197 |
| BSNIP2 | 1432 | 730 | 718 | 233 |
| Calgary | 249 | 88 | 88 | 88 |
| CALM | 314 | 314 | 313 | 189 |
| CAMCAN | 641 | 641 | 613 | 613 |
| VUMC-ASD | 173 | 164 | 132 | 63 |
| CUTTING | 1661 | 638 | 632 | 597 |
| dHCP | 753 | 668 | 285 | 285 |
| DLBS | 955 | 464 | 444 | 444 |
| EBDT | 1389 | 468 | 455 | 455 |
| HABSHD | 3767 | 2762 | 2558 | 1885 |
| HBCD | 546 | 500 | 285 | 285 |
| HBN | 2760 | 2754 | 2519 | 1049 |
| HCPA | 719 | 719 | 718 | 718 |
| HCPBaby | 411 | 211 | 197 | 197 |
| HCPD | 621 | 621 | 621 | 621 |
| HCP | 1109 | 1065 | 1064 | 1064 |
| ABC-Babies | 180 | 142 | 85 | 85 |
| IBIS | 494 | 272 | 217 | 167 |
| ICBM | 264 | 192 | 185 | 185 |
| Lexical | 113 | 113 | 112 | 112 |
| MAP | 1579 | 589 | 393 | 273 |
| MARS | 347 | 184 | 172 | 143 |
| MASiVar | 118 | 83 | 83 | 83 |
| MORGAN | 414 | 256 | 254 | 110 |
| NACC | 1768 | 1353 | 677 | 485 |
| NKI | 2224 | 1289 | 1205 | 840 |
| PING | 762 | 762 | 695 | 695 |

| | | | | |
|---|---|---|---|---|
| QTAB | 717 | 415 | 414 | 414 |
| ROS | 127 | 77 | 46 | 35 |
| SCAN | 303 | 288 | 260 | 188 |
| SWU | 460 | 234 | 231 | 231 |
| TempleSocial | 108 | 108 | 103 | 103 |
| UCLA | 262 | 262 | 248 | 116 |
| UKBB | 10425 | 10425 | 8509 | 8509 |
| UPennRisk | 251 | 152 | 151 | 151 |
| UTAustin579 | 376 | 239 | 237 | 237 |
| VMAP_2.0 | 342 | 327 | 266 | 266 |
| VMAP | 1239 | 328 | 310 | 160 |
| TN Alz Project | 194 | 168 | 159 | 58 |
| WRAP | 570 | 343 | 338 | 332 |
| ***Total*** | ***75036*** | ***46846*** | ***41891*** | ***35120*** |

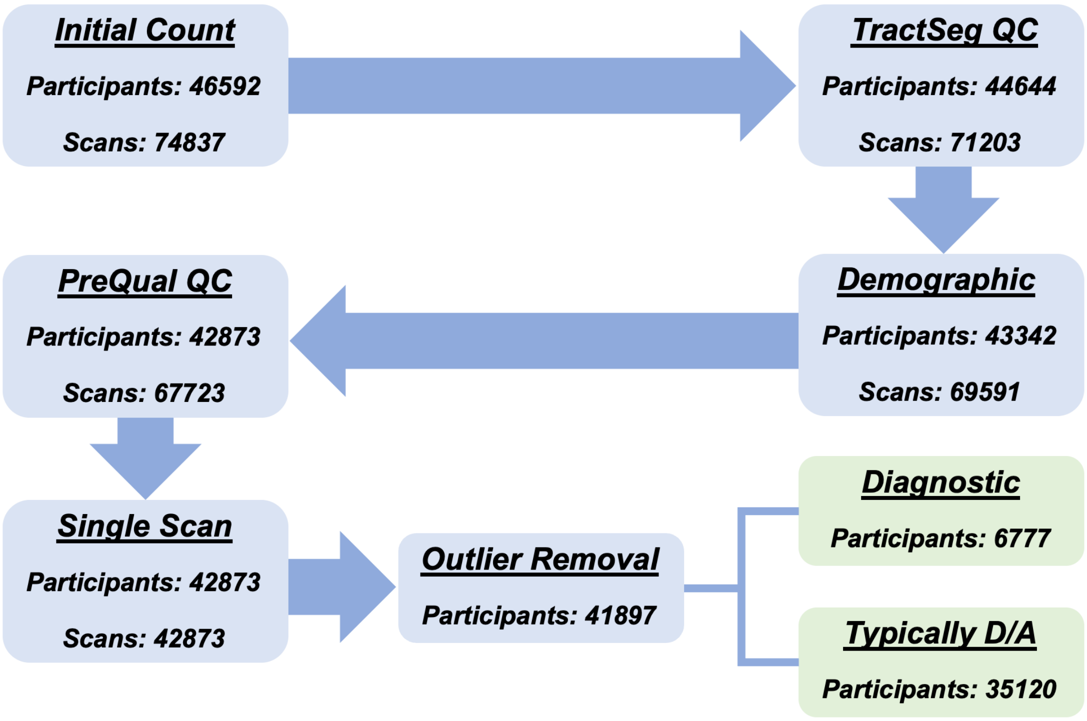


**Supplemental Figure SB.F1. Flowchart of Quality Control Sample Sizes.** Starting with data from 46,592 participants, the several rounds of quality control procedures resulted in a total of 35,120 typically developing and aging participants that were used to create the WM brain charts and 6,777 participants coming from various diagnostic groups.

**Supplemental Table SB.T2.** Demographic information for the typically developing and aging participants used to create the WM brain charts.

| ***Race Category*** | **Hispanic/Latino** | | **Non-Hispanic/Latino** | | **Ethnicity Unreported** | | ***Total*** |
|---|---|---|---|---|---|---|---|
| | *Female* | *Male* | *Female* | *Male* | *Female* | *Male* | |
| **American Indian/Alaska** | 2 | 0 | 7 | 7 | 7 | 4 | *27* |
| **Asian** | 1 | 1 | 63 | 55 | 133 | 141 | *394* |
| **Black or African American** | 2 | 4 | 430 | 205 | 567 | 288 | *1496* |
| **White** | 53 | 51 | 1537 | 992 | 6413 | 4827 | *12873* |
| **More than One** | 36 | 52 | 44 | 59 | 70 | 52 | *313* |
| **Unknown** | 134 | 186 | 3667 | 3977 | 5774 | 5278 | *19016* |
| ***Total*** | *228* | *294* | *5748* | *5295* | *12965* | *10590* | *35120* |

***Total Number of Males:*** *16179* | ***Total Number of Females:*** *18941*

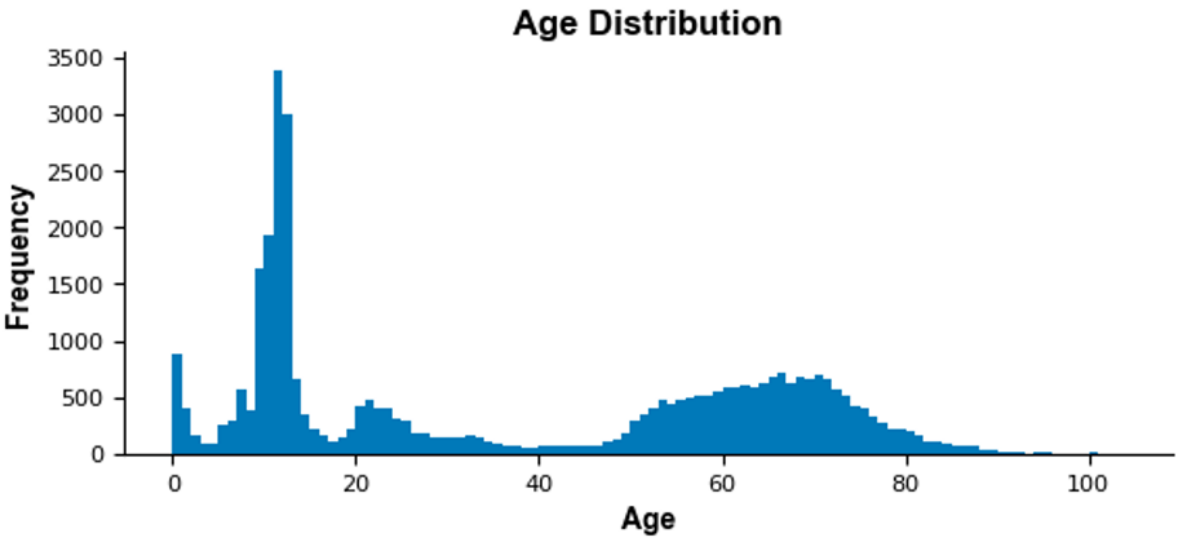


**Supplementary Figure SB.F2. Age histogram for participants used in normative modeling.** Note that ages are binned in 1-year intervals.

**Supplemental Table SB.T3.** Acquisition information for dMRI data from each dataset, along with the respective scanning/facility location.

| Dataset | Voxel Size | Acquisition Parameters | Facility Location |
|---|---|---|---|
| ABCD[1] | 1.7 x 1.7 x 1.7 | 7 b0; 6 b500; 15 b1000; 15 b2000; 60 b3000 | Varied (USA) |
| ADNI[2] | 2.0 x 2.0 x 2.0 | (Varied) 7 b0; 48 b1000 | Varied (USA) |
| Ageility[3] | 2.0 x 2.0 x 2.0 | 1 b0; 64 b1000; 64 b3000 | Newcastle, Australia |
| AOMIC ID1000[4] | 2.0 x 2.0 x 2.0 | 1 b0; 32 b1000 [sequence acquired x3] | Amsterdam, Netherlands |
| AOMIC PIOP1[4] | 2.0 x 2.0 x 2.0 | 1 b0; 32 b1000 | Amsterdam, Netherlands |
| AOMIC PIOP2[4] | 2.0 x 2.0 x 2.0 | 1 b0; 32 b1000 | Amsterdam, Netherlands |
| BANDA[5] | 1.5 x 1.5 x 1.5 | 28 b0; 190 b1500; 190 b3000 | Boston, Massachusetts (USA) |
| BIOCARD[6] | 0.828 x 0.828 x 2.2 | 1 b0; 32 b700 | Baltimore, Maryland (USA) |
| BLSA[7] | 0.8125 x 0.8125 x 2.2[$] | 1 b0; 32 b700 | Baltimore, Maryland (USA) |
| BSNIP1[8] | 1.72 x 1.72 x 3.0 | 1 b0; 32 b1000 | Varied (USA) |
| BSNIP2[9] | 2.0 x 2.0 x 2.0 | 7 b0; 64 b900 | Varied (USA) |
| Calgary[10] | 0.78 x 0.78 x 2.2 | 5 b0; 30 b800 | Calgary, Canada |
| CALM[11] | 2.0 x 2.0 x 2.0 | 5 b0; 64 b1000 | Cambridge, England |
| CAMCAN[12] | 2.0 x 2.0 x 2.0 | 3 b0; 30 b1000; 30 b2000 | Cambridge, England |
| VUMC-ASD | 2.5 x 2.5 x 2.5 | 1 b0; 92 b1600<br>3 b0; 6 b500; 32 b1000; 64 b2000 | Nashville, Tennessee (USA) |
| CUTTING | 2.5 x 2.5 x 2.5 | 1 b0; 60 b2000 | Nashville, Tennessee (USA) |
| dHCP[13] | 1.5 x 1.5 x 1.5 | 20 b0; 64 b400; 88 b1000; 128 b2600 | United Kingdom |
| DLBS[14] | 1.75 x 1.75 x 3.0 | 1 b0; 30 b1000 | Dallas, Texas (USA) |
| EBDT[15] | 2.0 x 2.0 x 2.0 | 7 b0; 42 b1000 | Chapel Hill, North Carolina (USA) |
| HABSHD[16] | 1.72 x 1.72 x 2.5 | 4 b0; 64 b1000 [sequence acquired x3] | Fort Worth, Texas (USA) |
| HBCD[17] | 1.7 x 1.7 x 1.7 | 22 b0; 12 b500; 24 b1000; 36 b2000; 58 b3000 | Varied (USA) |
| HBN[18] | 1.8 x 1.8 x1.8 | 1 b0; 64 b1000; 64 b2000 | Varied (USA) |
| HCP[19] | 1.25 x 1.25 x 1.25 | 18 b0; 90 b1000; 90 b2000; 90 b3000 | St. Louis, Missouri; Minneapolis, Michigan (USA) |
| HCPA[20] | 1.5 x 1.5 x 1.5 | 28 b0; 186 b1500; 184 b3000 | St. Louis, Missouri; Minneapolis, Michigan (USA) |
| HCPBaby[21] | 1.5 x 1.5 x 1.5 | 7 b0; 18 b500; 24 b1000; 108 b1500; 48 b2000; 68 b2500; 170 b3000 | Chapel Hill, North Carolina; Minneapolis, Michigan (USA) |
| HCPD[22] | 1.5 x 1.5 x 1.5 | 28 b0; 186 b1500; 184 b3000 | St. Louis, Missouri; Minneapolis, Michigan (USA) |
| ABC-Babies | 1.78 x 1.78 x2.2 | 1 b0; 32 b700; 64 b2000 | Nashville, Tennessee (USA) |
| IBIS[23] | 2.0 x 2.0 x 2.0 | 1 b0; multi-shell (b=100-1000 in steps of 100), 2-3 volumes/shell | Varied (USA) |
| ICBM[24] | 1.25 x 1.25 x 2.5 | 5 b0; 30 b1000 | Los Angeles, California (USA) |
| Lexical[25] | 2.0 x 2.0 x 2.0 | 1 b0; 64 b1000 | Chicago, Illinois (USA) |
| MASiVar[26] | 2.14 x 2.14 x 2.2 | 34 b0; 86 b1000; 112 b2000 | Boston, Massachusetts (USA) |
| MORGAN | 2.5 x 2.5 x 2.5 | 1 b0; 92 b1600 | Nashville, Tennessee (USA) |
| NACC[27] | 2.0 x 2.0 x 2.0 | (Varied) 1 b0; 64 b1000 | Varied (USA) |
| NKI[28] | 2.0 x 2.0 x 2.0 | 9 b0; 128 b1500 | Rockland County, New York (USA) |
| PING[29] | 2.5 x 2.5 x 2.5 | 1 b0; 32 b1000 | Varied (USA) |
| QTAB[30] | 2.0 x 2.0 x 2.0 | 3 b0; 5 b1000; 15 b3000 | Queensland, Australia |
| ROS[31] | 2.0 x 2.0 x 2.0 | 6 b0; 40 b1000 | Chicago, Illinois (USA) |
| MAP[32] | 2.0 x 2.0 x 2.0 | 6 b0; 40 b1000 | Chicago, Illinois (USA) |
| MARS[33] | 2.0 x 2.0 x 2.0 | 1 b0; 40 b1000 | Chicago, Illinois (USA) |
| SCAN[34] | 2.0 x 2.0 x 2.0 | 13 b0; 6 b500; 48 b1000; 60 b2000 | Varied (USA) |
| SWU[35] | 2.0 x 2.0 x 2.0 | 3 b0; 30 b1000 | Chongqing, China |
| TempleSocial[36,37] | 2.0 x 2.0 x 2.0 | 3 b0; 6 b300; 21 b1000; 24 b2000; 12 b3200; 19 b3300; 61 b5000 | Philadelphia, Pennsylvania (USA) |
| UCLA_LA5c[38] | 1.98 x 1.98 x2.0 | 1 b0; 64 b1000 | Los Angeles, California (USA) |
| UKBB[39] | 2.02 x 2.02 x 2 | 8 b0; 50 b1000; 50 b2000 | United Kingdom |
| UPennRisk[40,41] | 1.875 x 1.875 x 2.0 | 1 b0; 30 b1000 | Philadelphia, Pennsylvania (USA) |
| UTAustin579[42] | 2.0 x 2.0 x 4.0 | 6 b0; 64 b800 | Austin, Texas (USA) |
| VMAP_2.0[43] | 2.33 x 2.33 x 2.5 | 1 b0; 38 b1000; 56 b2000 | Nashville, Tennessee (USA) |
| VMAP[43] | 2.0 x 2.0 x 2.0 | 1 b0; 32 b1000 | Nashville, Tennessee (USA) |
| TN Alzheimer's Project | 2.33 x 2.33 x 2.5 | 1 b0; 38 b1000; 56 b2000 | Nashville, Tennessee (USA) |
| WRAP[44] | 0.9375 x 0.9375 x 2.5 | 8 b0; 40 b1300 | Madison, Wisconsin ((USA) |

[$] BLSA was acquired at 2.2 x 2.2 x 2.2 $mm^3$, but resampled to 0.8125 x 0.8125 x 2.2 $mm^3$ in k-space

**Supplementary Table SB.T4.** The 72 TractSeg[45] tract definitions with the corresponding tract groups. Note that for fitting out-of-sample datasets, the values in the "Tract Name" column should be used for the CSV when performing out-of-sample alignment.

| Tract Name | Tract Description | Tract Group |
|---|---|---|
| AF_left | Arcuate Fasciculus Left | Association |
| AF_right | Arcuate Fasciculus Right | Association |
| IFO_left | Inferior Occipito-Frontal Fascicle Left | Association |
| IFO_right | Inferior Occipito-Frontal Fascicle Right | Association |
| ILF_left | Inferior Longitudinal Fascicle Left | Association |
| ILF_right | Inferior Longitudinal Fascicle Right | Association |
| MLF_right | Middle Longitudinal Fascicle Right | Association |
| MLF_left | Middle Longitudinal Fascicle Left | Association |
| SLF_I_left | Superior Longitudinal Fascicle I Left | Association |
| SLF_I_right | Superior Longitudinal Fascicle I Right | Association |
| SLF_II_left | Superior Longitudinal Fascicle II Left | Association |
| SLF_II_right | Superior Longitudinal Fascicle II Right | Association |
| SLF_III_left | Superior Longitudinal Fascicle III Left | Association |
| SLF_III_right | Superior Longitudinal Fascicle III Right | Association |
| UF_left | Uncinate Fasciculus Left | Association |
| UF_right | Uncinate Fasciculus Right | Association |
| CG_left | Cingulum Left | Association |
| CG_right | Cingulum Right | Association |
| CC_1 | Corpus Callosum 1 (Rostrum) | Commissural |
| CC_2 | Corpus Callosum 2 (Genu) | Commissural |
| CC_3 | Corpus Callosum 3 (Rostral body - premotor) | Commissural |
| CC_4 | Corpus Callosum 4 (Anterior midbody - primary motor) | Commissural |
| CC_5 | Corpus Callosum 5 (Posterior midbody - primary somatosensory) | Commissural |
| CC_6 | Corpus Callosum 6 (Isthmus) | Commissural |
| CC_7 | Corpus Callosum 7 (Splenium) | Commissural |
| CC | Corpus Callosum | Commissural |
| CA | Anterior Commissure | Commissural |
| ATR_left | Anterior Thalamic Radiation Left | Thalamic |
| ATR_right | Anterior Thalamic Radiation Right | Thalamic |
| T_PREF_left | Thalamic-Prefrontal Left | Thalamic |
| T_PREF_right | Thalamic-Prefrontal Right | Thalamic |
| T_PREM_left | Thalamic-Premotor Left | Thalamic |
| T_PREM_right | Thalamic-Premotor Right | Thalamic |
| T_PREC_left | Thalamic-Precentral Left | Thalamic |
| T_PREC_right | Thalamic-Precentral Right | Thalamic |
| STR_left | Superior Thalamic Radiation Left | Thalamic |
| STR_right | Superior Thalamic Radiation Right | Thalamic |
| T_POSTC_left | Thalamic-Postcentral Left | Thalamic |
| T_POSTC_right | Thalamic-Postcentral Right | Thalamic |
| T_PAR_left | Thalamic-Parietal Left | Thalamic |
| T_PAR_right | Thalamic-Parietal Right | Thalamic |
| T_OCC_left | Thalamic-Occipital Left | Thalamic |
| T_OCC_right | Thalamic-Occipital Right | Thalamic |
| ST_FO_left | Striato-Fronto-Orbital Left | Striatothalamic |
| ST_FO_right | Striato-Fronto-Orbital Right | Striatothalamic |
| ST_PREF_left | Striato-Prefrontal Left | Striatothalamic |
| ST_PREF_right | Striato-Prefrontal Right | Striatothalamic |
| ST_PREM_left | Striato-Premotor Left | Striatothalamic |
| ST_PREM_right | Striato-Premotor Right | Striatothalamic |
| ST_PREC_left | Striato-Precentral Left | Striatothalamic |
| ST_PREC_right | Striato-Precentral Right | Striatothalamic |
| ST_POSTC_left | Striato-Postcentral Left | Striatothalamic |
| ST_POSTC_right | Striato-Postcentral Right | Striatothalamic |
| ST_PAR_left | Striato-Parietal Left | Striatothalamic |
| ST_PAR_right | Striato-Parietal Right | Striatothalamic |
| ST_OCC_left | Striato-Occipital Left | Striatothalamic |
| ST_OCC_right | Striato-Occipital Right | Striatothalamic |
| CST_left | Corticospinal Tract Left | Projection |
| CST_right | Corticospinal Tract Right | Projection |
| FPT_left | Frontopontine Tract Left | Projection |

| | | |
|---|---|---|
| FPT_right | Frontopontine Tract Right | Projection |
| ICP_left | Inferior Cerebellar Peduncle Left | Projection |
| ICP_right | Inferior Cerebellar Peduncle Right | Projection |
| OR_left | Optic Radiation Left | Projection |
| OR_right | Optic Radiation Right | Projection |
| POPT_left | Parieto-Occipito-Temporal Left | Projection |
| POPT_right | Parieto-Occipito-Temporal Right | Projection |
| SCP_left | Superior Cerebellar Peduncle Left | Projection |
| SCP_right | Superior Cerebellar Peduncle Right | Projection |
| MCP | Middle Cerebellar Peduncle | Projection |
| FX_left | Fornix Left | Projection |
| FX_right | Fornix Right | Projection |

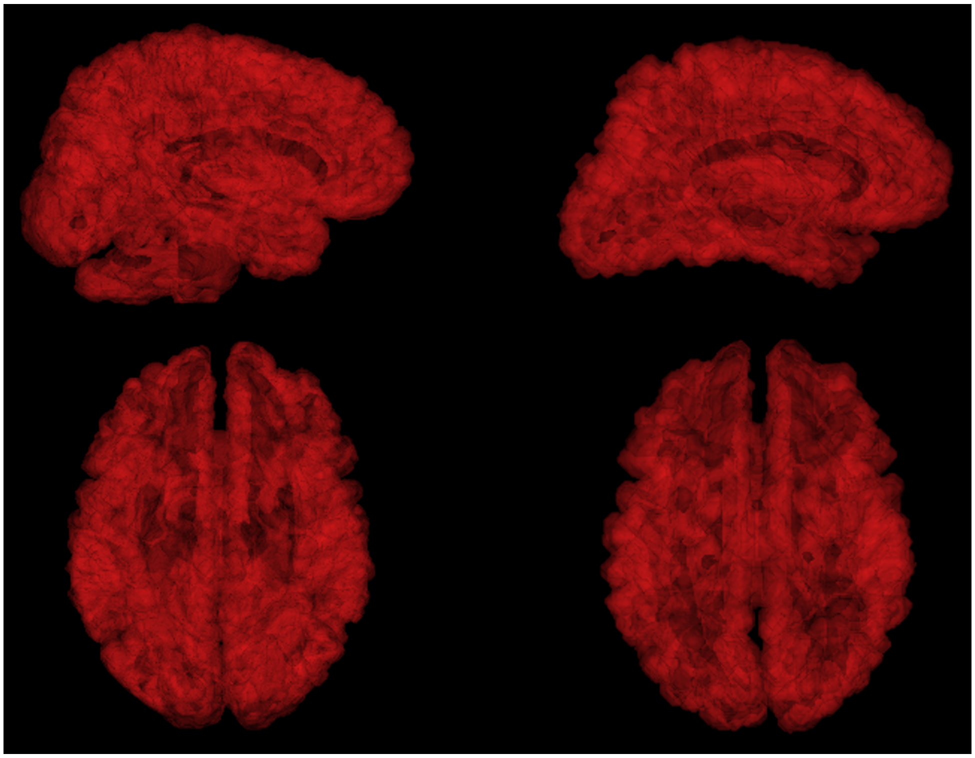

**Supplementary Figure SB.F3. Comparison of TractSeg vs FreeSurfer white matter coverage.** The overlap in coverage between (left) the aggregate of all TractSeg tracts and (right) the white matter mask from FreeSurfer is substantial. The most notable difference is that the FreeSurfer mask does not include the cerebellar or brainstem regions.

### C. **Additional Diagnostic Group Centile Deviations**

#### i. Sample Size of Diagnostic Groups

**Supplementary Table SB.T4. Participants classified as cognitively abnormal for each dataset and diagnosis. Values in** parentheses denote the number dropped from the +⁄− 4 standard deviation outlier removal detailed in the Methods.

| Dataset | Autism | ADHD | Alzheimer's | Anxiety | Bipolar | CN* | Defiance/ Conduct Disorder | Depression | Dyslexia | Epilepsy | MCI | Schizoaffective Disorder | Schizophrenia | Social Anxiety | Substance Abuse | Suicidality and Self-Injury | Other | *Total* |
|---|---|---|---|---|---|---|---|---|---|---|---|---|---|---|---|---|---|---|
| ABCD | | 619 | | 38 | 162 | | 89 | 148 | | | | | 33 | 40 | 25 | 60 | 97 | ***1311*** |
| ADNI | | | 206 (5) | | | | | | | | 439 (7) | | | | | | | ***645 (12)*** |
| BANDA | | | | 79 | | | | 65 | | | | | | | | | | ***144*** |
| BIOCARD | | | 3 | | | 1 | | | | | 29 | | | | | | | ***33*** |
| BLSA | | | 17 (4) | | | 4 | | | | | 32 (1) | | | | | | | ***53 (5)*** |
| BSNIP1 | | | | 1 | 61 (1) | | | 53 (2) | | | | 71 (3) | 125 (3) | | 26 (2) | | 20 (1) | ***357 (12)*** |
| BSNIP2 | | | | | 101 (4) | | | 10 | | | | 166 (4) | 192 (1) | | 15 | | 7 | ***491 (9)*** |
| CALM | 20 | 66 | | | | | | | 22 | | | | | | | | 16 | ***124*** |
| VUMC-ASD | 69 | | | | | | | | | | | | | | | | | ***69*** |
| CUTTING | | | | | | | | | | | | | | | | | 35 | ***35*** |
| HABSHD | | | 36 (1) | | | 34 | | | | | 603 (16) | | | | | | | ***673 (17)*** |
| HBN | 73 | 752 (4) | | 246 | 3 | | 37 (1) | 144 (1) | 69 | | | | 1 | 52 | | | 93 (2) | ***1470 (8)*** |
| IBIS | 50 (2) | | | | | | | | | | | | | | | | | ***50 (2)*** |
| MAP | | | 14 (1) | | | 1 | | | | | 105 (2) | | | | | | | ***120 (3)*** |
| MARS | | | | | | | | | | | 29 (1) | | | | | | | ***29 (1)*** |
| MORGAN | | | | | | | | | | 141 | | | | | | | 3 | ***144*** |
| NACC | | | 160 (3) | | | | | | | | 32 (3) | | | | | | | ***192 (6)*** |
| NKI | | 47 | | 27 | | | | 50 | | | | | | | 143 | | 98 | ***365*** |
| ROS | | | | | | | | | | | 11 | | | | | | | ***11*** |
| SCAN | | | 21 (1) | | | | | | | | 51 (2) | | | | | | | ***72 (3)*** |
| UCLA | | 40 | | | 46 | | | | | | | | 46 (1) | | | | | ***132 (1)*** |
| VMAP | | | 33 | | | 1 | | | | | 116 (2) | | | | | | | ***150 (2)*** |
| VMAP_TAP | | | 43 (2) | | | | | | | | 58 (3) | | | | | | | ***101 (5)*** |
| WRAP | | | 2 | | | | | | | | 4 | | | | | | | ***6*** |
| ***Total*** | ***212 (2)*** | ***1524 (4)*** | ***535 (17)*** | ***391*** | ***373 (5)*** | ***41*** | ***126 (1)*** | ***470 (3)*** | ***91*** | ***141*** | ***1509 (37)*** | ***237 (7)*** | ***397 (5)*** | ***92*** | ***209 (2)*** | ***60*** | ***369 (3)*** | ***6777 (87)*** |

#### ii. Centile Score Deviations of Other Diagnostic Groups

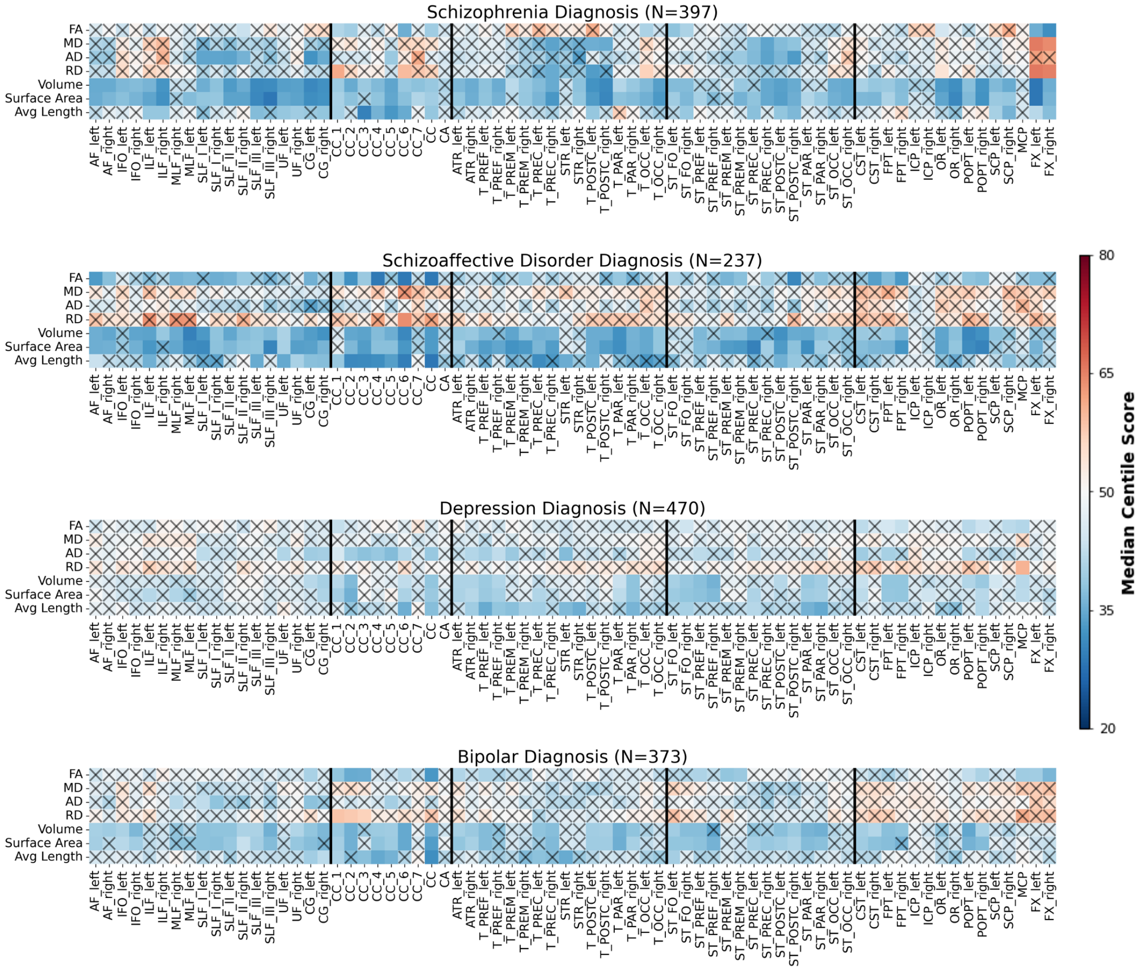


**Supplementary Figure SC.F1. Centile score differences across major depressive and psychotic disorders.** All major psychoses and major depressive disorder (top to bottom: schizophrenia, schizoaffective disorder, major depressive disorder, and bipolar disorder) showed localized reductions in tract macrostructure and microstructure under the one sample Wilcoxon after Bonferroni correction for multiple comparisons.

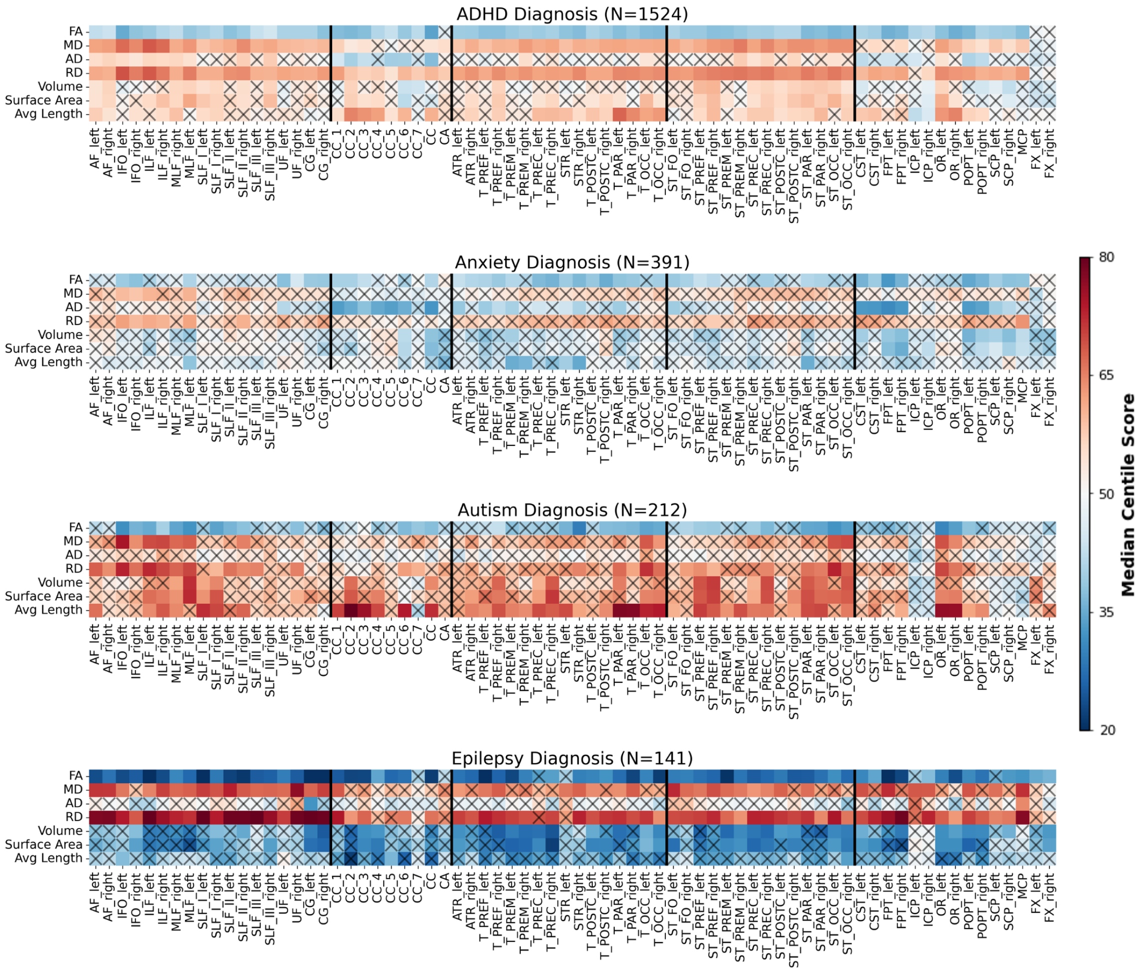


**Supplementary Figure SC.F2. Centile score differences across other neurocognitive disorders with significant deviations.** For many other clinical cohorts (top to bottom: attention-deficit hyperactivity disorder (ADHD), anxiety, autism spectrum disorder, and epilepsy), we observe either localized deviations in tract microstructure and macrostructure or consistent deviations across all tracts within specific macrostructural and microstructural features. Individuals with epilepsy and ADHD showed substantial deviations in FA and RD across the WM, suggesting diffuse microstructural abnormalities, alongside more localized reductions in volume, length, and surface area in select tracts. For individuals diagnosed with autism spectrum disorder, we observe increases in macrostructure across tracts, with a subset of tracts showing significant deviations. Finally, we observe sparsely significant deviations in microstructure and macrostructure for individuals diagnosed with anxiety.

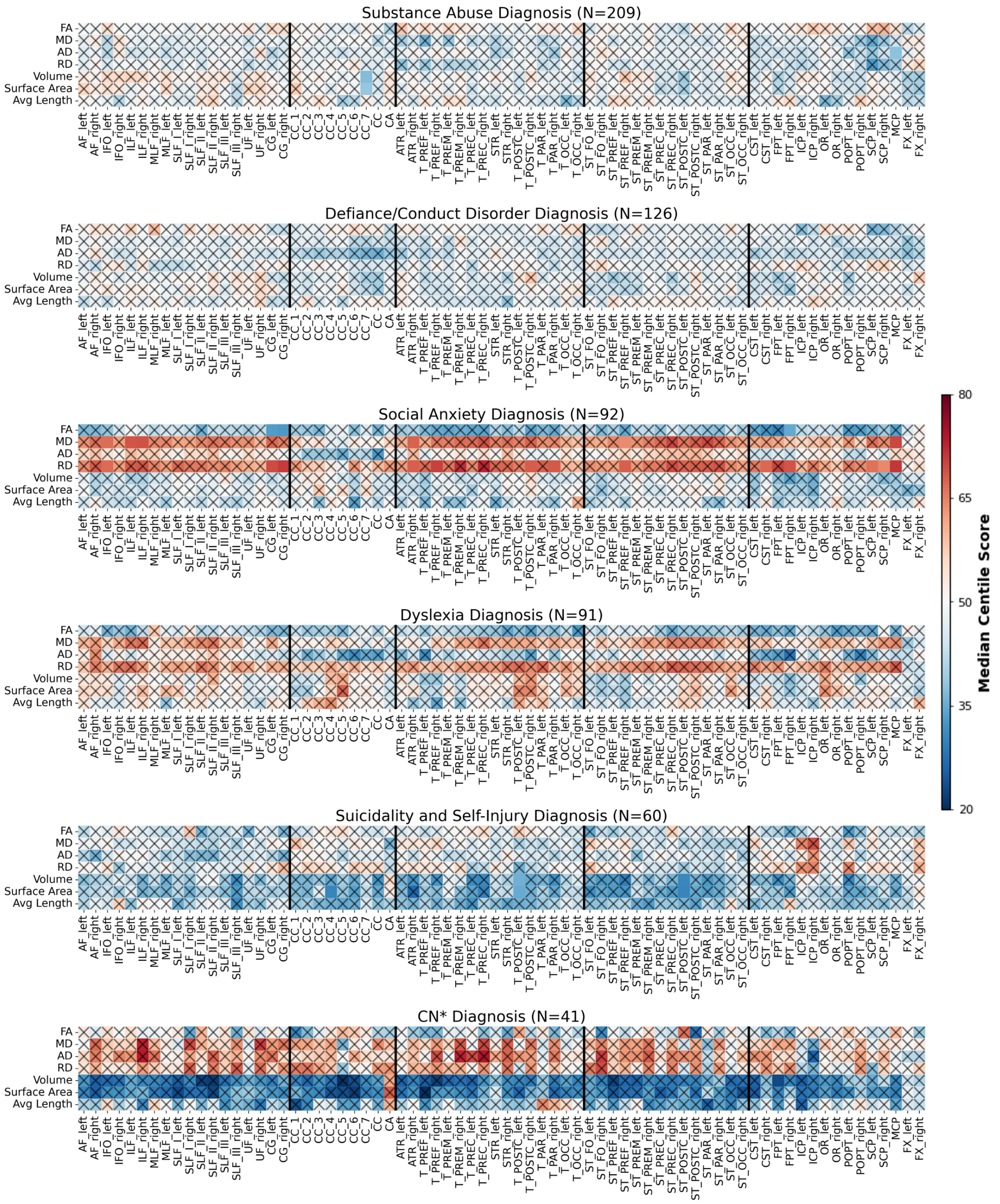


**Supplementary Figure SC.F3. Centile score differences across other neurocognitive disorders without significant deviations.** For many other clinical cohorts (from top to bottom: substance abuse, defiance/conduct disorders, social anxiety disorder, dyslexia, suicidality and self-injury, and individuals who start as cognitively unimpaired and then transition to cognitive impairment, or CN*), we do not observe significant deviations in centile scores. However, there are still non-significant patterns across tracts, such as reduced tract macrostructure in CN* participants and increased MD/RD in the social anxiety cohort.

##### iii. Effect Sizes of Centile Score Deviations

For all comparisons of clinical/non-typical cohorts, we use the two-sided Wilcoxon test, a non-parametric test, to assess significantly different centile distributions. Thus, we calculate effect size as the magnitude of the rank-biserial coefficient $r_b$:

$$r_b = 1 - \frac{4|W|}{N(N+1)} \quad (1)$$

where $W$ is the two-sided Wilcoxon statistic and $N$ is the sample size. For the cohorts with consistently significant features, we observe consistently larger effect sizes across tracts (**Supplementary Figure SC.F4, SC.F5**). Effect sizes for the other cohorts, which consist of mostly non-significant features, have much smaller and more sporadic effect sizes (**Supplementary Figure SC.F6**). Still, we observe moderate effect sizes in some features for some of these diagnostic groups. We also observe large effect sizes for the nCMD measurements (**Supplementary Table SC.T2**); however, we note that for cohorts with smaller sample sizes, the calculated effect sizes may be over or underinflated.

**Supplemental Table SC.T2.** Rank-biserial effect sizes for non-typical cohorts of the normalized centile mahalanobis distance (nCMD) metrics.

| Cohort | nCMD Micro | nCMD Macro | nCMD Combined |
|---|---|---|---|
| Autism (N=212) | 0.436 | 0.495 | 0.558 |
| MCI (N=1509) | 0.549 | 0.212 | 0.485 |
| Alzheimer's Disease (N=535) | 0.741 | 0.432 | 0.712 |
| ADHD (N=1524) | 0.057 | 0.303 | 0.162 |
| Schizophrenia (N=397) | 0.457 | 0.040 | 0.407 |
| Depression (N=470) | 0.031 | 0.059 | 0.103 |
| Anxiety (N=391) | 0.140 | 0.152 | 0.052 |

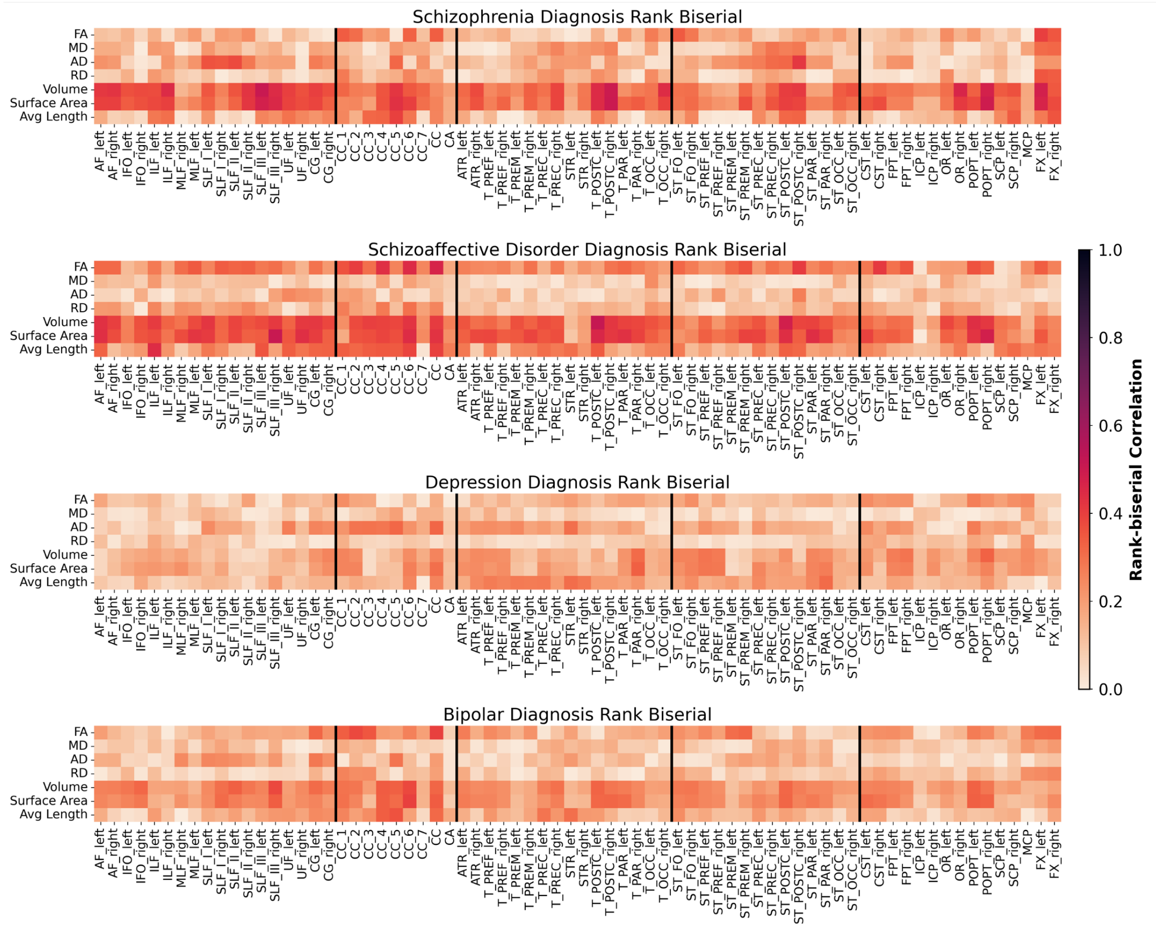


**Supplementary Figure SC.F4. Effect size magnitudes of centile distances for major depressive and psychotic diagnostic groups.** Rank-biserial correlations (magnitude) of major depressive and psychotic diagnostic group centile score differences from the median centile (from top to bottom, schizophrenia, schizoaffective disorder, depression, bipolar disorder). Larger effect sizes were observed for macrostructural measures when assessing schizophrenia, schizoaffective disorder, and bipolar disorder group deviations.

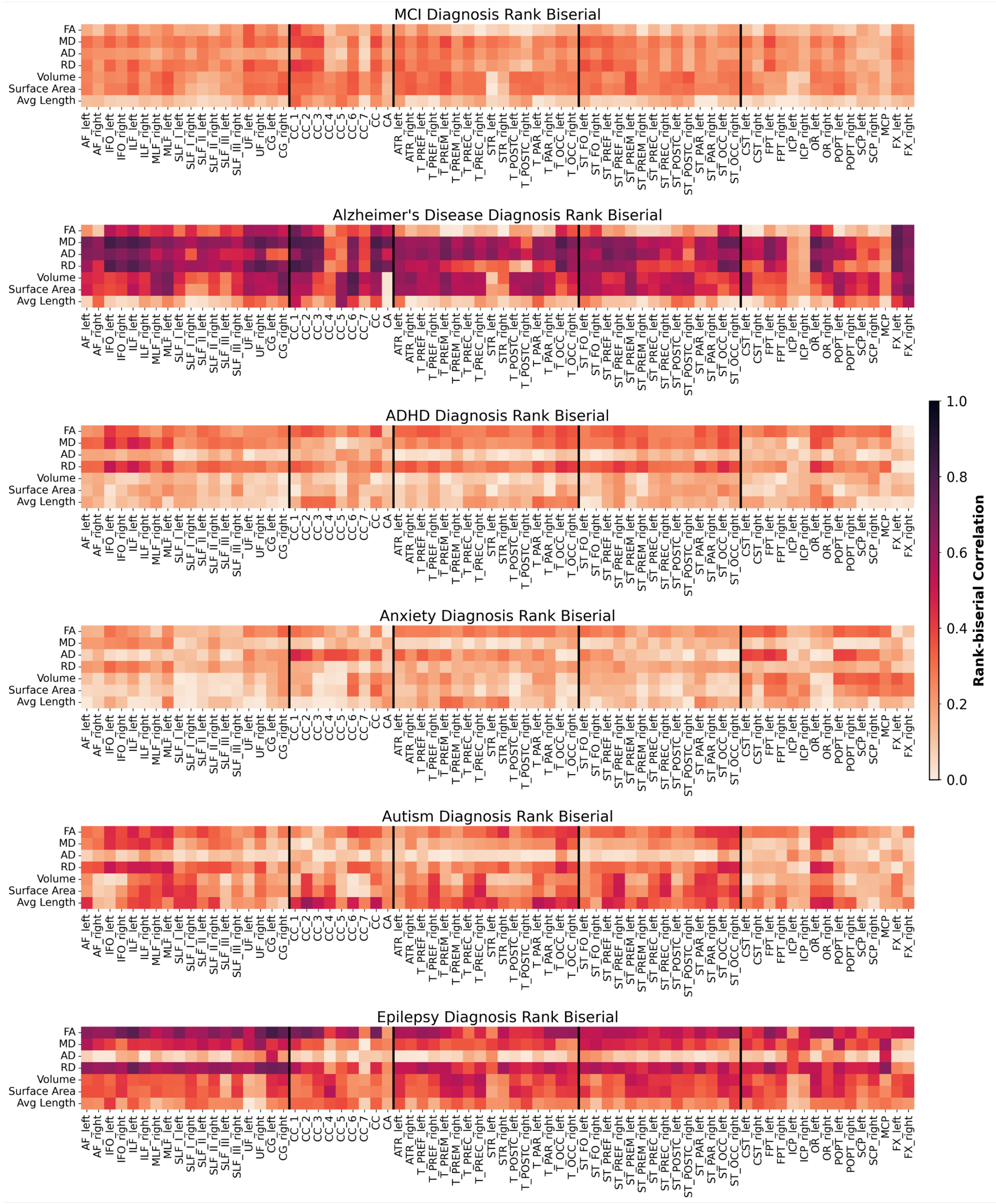


**Supplementary Figure SC.F5. Effect size magnitudes of centile distances for other diagnostic groups with significant effects.** Rank-biserial correlations (magnitude) of centile score differences from the median centile for other diagnostic groups with significant deviations (from top to bottom, mild cognitive impairment, Alzheimer's disease, ADHD, anxiety, autism spectrum disorder, epilepsy). Large effect sizes were observed across most features for Alzheimer's disease as well as FA, MD, RD for epilepsy. Moderate effect sizes were seen across features for the mild cognitive impairment diagnostic group and for several features of the ADHD, autism, and anxiety diagnostic groups.

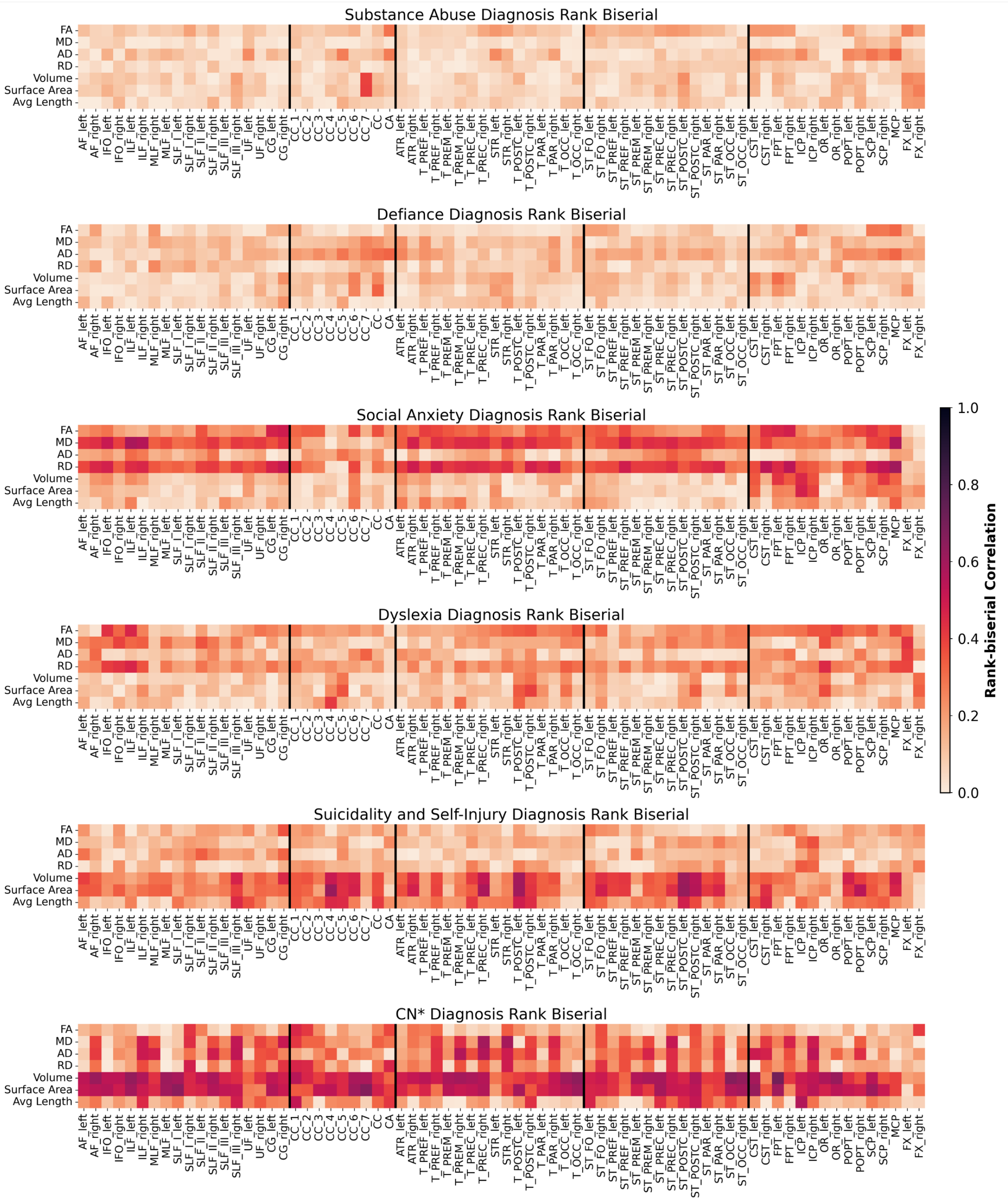


**Supplementary Figure SC.F6. Effect size magnitudes of centile distances for other diagnostic groups with non-significant effects.** Rank-biserial correlations (magnitude) of centile score differences from the median centile for other diagnostic groups with non-significant deviations (from top to bottom, substance abuse, defiance/conduct disorder, social anxiety, dyslexia, suicidality and self-injury, and individuals who start as cognitively unimpaired and then transition to cognitive impairment, or CN*). Although we did not observe significant centile score deviations for these diagnostic groups, the differences have moderate effect sizes for macrostructural features in the CN* and suicidality and self-injury diagnostic groups, as well as FA, MD, RD for social anxiety.

### D. **Model Fitting**

#### i. Number of fractional polynomial terms for $\boldsymbol{\mu}$ and $\boldsymbol{\sigma}$

Upon normalization, the macrostructural features appear to have a slightly less complex dependency on age, as evidenced by the decrease in number of terms for best fit models of $\mu$ (**Supplemental Figures SD.F1, SD.F2**). This trend is more apparent for surface area than it is for volume or average length. We hypothesize this result may be due to a removal of variance introduced by the normalization of global cerebral tissue volume estimates, as people with larger heads will likely have larger WM tract volumes as well. However, this hypothesis was not formally tested in this work. For $\sigma$, however, there appears to be no large change in how non-linear the relationship with age is after normalization. Additionally, $\sigma$, which is related to the variance or spread of the distribution, appears to have a less complex age-related effect than the location of the distribution, or $\mu$. This can be seen in the majority of best fit models having 2 or 3 fractional polynomial terms for $\mu$, whereas most models have only 1 or 2 terms for $\sigma$.

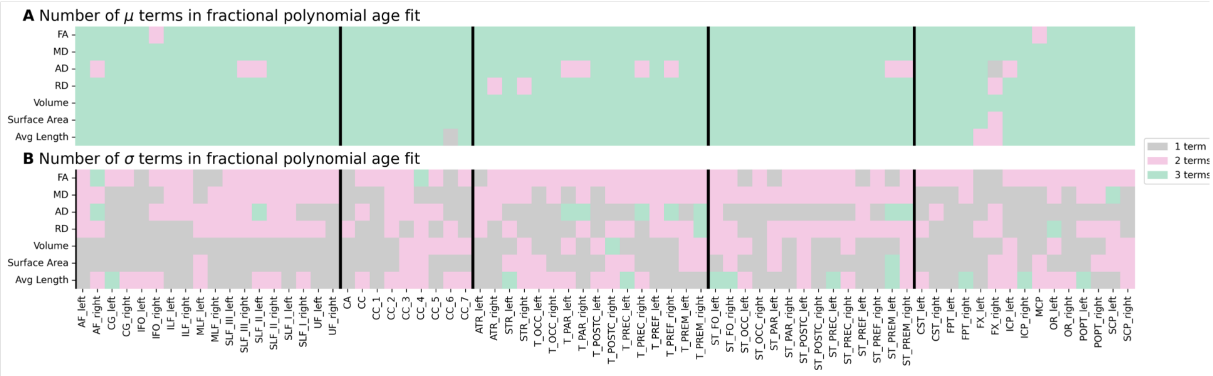


**Supplementary Figure SD.F1. Fractional polynomial term numbers.** The greater number of fractional polynomial terms for A.) $\mu$, related to the location of the generalized gamma distribution, as opposed to B.) $\sigma$, which is related to the scaling of the distribution, suggests that the location of the distribution has a more non-linear relationship with age than the scaling.

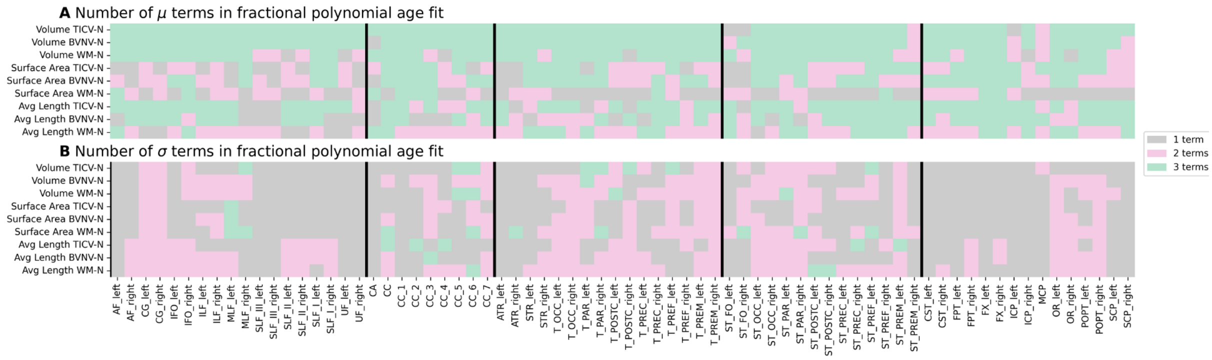


**Supplementary Figure SD.F2. Fractional polynomial term numbers – normalized measures.** Normalized macrostructural measures appear to have a less non-linear relationship with age than unnormalized measures, demonstrated by the shift from green boxes in **Supplemental Figure SD.F1** (above) to pink and gray boxes for A.) the $\mu$ term. However, normalization of macrostructural measures does not appear to change the non-linearity of B.) the $\sigma$ term when compared to the unnormalized measures.

#### ii. Empirical Model Stability Analysis

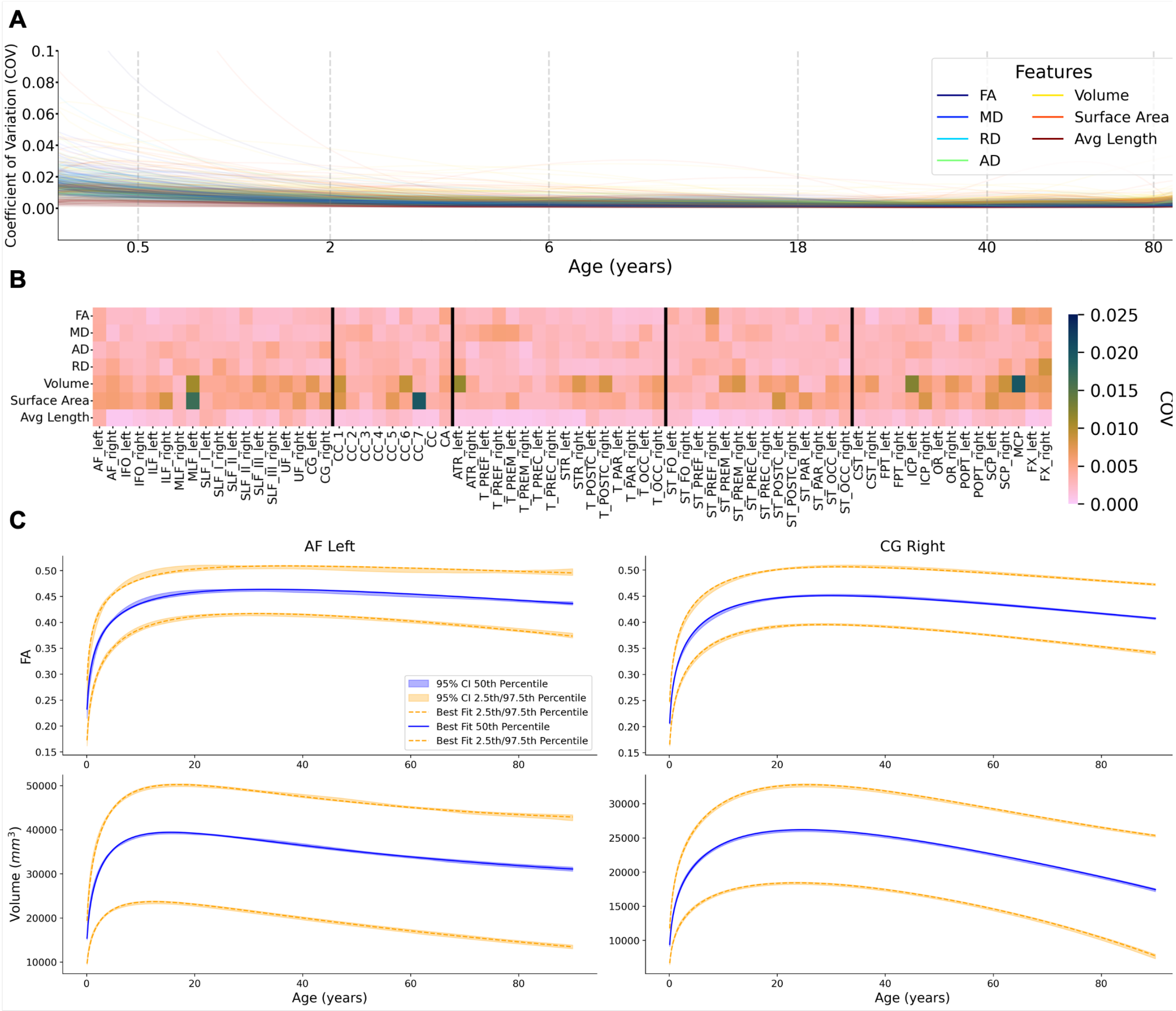


**Supplementary Figure SD.F3. Model stability assessment using leave-one-study-out (LOSO) bootstrapping.** We observe WM brain chart models to be stable across the entire lifespan. (A) Coefficient of variation (COV) from a LOSO bootstrapping analysis on the WM brain charts suggest a high stability across the majority of the lifespan for all models, with most models having a maximum of 1% COV at a given point in the lifespan. Age axis is log-scaled to highlight the spread of models at the beginning of the lifespan. (B) Averaging COV across all points in the lifespan demonstrate that most models have high stability as seen by low COV, with macrostructural models having a slightly larger COV on average. (C) Example plots of median (blue), and 2.5$^{th}$ and 97.5$^{th}$ (orange) percentiles with 95% confidence intervals derived from the LOSO bootstrapping show high stability across the entire lifespan and percentiles (top row is fractional anisotropy, bottom row is volume; left column is the left arcuate fasciculus (AF left), right column is the right cingulate gyrus (CG right)).

#### iii. Voxel Size Effect on FA

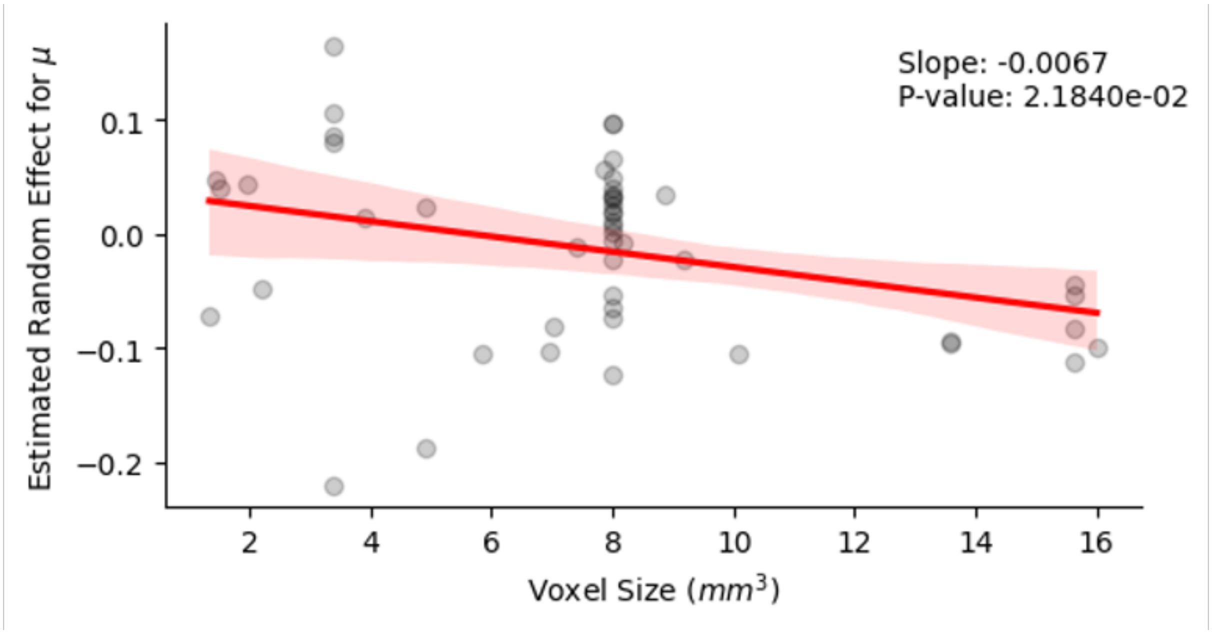


**Supplementary Figure SD.F4. Voxel size correlation with dataset-estimated shift.** We observe that for the whole-brain white matter FA, voxel size has a significant negative correlation with the estimated values for the GAMLSS random effect terms for μ. This indicates that the GAMLSS model is appropriately modeling acquisition-related variability across datasets for the brain charts. Note that in the GAMLSS formulation, μ is log-transformed, and thus the random effect terms are multiplicative instead of additive. For example, a random effect value of -0.1 corresponds to a multiplicative shift in the dataset values by $e^{-0.1} = 0.904$.

#### iv. Batch Correction and Site Harmonization

As described in the Methods section, we use "primary study" or "dataset" as the batch effect term to be estimated in GAMLSS. However, for many studies there are subsets of the data acquired with different acquisition parameters or physical scanners. Thus, we run a sensitivity analysis comparing the estimated centile scores of participants when using "dataset" as a batch variable to centile scores estimated using a combination of physical site, scanner, and acquisition as the batch variable (**Supplemental Figure SD.F5**). Across all tracts, the correlations between "dataset"-batch and "site-scanner-acquisition"-batch centile scores for participants were higher for macrostructural and lower for microstructural features. However, all correlations were greater than 0.88, with the majority being above 0.95. Within the microstructural features, AD centile scores showed the lowest correlation coefficients on average, while RD correlations were higher on average. Further, some tracts appeared to have higher correlation coefficients than others on average, with no distinct trends within tract groups.

While a nested covariance structure for site, scanner, and acquisition might be a more appropriate modeling of the batch effects, GAMLSS does not currently support specification of such a nested covariance structure. Consequently, in the model, any given "batch" for a subset of a dataset is inherently considered to be as different from any other subset as it is from any other primary dataset. If such a nested covariance structure could be specified, the centile scores might be even more highly correlated between the “dataset”-batch and “site-scanner-acquisition”-batch models. While we have reduced variability introduced in data processing through containerization of all pipelines, these data processing considerations cannot account for variability introduced in any part of the raw data acquisition.

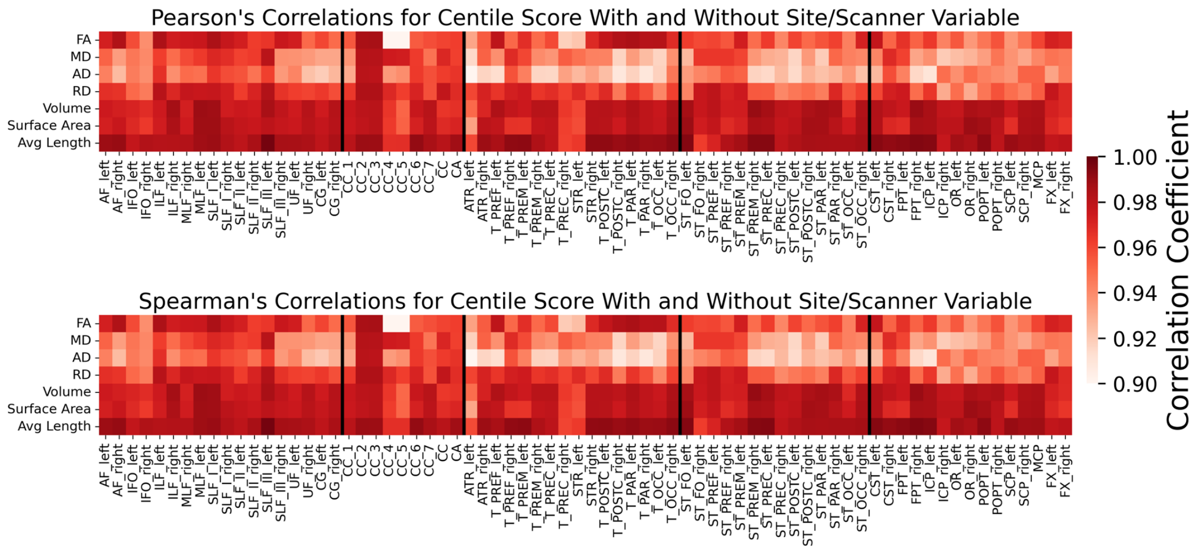


**Supplementary Figure SD.F5. Correlation of centile scores for scanner-site-acquisition-batch vs dataset-batch models.** When encoding site, dataset, and scanner as the random effect to be estimated in the GAMLSS model fitting, correlations of centile scores are high when compared to the model that uses only dataset as the random effect variable. Both (Top) Pearson’s and (Bottom) Spearman’s correlation coefficients of centile scores are above 0.88 for all models, with most models above 0.95. Macrostructural model correlations are noticeably higher than those for microstructural feature models. Additionally, AD models appear to have the lowest average correlation out of all feature models.

#### v. Sex-related Effects on White Matter Brain Charts

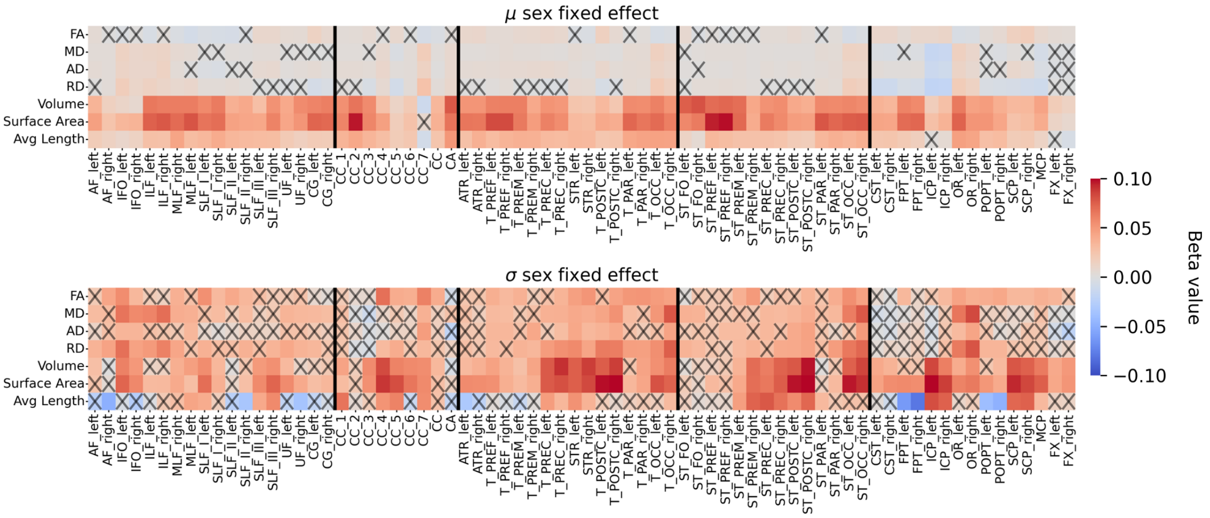


**Supplementary Figure SD.F6. Significance of sex fixed effect across models**. In the GAMLSS fitting, we find that the fixed effect of sex for $\mu$ (top) is significant for most models, whereas the sex fixed effect for $\sigma$ (bottom) is much less significant (after Bonferroni correction for multiple comparisons). Positive (red) indicates that corresponding trajectory values for males are larger than those of females, whereas blue indicates the opposite. Macrostructural trajectories tend to have much larger values for $\mu$ than microstructural trajectories. All trajectories tend to have larger male variability, indicated by the positive (red) values for the $\sigma$ fixed effect. Note that, due to the GAMLSS model specification, fixed effect values are logarithmically scaled and multiplicative in nature.

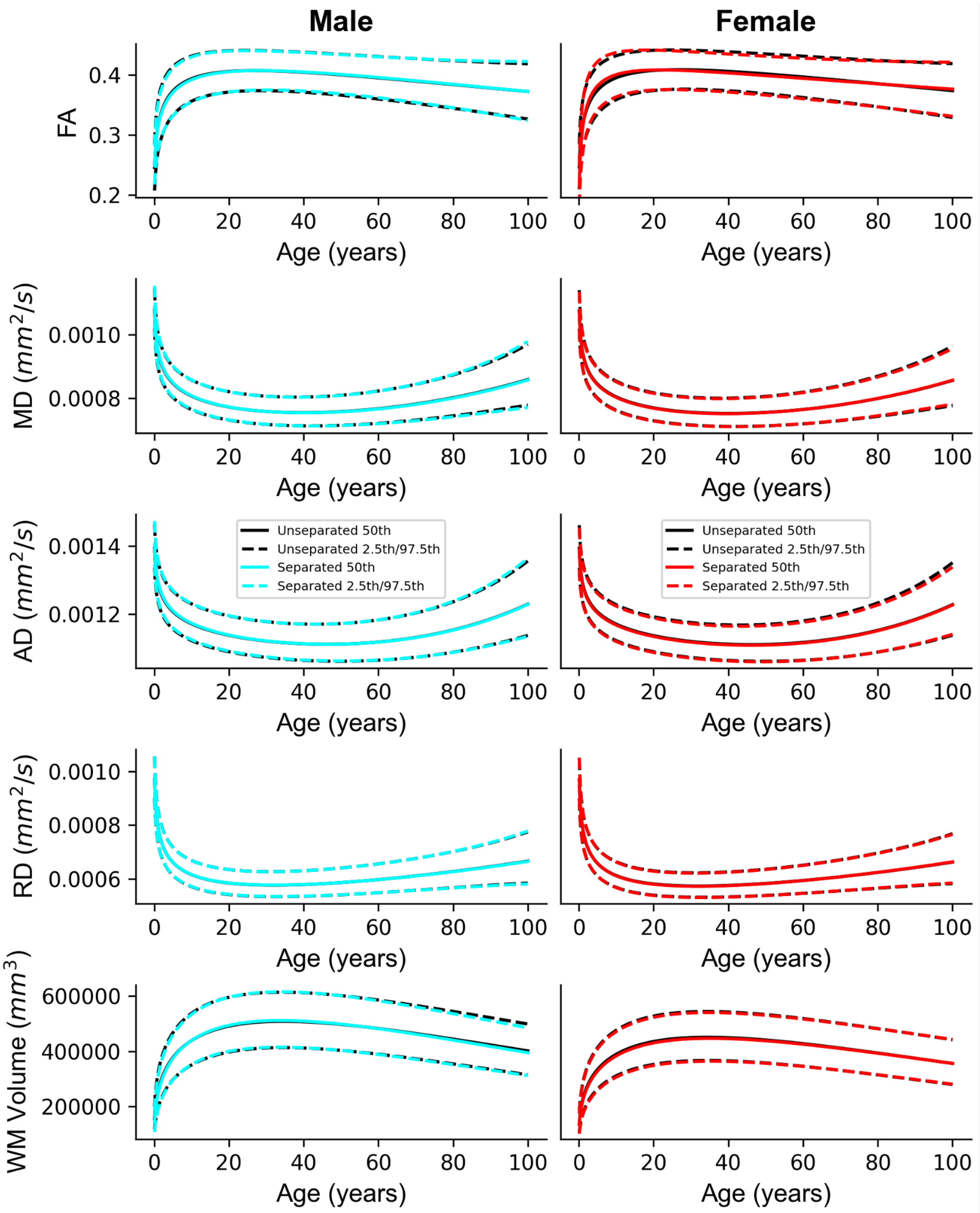


**Supplementary Figure SD.F7. Comparison of sex-separated vs unseparated models.** Normative models fit with sex as a linear effect (”Unseparated” - black lines) are very similar to models that were fit with male and female populations completely separated (“Separated” – cyan/red lines), indicating that the GAMLSS model fitting for the WM brain charts is appropriately capturing the sex-specific trajectories for microstructure and macrostructure (from top to bottom: global WM features of FA, MD, AD, RD, and volume). Male trajectories are plotted in the left and female trajectories are plotted on the right.

##### vi. Maxima and Minima of White Matter Tract Trajectories

Aggregated across all 72 tracts (**Supplemental Figure SD.F8**), we observed systematic variation in the timing of developmental milestones. Among microstructural metrics, axial diffusivity (AD) generally peaked later than other features, suggesting prolonged maturation of myelin-sensitive properties. Within several tract classes, we also observed an anterior-to-posterior patterning - where tracts located anteriorly reached peak maturation earlier than more posterior tracts, reflecting coordinated spatial gradients in WM development[46].

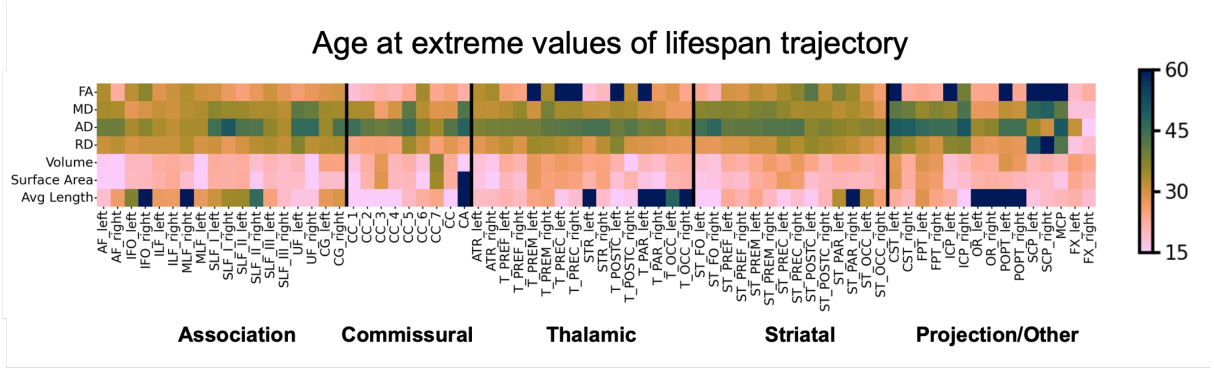


**Supplementary Figure SD.F8. Age at extreme values of tract brain charts.** A heatmap summarizes the estimated age at which each feature reaches its extreme value (peak or trough) across all 72 measured tracts, organized by tract class. This demonstrates systematic variations in milestone timing depending on the specific feature, pathway, and tract group. Across many tracts, macrostructural features tend to reach their milestones earlier in life compared to microstructural features.

#### E. **Comparison to Existing Reference Charts**

The investigation of age-related changes in WM using neuroimaging data has been a well-established area of research for decades. Fewer, more recent works have been published on the use of neuroimaging data to construct growth charts for WM measurements[47,48], but have not been as extensive with regard to the number of features. Specifically, Bethlehem et al. only investigate global WM volume, whereas Zhu et al. investigate only microstructure through FA.

##### i. White Matter Lifespan Modeling in the Literature

**Supplemental Table SE.T1.** Past literature studying white matter lifespan trajectories.

| Study | Age Range (years) | Participants | Datasets | Microstructure | Macrostructure | White Matter Info |
|---|---|---|---|---|---|---|
| Storsve et al. 2016 [49] | 23-87 | 201 (402 sessions) | 1 | DTI | N/A | TRACULA (18 tracts) |
| Giorgio et al. 2010 [50] | 23-81 | 66 | 1 | DTI | N/A | WM Voxels |

| | | | | | | |
|---|---|---|---|---|---|---|
| Beck et al. 2021 [51] | 18-94 | 573 (702 sessions) | 2 | DTI, NODDI, DKI, RSI, WMTI, SMT mc: 6 models, 20 scalars | N/A | TBSS skeleton |
| Henriques et al. 2023 [52] | 18-88 | 636 | 1 | DTI, DKI, NODDI | N/A | JHU Atlas (48 ROIs) |
| Toschi et al., 2020 [53] | 13-62 | 91 | 1 | DTI, CHARMED | N/A | TBSS skeleton + JHU overlap (50 ROIs) |
| Lebel et al. 2012 [54] | 5-83 | 403 | 1 | DTI | WM Volume | 12 pathways (Manual) |
| Slater et al. 2019 [55] | 7-84 | 801 | 1 | DTI, NODDI, g-ratio | N/A | 20 bilateral + callosal tracts (AFQ) |
| Yeatman et al. 2014 [56] | 7-85 | 102 | 1 | DTI (FA,MD only) | N/A | 24 pathways (AFQ) |
| Groves et al. 2012 [57] | 8-85 | 484 | 2 | DTI (FA,MD,MO) | N/A | TBSS skeleton + ICA |
| Hoagey et al. 2019 [58] | 20-94 | 186 | 1 | DTI (FA,MD) | N/A | Population-based WM Voxels |
| Kochunov et al. 2011 [59] | 11-90 | 1031 | 1 | DTI (FA only) | N/A | TBSS skeleton + JHU overlap (11 ROIs) |
| Bender et al. 2015 [60] | 19-78 | 96 (192 sessions) | 1 (2 timepoints ) | DTI (FA,AD,RD) | N/A | TBSS skeleton + JHU overlap (13 ROIs) |
| Sexton et al. 2014 [61] | 20-84 | 203 (406 sessions) | 1 (2 timepoints ) | DTI | N/A | TBSS skeleton + manual ROIs (4 ROIs) |
| Sala et al. 2012 [62] | 13-70 | 84 | 1 | DTI | Tract volume | 8 ROIs from FA atlas |
| Westlye et al. 2010 [63] | 8-85 | 430 | 2 | DTI | WM Volume | TBSS skeleton + JHU overlap (16 ROIs) |
| Billiet et al. 2015 [64] | 17-70 | 59 | 1 | DTI, NODDI, DKI, MET2 | N/A | Voxel-wise in WM mask; also 17 JHU ROIs |
| Molloy et al. 2021 [65] | 18-75 | 79 | 1 | DTI | N/A | TBSS skeleton + JHU overlap (20 ROIs) |
| Ardekani et al. 2007 [66] | 26-69 | 20 | 1 | DTI | N/A | 6 Manually Traced ROIs |
| Schilling et al., 2023 [67] | 0-100 | 2789 | 4 | DTI,NODDI | Tract volumes, lengths, areas, etc. | 63 TractSeg ROIs |

| | | | | | | |
|---|---|---|---|---|---|---|
| Conte et al. 2024 [68] | 2wk-100yrs | 6186 | 12 | FA,MD | WM Volume; Tract Density | XTRACT (29 tracts) |

#### ii. Comparison to Bethlehem et al.

As our method for assessing global WM macrostructure follows Bethlehem et al.[48] in the use of FreeSurfer and GAMLSS, we can more directly compare our brain charts to theirs (**Supplemental Figure SE.F1**). Notably, our curves follow the same trend of a rapid increase during infancy that plateaus during early adulthood, then finally dropping off during aging. We believe this agreement in capturing the general trend indicates that our brain charts are comparable to other established normative models for neuroimaging.

However, we note that there are some slight differences in the two sets of global WM volume curves. While the age at peak is not very representative, this value has tended to vary in the literature quite a bit as well. Specifically, our observed peak for WM volume is slightly less that Lebel et al. (peak at 37 years) [14], about 15 years later than Courchesne et al. (beginning of the 4th decade) [15] and 5 years later than Bethlehem et al. [13], and 10 years earlier than Giorgio et al.[16]. Furthermore, the Bethlehem et al. curves were fit using pre-natal data and used multiple different methods to obtain WM volume measurements, specifically, different FreeSurfer versions and hand drawn ROIs (**Supplemental Table SE.T1**). In contrast, our curves contained only post-natal data and stayed consistent with our data processing approach. Finally, Bethlehem et al. included multiple scans per participant, which was not explicitly modeled in their GAMLSS formula, whereas we only include one scan per participant. These factors, in addition to the largely increased sample size used by Bethlehem et al., could be contributing to the slight differences between the curves.

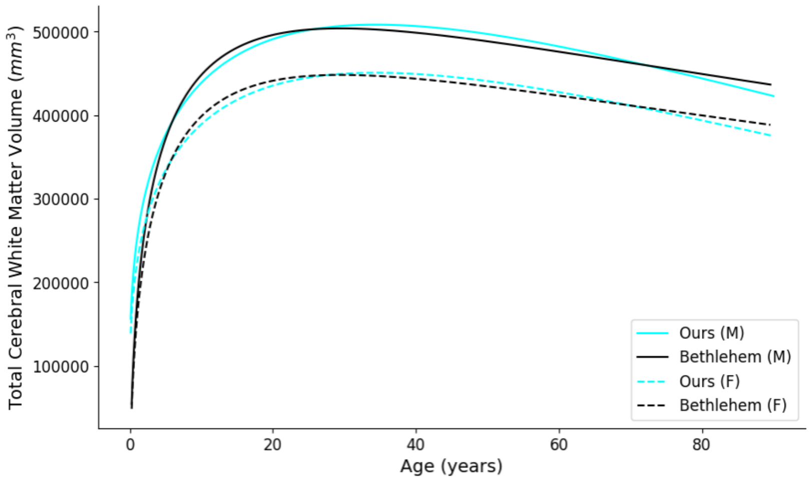


**Supplementary Figure SE.F1. Comparison of our global white matter volume curve to Bethlehem et al.** Our brain charts follow the same trend as Bethlehem et al., which is a rapid increase during infancy that plateaus during early adulthood and finally drops off during aging. We believe this agreement in capturing the general trend indicates that our brain charts are comparable to other established normative models for neuroimaging.

**Supplemental Table SE.T2.** Comparison of methods for our work to Bethlehem et al. (2022)

| *Study* | *Participants* | *Repeated Scans* | *Normative Modeling* | *Processing Variability* | *Distribution Family* | *Out-of-sample Centile Scoring* | *Public release* |
|---|---|---|---|---|---|---|---|
| **Bethlehem** | 101,457 | Yes | GAMLSS; fractional polynomials for age | Yes; modeled in GAMLSS | Generalized Gamma | Yes, MLE | Website* |
| **Our Work** | 35,120 | No | GAMLSS; fractional polynomials for age | No | Generalized Gamma | Yes, MLE | Zenodo& |

*https://brainchart.shinyapps.io/brainchart/; &https://zenodo.org/records/17561821

##### iii. Comparison to Zhu et al.

The FA brain charts created by Zhu et al. use the preprocessing pipeline defined by the ENIGMA consortium[69] to obtain quantitative values of microstructure within regions of interest (ROIs) defined by the JHU DTI atlas (**Supplemental Table SE.T2**). While these atlas-based approaches are commonly used methods for quantifying microstructure within specific brain regions, we note that such practices are not subject specific. Furthermore, these methods do not permit assignment of a single voxel to multiple different tracts, resulting in ROIs that do not completely represent complete white matter pathways (**Supplemental Figure SE.F3**). In our approach, even though TractSeg predefines 72 white matter tracts, the approach is tractography-based, allowing us to study subject-specific white matter pathways derived directly from an individuals' dMRI data. This is an especially important aspect of our methodology, as white matter pathways are known to overlap in space[70,71], and tractography-based methods allow assignment of a single voxel to multiple different tracts. In this regard, we are the first to use WM tracts/pathways for constructing WM brain charts, as we are not using the atlas-based methods used in other research papers.

**Supplemental Table SE.T3.** Comparison of our WM brain charts to existing microstructural brain charts in the literature.

| **Study** | **Age Range** | **Datasets** | **Sample Size for Brain Charts** | **Total Sample Size (including validation)** | **Preprocessing** | **Normative modeling** | **Features {total number}** |
|---|---|---|---|---|---|---|---|
| Zhu et al. 2025 [47] | 3-95 | 10 | 13,297 | 40,898 | EPI- and eddy-current distortion correction; TBSS with JHU labels[69] | GAM + ComBat variations | FA {25} |

| | | | | | | | |
|---|---|---|---|---|---|---|---|
| Villalón-Reina et al. (preprint) [72] | 4-91 | 19 | 54,583 | 54,583 | EPI- and eddy-current distortion correction; TBSS with JHU labels[69] | HBR | FA, MD, AD, RD {88} |
| Our work | 0-100 | 50 | 35,120 | 41,897 | DWI Preprocessing (PreQual); DTI Fitting (MRtrix); Tractography (TractSeg); Whole-brain White Matter Segmentation (FreeSurfer) | GAMLSS | FA, MD, AD, RD, Volume, Surface Area, Average Length, Normalized Macrostructure {1157} |

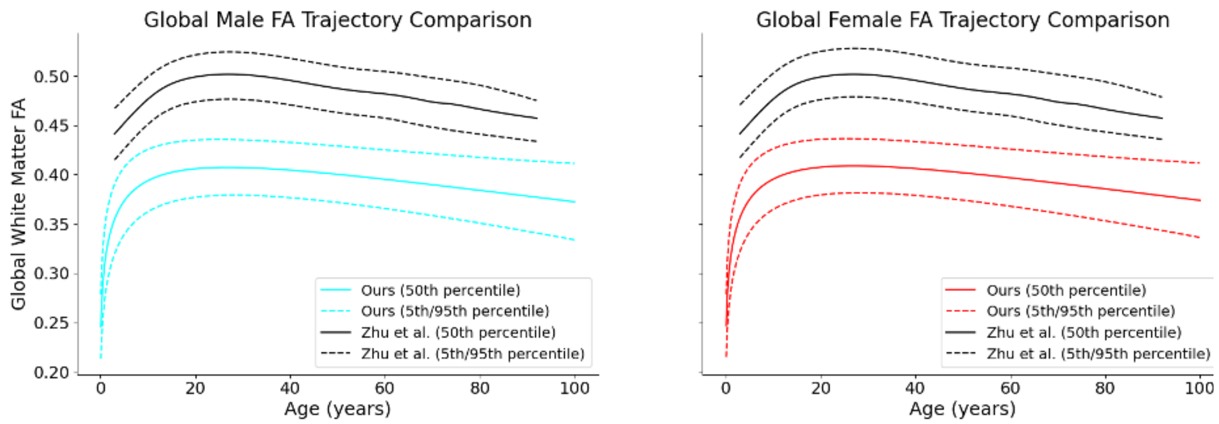


**Supplementary Figure SE.F2. Comparison of lifespan global FA trajectory to Zhu et al.** Our trajectory follows the same general trend as Zhu et al., with a sharp increase in adolescence, followed by a plateau in early adulthood, and then a decline in aging. Noticeably, the trajectories from Zhu et al. sit at higher FA values, due to the use of phenotype extraction along only the TBSS-defined white matter skeleton. (**Supplemental Figure SE.F4**).

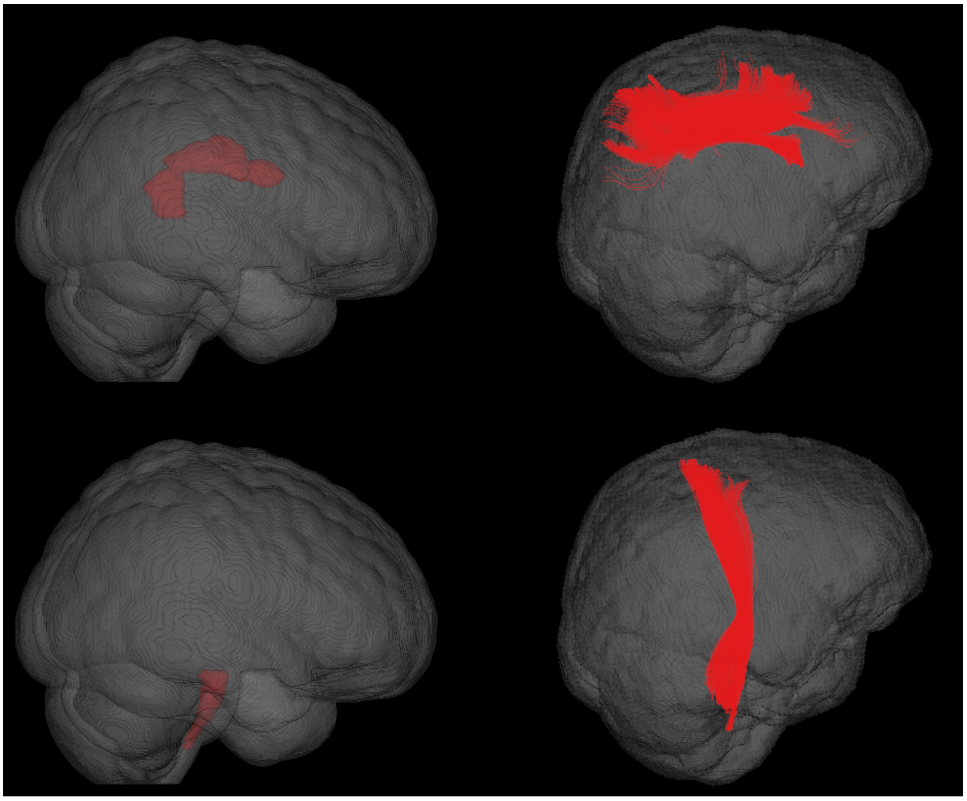

**Supplementary Figure SE.F3. Comparison to Zhu et al. for tract specific white matter coverage.** Zhu et al.'s method for tract-specific white matter coverage is based on the JHU atlas labels (left), whereas our method is based on subject-specific tractography using TractSeg (right). Notably, atlas-based approaches result in fundamentally spatially separated regions that do not constitute full pathways. Visualized are the left superior longitudinal fascicle (top) and the left corticospinal tract (bottom).

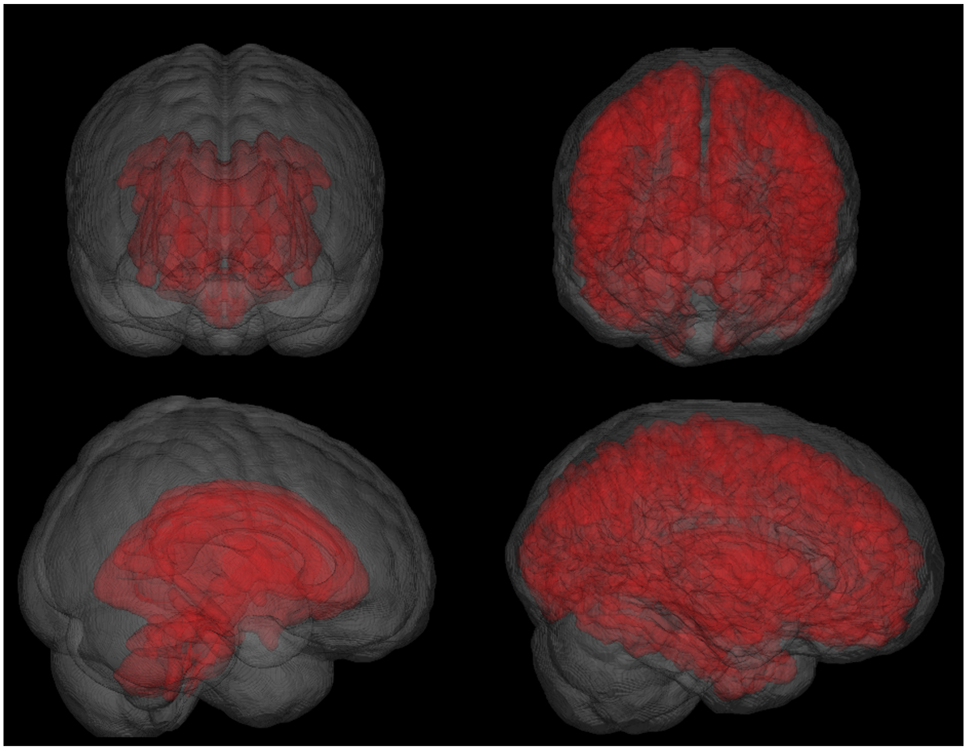

**Supplementary Figure SE.F4. Comparison to Zhu et al. white matter coverage for global microstructure.** Zhu et al.'s method utilizes the combined JHU atlas labels (left) to define white matter coverage for assessing global microstructure. In contrast, our method uses the general white matter mask derived from FreeSurfer (right). The FreeSurfer mask provides broader overall white matter coverage compared to the JHU atlas, which encompasses only specific white matter tracts. Furthermore, the FreeSurfer-based segmentation captures greater subject-specific anatomical variation by not relying on the boundaries of an atlas generated from a small cohort of young adults. The JHU labels are visualized with respect to the corresponding atlas, whereas the FreeSurfer labels are visualized from a particular individual from our database.

### F. **Data Quality**

As mentioned in the main manuscript, we conducted a thorough quality control process that involved visualizing each scan and performing quantitative outlier removal (see Methods: Data Selection). The final results of this process can be found in **Supplemental Table SB.T1**. Notably, infant datasets tend to have many more scans that are not included in the final participant pool for creating the brain charts. We attribute this effect largely to TractSeg, which was trained on an adult population. As infant data are more out-of-distribution compared to other datasets that the TractSeg models were not trained on, many infant scans showed poor tract reconstruction.

Furthermore, we conducted an analysis to assess both image quality across datasets and how image quality affects downstream centile score estimates. Specifically, we visualized distributions for the PreQual image quality metrics (IQMs) of mean relative displacement, the chi-squared error metric in shells used for diffusion tensor fitting, contrast-to-noise ratio (CNR) as calculated by EDDY for the main shell used for tensor fitting for scans included in creating the brain charts (**Supplemental Figure SF.F1**). We do not observe IQMs to be influenced by age at time of scan, as distributions are not consistently higher or lower at any given age range. In other words, IQM distribution appeared to be dataset-dependent rather than age-dependent.

For assessing correlation of estimated centile scores with IQMs, we computed Spearman's correlation coefficients between centile scores and each IQM (**Supplemental Figure SF.F2**). All correlations were either low or non-existent, with nearly all correlations is below 0.15 and none above 0.19. These results suggest that image quality of the dMRI data does not largely affect the estimated centile scores.

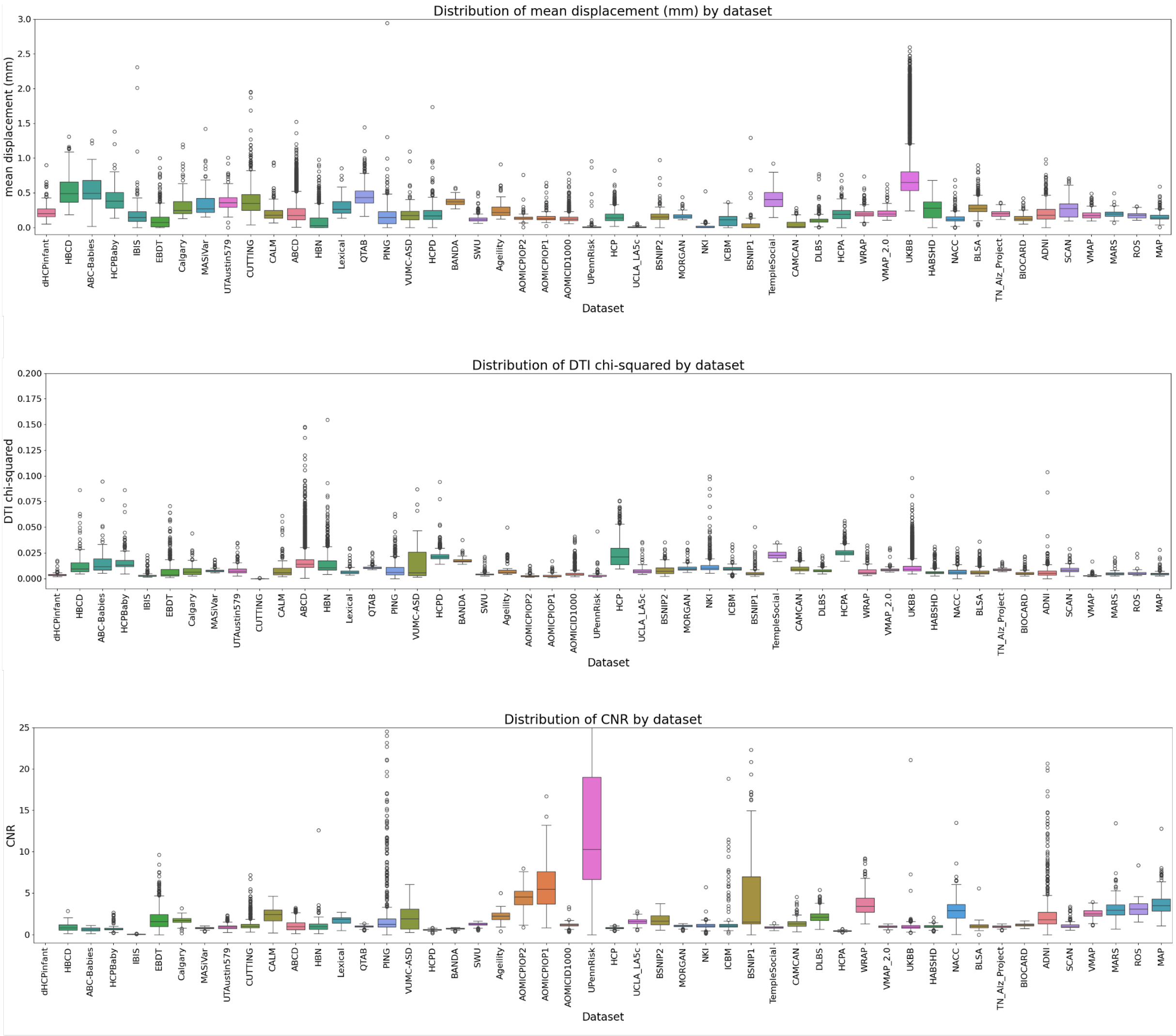


**Supplementary Figure SF.F1. Distribution of PreQual image quality metrics by dataset.** We do not observe age at time of scans to largely influence image quality metrics (IQMs) for dMRI preprocessing, as IQM distributions are not consistently higher or lower at any age range. Plotted are (top) motion as mean relative displacement, (middle) chi-squared error metric in the shells used for the diffusion tensor imaging fit, and (bottom) contrast to noise ratio (CNR) as calculated by eddy for the main shell used for the tensor fitting. Datasets are ordered by mean age of participants, with the youngest ages on the left and oldest on the right.

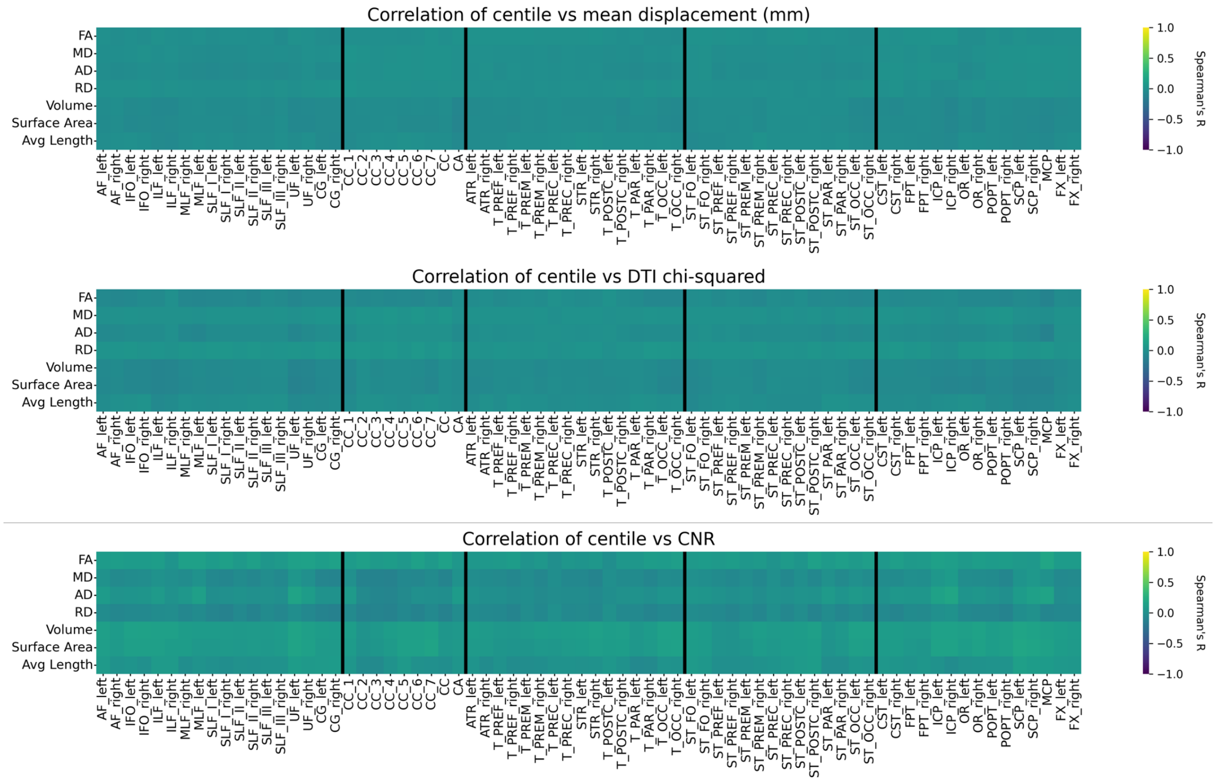


**Supplementary Figure SF.F2. Correlation of PreQual image quality metrics with centile scores.** We do not observe image quality metrics (IQMs) for dMRI preprocessing to be correlated with observed centile scores, as the absolute value of nearly all correlations is below 0.15, with none above 0.19. Plotted are Spearman's correlation coefficients with (top) motion as mean relative displacement, (middle) chi-squared error metric in the shells used for the diffusion tensor imaging fit, and (bottom) contrast to noise ratio (CNR) as calculated by eddy for the main shell used for the tensor fitting.

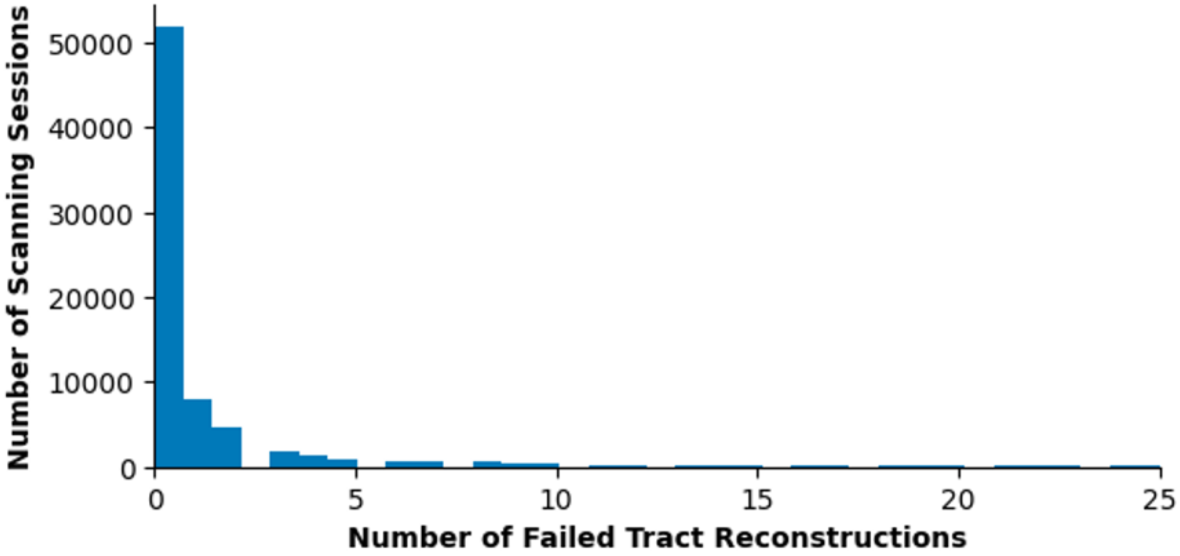


**Supplementary Figure SF.F3. Tract reconstruction failures per session.** While the majority of scanning sessions reconstructed most of the 72 TractSeg tracts, there were several scanning sessions that failed tract reconstruction for a large portion of tracts.

### G. **Anomaly Detection**

To assess the utility of our brain charts for performing predictive analyses on clinical cohorts, we performed a classification task to distinguish participants with Alzheimer's disease and mild cognitive impairment (MCI) from typically developing and aging individuals. For the classification, we used centile scores for the microstructural features of FA, MD, AD, and RD and for the macrostructural features of volume, surface area, and average length for all 72 tracts (504 total centile score features). A multilayer perceptron (MLP) with two hidden layers was trained on the 504-dimensional input, using ReLU activation between layers, class-weighted Cross Entropy as a loss function, and the Adam optimizer with a learning rate of 0.001. The network was trained with 5-fold cross validation, with classes evenly split across folds. For individuals whose scans did not reconstruct all 72 tracts in the processing pipeline, the *miceforest* algorithm was used for data imputation (https://github.com/AnotherSamWilson/miceforest). We report area under the receiver operating characteristic curve (ROC-AUC) and balanced accuracy in **Suppplemental Tables SG.T1 and SG.T2**. The 0.910 +/- 0.910 and 0.829 +/- 0.015 average AUCs across folds indicate promising discriminative performance; however, we still advocate that the utility of these brain charts should be as research tools rather than clinical diagnostic tools.

**Supplemental Table SG.T1. Results of Alzheimer's disease classification experiment.**

| Fold | Balanced Accuracy | ROC-AUC |
|---|---|---|
| **Fold 0** | 0.810 | 0.888 |
| **Fold 1** | 0.807 | 0.901 |
| **Fold 2** | 0.820 | 0.907 |
| **Fold 3** | 0.866 | 0.937 |
| **Fold 4** | 0.789 | 0.916 |
| **Average** | **0.819 +/- 0.029** | **0.910 +/- 0.018** |

**Supplemental Table SG.T2. Results of mild cognitive impairment classification experiment.**

| Fold | Balanced Accuracy | ROC-AUC |
|---|---|---|
| **Fold 0** | 0.746 | 0.851 |
| **Fold 1** | 0.721 | 0.818 |
| **Fold 2** | 0.721 | 0.812 |
| **Fold 3** | 0.737 | 0.833 |
| **Fold 4** | 0.712 | 0.832 |
| **Average** | **0.728 +/- 0.014** | **0.829 +/- 0.015** |

### H. **Out-of-Sample Alignment**

#### i. Stability of Out-of-Sample Alignment

To assess the stability of the out-of-sample (OOS) alignment process, we performed OOS alignment for the HCPA dataset at varying sample sizes. At each sample size (from 700 to 50 in increments of 50), we randomly selected participants from the dataset without replacement, with 100 different bootstrapped assessments at each sample size. We did this for both FA and volume of the right cingulum and isthmus of the corpus callosum. Across all features, we observed that the OOS alignment process is more stable when estimating the $\mu$ random effect term compared to the $\sigma$ random effect term (**Supplemental Figure SH.F1**). Although stability of estimates vary across the tracts, features, and parameters, we observe a conservative threshold for stability in OOS alignment at N=100. This threshold is similar to that from Bethlehem et al.[48], suggesting that our method is in agreement with existing normative brain charts.

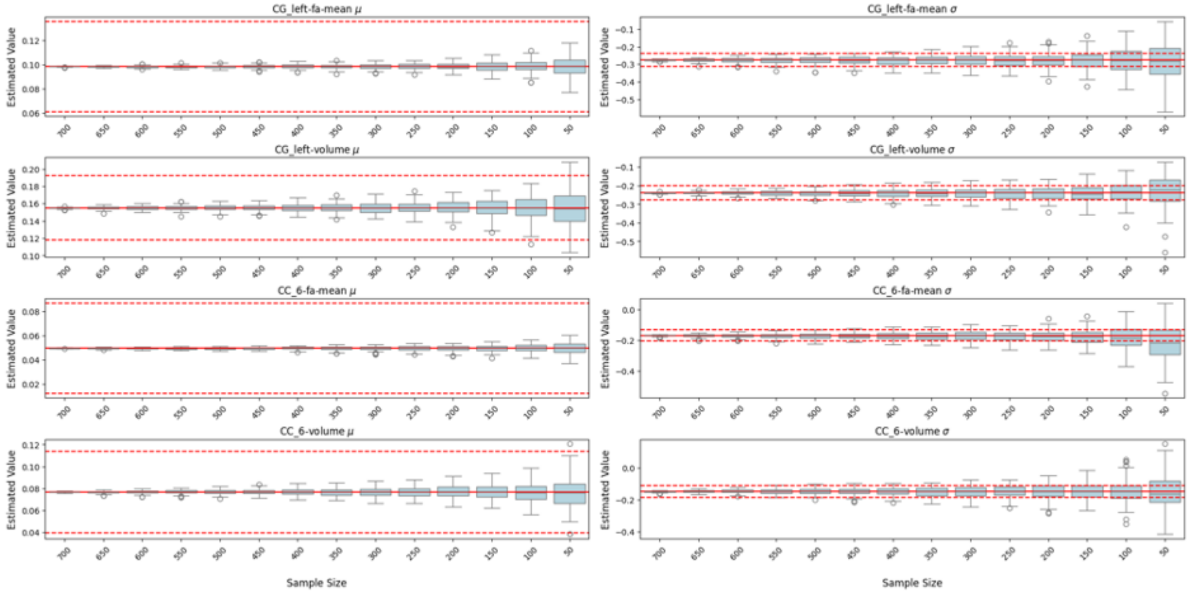


**Supplementary Figure SH.F1. Out-of-sample alignment bootstrapping experiment for determining minimum sample size.** Out-of-sample (OOS) alignment appears stable and within the GAMLSS-estimated standard error (solid red lines indicate the GAMLSS-estimated value, dashed red lines indicate error bars) for sample sizes greater than or equal to 100. The estimated random effect term for $\mu$ (left) appears more stable for OOS alignment than it is for $\sigma$ (right), as the distributions for bootstrapped random effects fall within the GAMLSS error range at smaller sample sizes for $\mu$. We observe varying levels of stability for different features and tracts (from top to bottom: FA of right cingulum, volume of right cingulum, FA of isthmus of corpus callosum, volume of isthmus of corpus callosum), with N=100 as a conservative threshold.

We note that, as part of our postprocessing pipeline, constrained spherical deconvolution (CSD) is run to extract peak directions for downstream tractography. Although not evaluated in this manuscript, it is generally accepted that CSD requires roughly 30 directions for accurate estimation; however, CSD can also be performed with fewer directions, albeit with less stable estimates.

#### ii. Alignment with Other Preprocessing Pipelines

The DWI preprocessing pipeline that we use, PreQual, is generically state-of-the-art, as it contains all the fundamental steps that are considered "essential" (corrections for susceptibility-distortions, motion, eddy-current-distortions)[73]. These fundamental steps are present in most dMRI preprocessing workflows. Although we used PreQual to preprocess our DWI data, we acknowledge that alternative tools for DWI preprocessing exist, such as DTIPrep/Dmriprep[74,75] DSI-Studio[76], ExploreDTI[77], and TORTOISE[78]. We also include additional steps that are considered beneficial for data quality (denoising; slice-wise outlier imputation). Thus, researchers can generically use different preprocessing pipelines.

To assess whether or not data corrected with different DWI preprocessing pipelines are alignable to our brain charts, in an extreme case, we performed out-of-sample alignment using data from the AOMICPIOP2 dataset[4] that had not had any DWI preprocessing. We then compared the original centile scores for each participant to those obtained via the raw data alignment. Most features are highly correlated (**Supplemental Figure SH.F2**), suggesting that even in this extreme case, data are able to be aligned to our brain charts without having run data through the PreQual preprocessing pipeline. However, we do note that there are a few features, notably smaller tracts like the fornix (FX) or anterior commissure (CA), that are not as highly correlated. Regarding specific tract groups, we observed that cerebellar tracts (MCP, ICP, SCP) and projection tracts (FPT, CST) also exhibited lower correlations than other tract groups, likely due to their increased sensitivity to distortion artifacts in these brain regions.[79] We did not observe any features to be more or less correlated across tracts when compared to other features.

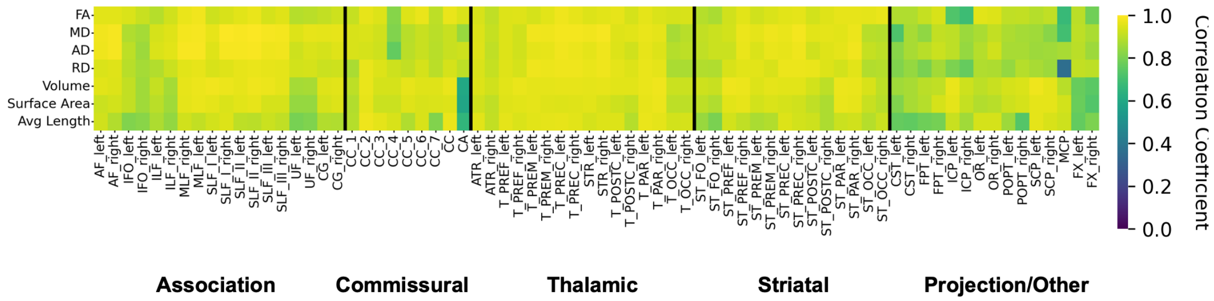


**Supplementary Figure SH.F2. Pearson correlation of centile scores for the AOMICPIOP2 dataset with and without dMRI preprocessing.** Centile scores from both preprocessed and raw data are highly correlated across most tracts and features; this extreme case indicates that DWI data that have undergone different preprocessing workflows are can be appropriately aligned to our brain charts. Measurements from smaller tracts (FX, CA) and cerebellar/projection tracts are most affected by lack of DWI preprocessing, showing reduced correlations when compared to other tracts.

#### iii. How to Perform Out-of-Sample Data Alignment

One of the most important aspects of brain charts is the ability to score new data within the normative trajectories to determine how abnormal quantitative brain metrics are. We have demonstrated how we perform alignment of new datasets to the centile curves (see Methods: Maximum Likelihood Estimation for Out-of-Sample Datasets). For any researchers who would like to use these brain charts, we release a Docker container as an easily reproducible method of performing this alignment with new data and provide the following tutorial:

1.) Before doing anything, Docker needs to be installed. Official instructions can be found here: (https://docs.docker.com/get-started/get-docker/). [80] Docker must be running properly before proceeding.

2.) Download the Docker image (as a .tar file) from Zenodo at this link: https://zenodo.org/records/17561821

Save the .tar file somewhere it can be easily found later, such as a Downloads folder.

3.) Open a terminal (command line tool). Load the Docker image, using the following command-line prompt:

```
docker load -i </path/to/docker/.tar/file>
```

The Docker image can also be loaded through the Docker Desktop GUI.

4.) Confirm that the Docker image is loaded by running:

```
docker images
```

5.) Before alignment, the data must be properly formatted in a CSV file that can be read by the Docker image. The CSV is required to have columns $age$, $sex$, and $diagnosis$, where $age \geq 0$ is a numerical value, $sex$ is a binary variable where "male" is encoded as $1$ and "female" is encoded as $0$, and $diagnosis$ is a categorical variable. Typically developing/aging (also referred to as "cognitively normal") participants are encoded as "CN" for $diagnosis$, and to perform alignment there must be rows in the CSV file that contain "CN" as the $diagnosis$. For better alignment, ensure as many "CN" participants as possible, and note that having a small number of participants may result in poorly aligned data and thus poorly estimated centile scores. There must also be at least one quantitative variable column in the CSV file, where quantitative variables are named as:

```
<tract>-<measure>
```

`<tract>` must be one of the TractSeg defined tract names found in Supplementary Table S4, whereas `<measure>` must be one of `{fa-mean, md-mean, ad-mean, rd-mean, volume, surface_area, avg_length}`. Thus, the CSV should follow formatting such as:

| age | sex | diagnosis | AF_left-fa-mean | AF_right-md-mean | … |
|---|---|---|---|---|---|
| 75.1 | 0 | CN | 0.453 | 0.00110 | … |
| 45 | 1 | CN | 0.562 | 0.00140 | … |
| 62.5 | 1 | AD | 0.398 | 0.00098 | … |
| … | … | … | … | … | … |

Note that in cases where rows have empty entries, centile scores will not be calculated for these metrics and as a result will have a missing entry in the respective centile score output. Further, only rows labeled with a "CN" $diagnosis$

and with non-missing centile values will be used for estimating the random effect terms for a particular measure.

6.) Run the Docker using the following command:

```
docker run --rm -v </path/to/OOS.csv>:/INPUTS/input.csv \

-v </path/to/output/directory>:/OUTPUTS \

r_lifespan_env \

python3 /WMLifespan/scripts/perform_OOS_alignment.py \

/INPUTS/input.csv /OUTPUTS/aligned.csv
```

where `aligned.csv` is the destination file you wish to save the aligned centile score values.

As detailed in the Methods section, these normative curves are cross-sectional in nature. Thus, researchers performing out-of-sample alignment should only include cross-sectional data in the CSV file, or one scan per participant. Should researchers wish to evaluate longitudinal data with the cross-sectional models, the flag can be used to also save the estimated random effect terms for the dataset. We also note that this alignment to the normative models assumes that the data in the CSV file come from the same primary dataset. Calculation of centile scores for multiple datasets need to be done in separate Docker commands, each with their own distinct input CSV file.

#### iv. How to obtain centile trajectories of features

For researchers who wish to examine the normative trajectories of features more closely, we also provide a method for obtaining centiles of trajectories in a CSV format:

1.) To begin, follow steps 1.) through 4.) of the section “How to Perform Out-of-Sample Data Alignment” in order to properly set up the Docker image.

2.) To obtain centile curves for a given tract and metric, run the following command:

```
docker run --rm \

-v </path/to/output/directory>:/OUTPUTS \

r_lifespan_env \

python3 /WMLifespan/scripts/output_centile_curves.py \

<tract> <measure> /OUTPUTS/centiles.csv
```

where `centiles.csv` is the destination file you wish to save discrete values of the normative trajectory for the given `<tract>` and `<measure>`. Note that `<tract>` must be one of the TractSeg defined tract names found in Supplementary Table S4, whereas `<measure>` must be one of `{fa-mean, md-mean, ad-mean, rd-mean, volume, surface_area, avg_length}`.

### I. Considerations for Cross-sectional vs. Longitudinal Brain Charts

Previous work has shown that cross-sectional normative models can be inaccurate in predicting measured longitudinal trajectories[81], as longitudinal models can better separate true developmental change from generational differences that exist in the data and cross-sectional data may be unrepresentative of individuals. Although many of our datasets contained longitudinal data, there were several challenges we encountered and aspects about the normative models we needed to consider for longitudinal brain charts[82]. First, despite the subset of datasets with participant data at multiple time points, for the majority of participants in the study we only have cross-sectional data available. Incorporating longitudinal data would require estimation of random effects for each participant, introducing significant computational complexity in the model. We found that we did not have enough data points to stably estimate individual-level random effects. Furthermore, having to estimate these individual-level random effects would greatly decrease the stability and increase complexity of the out-of-sample centile scoring process, as alignment with $N$ participants would require estimation of $2N$ terms instead of only 2 terms with the cross-sectional models. These constraints and considerations led us to develop a robust cross-sectional approach that maintains generalizability and computational feasibility. We acknowledge the potential benefits of longitudinal modeling and suggest this as an important avenue for future research.
